# Engineering compact bacterial combinatorial promoters for two-input chemical AND switching

**DOI:** 10.64898/2026.03.25.714203

**Authors:** Satya Prakash, Alfonso Jaramillo

## Abstract

Engineering compact bacterial promoters that integrate two chemical inputs would simplify biosensors, synthetic circuits, and dynamic metabolic control, but such promoters remain difficult to build because a second operator can increase partial-state leak, reduce the fully induced state, and create sequence- or reporter-context effects. Here we engineered 12 compact Marionette-derived two-input promoter architectures in *Escherichia coli* by combining defined template scaffolds with added operators, then quantified 19 reporter-specific four-state truth tables under a predefined operational criterion for chemical-input AND behaviour. Single-input template-promoter controls in matched reporter contexts helped separate weak parent-scaffold output from effects introduced by the second operator. Nine curated architectures passed in at least one reporter, and selected constructs delivered high-utility AND responses, including [11]/max-off separation up to >1,367. The full pass–fail dataset showed that a statistically higher [11] state does not necessarily create a useful switch. Practical utility depended mainly on suppressing [10] and [01] leakage, matching the template scaffold to the inserted operator, and controlling long-operator orientation and local sequence context. Reciprocal architectures behaved differently, and 4 of 7 dual-reporter architectures changed operational classification with reporter context. Thus, this work delivers a compact set of candidate Marionette-compatible two-input promoter parts and a source-traceable design framework for engineering bacterial chemical-input AND switches.

## Introduction

Synthetic biology relies heavily on promoters that can integrate multiple signals to control cellular circuits, biosensors, and dynamic metabolic pathways^1^. In bacteria, such behaviour is often implemented by layering single-input regulators across several genes, but compact combinatorial promoters remain attractive because they reduce circuit size, shorten response paths, and provide direct experimental access to how local sequence architecture shapes transcription^2–7^. Previous studies have already shown that combinatorial promoter performance depends on context, operator arrangement, and promoter geometry^2–11^. The practical challenge is that these dependencies still make compact two-input promoters difficult to design prospectively.

Adding a second operator to a bacterial promoter can elevate partially induced states, reduce the effective separation of the fully induced state, or abolish repression altogether. Reciprocal architectures rarely behave equivalently, and reporter context can change the apparent logic outcome even when the same promoter sequence is retained^9–11^. Long operators further complicate the design space because they can introduce promoter-like motifs or perturb the early transcribed region. These issues are familiar in principle, but they are often discussed from a small number of successful end-point constructs rather than from mixed datasets that retain failures alongside passes.

We therefore asked not only which architectures satisfy an operational AND criterion, but which architectural features distinguish success from failure within a controlled two-input dataset. Rather than exhaustively sampling every possible promoter-operator pairing, we assembled a comparative panel that spans different scaffolds, added operators, operator lengths, orientations, downstream positions, and reciprocal arrangements. This framing treats non-passing architectures as informative negative results rather than as discarded intermediate attempts.

To minimise upstream sensor cross-talk while focusing the analysis on promoter architecture, we built the study around the Marionette *Escherichia coli* platform^13^. This background provides a shared set of chromosomally encoded, small-molecule-responsive repressors and therefore lets the plasmid-borne output promoters be compared in a common regulatory context. Because several Marionette modules contain evolved components^13^, we treat the present study as a benchmark of compact promoter design within an optimised sensor framework rather than as an exhaustive survey of all possible wild-type regulator combinations.

Previous work on Marionette primarily optimised ligand specificity, dynamic range, and cross-talk^13^. Here we use that platform to benchmark which compact two-input promoter architectures remain functional once a second operator is introduced into or near the early transcribed region. The resulting dataset therefore complements prior mechanistic and engineering studies by making scaffold compatibility, leakage, reciprocal asymmetry, and reporter dependence directly comparable within a single experimental framework.

Throughout this study, truth-table language is operational in inducer space. Because the chemical inputs relieve repression, the chemical [11] state corresponds to both repressor branches being inactive; in that sense, a chemical-input AND response is implemented through a transcription-factor-level NOR-like transfer function. We therefore use inducer-conditioned truth tables as an engineering benchmark, while avoiding claims of mechanistic promoter-level AND cooperativity that are not resolved by the present fluorescence data alone. The central claim of this benchmark is that compact Marionette two-input promoters must be evaluated as complete promoter-gene contexts rather than as modular sums of two single-input elements; the mixed pass-fail dataset provides usable parts and empirical constraints within that defined scope.

## Results

### Library design, reporter selection, and comparative analysis framework

We analysed 12 prespecified curated architectures and one exploratory pTac-Van construct. Figure 1a summarises the design-to-analysis workflow, while Figure 1b and Table 1 summarise the curated architecture set. We designed each architecture from a template Marionette promoter and an added operator, tested it in the relevant reporter context, selected complete non-toxic four-state grids, fitted fluorescence-versus-OD slopes, applied a predefined operational pass/fail/N.C. classification, and assessed utility metrics separately. We tested each architecture as EGFP, mCherry, or both when data were available. We treated GFP and RFP as independent output reporters rather than normalisation controls.

**Fig. 1.**
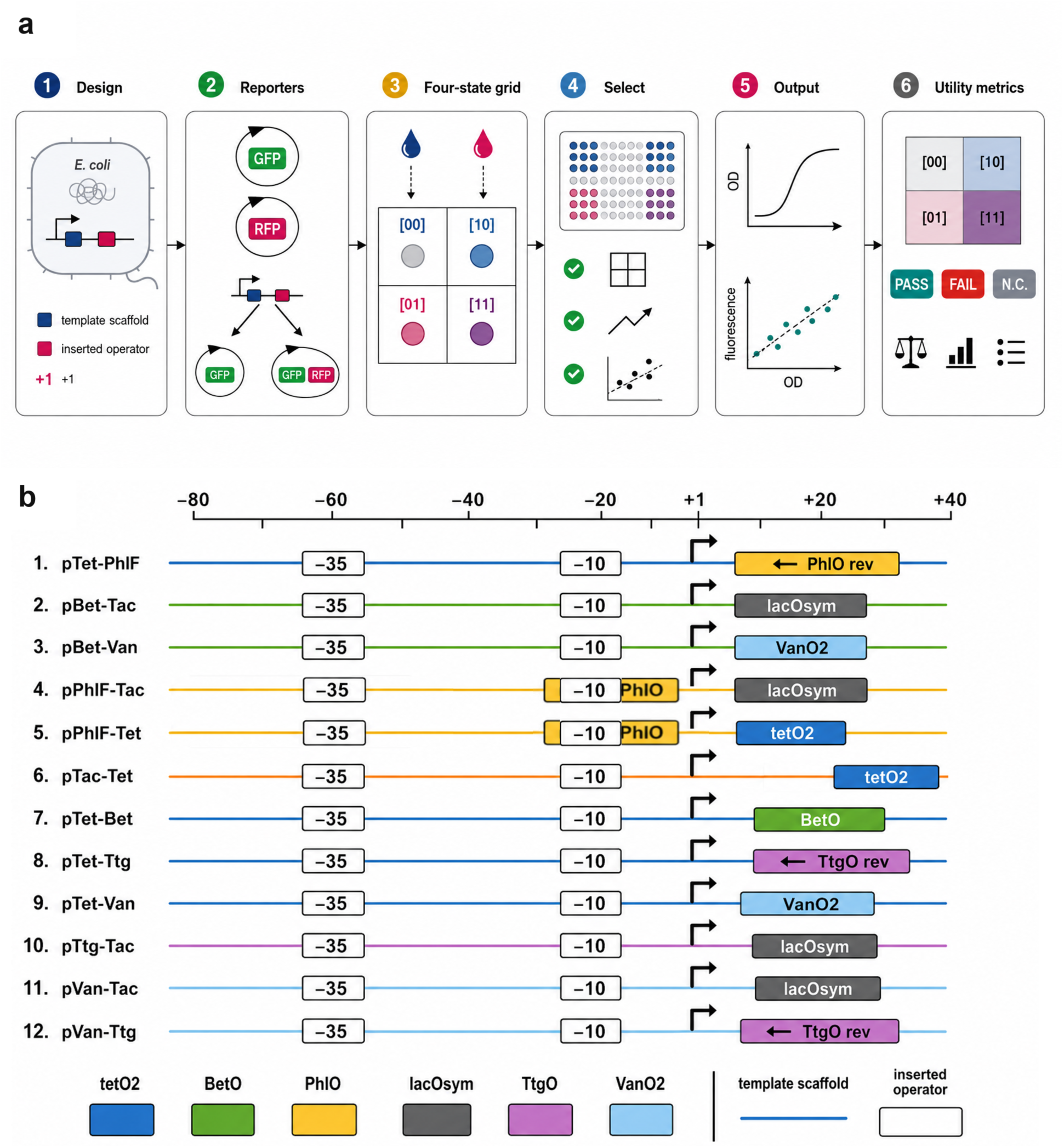
Experimental workflow and architectures of the curated engineered two-input promoter set. **a**, Workflow schematic for the analysis pipeline. The workflow runs from architecture design to reporter-context testing, complete non-toxic four-state grid selection, fluorescence-versus-OD slope fitting, predefined operational pass/fail/N.C. classification, and separate utility metrics. GFP and RFP were independent output reporters rather than normalisation controls. For each truth table, the first digit denotes the template-promoter input and the second digit denotes the added-operator input. pTac-Van was exploratory/non-curated and is documented separately in Supplementary Figure S17 and the Supplementary Table S7 annotation table. **b,** Architecture map for the 12 curated engineered two-input promoters. Constructs are aligned relative to the transcription start site (+1, black bent arrow). Coloured lines indicate the template scaffold, small boxed labels mark the annotated -35 and -10 elements, and larger coloured boxes indicate operator sequences. The pPhlF-derived architectures explicitly show the native PhlO motif carried by the scaffold, and the long PhlO and TtgO insertions used downstream in pTet-PhlF, pTet-Ttg, and pVan-Ttg are shown in the reverse orientation adopted in the final designs. Supplementary Figure S19 highlights representative sequence context around the -10 and +1 regions, and Supplementary Table S7 provides the corresponding annotated sequences.

**Table 1.**
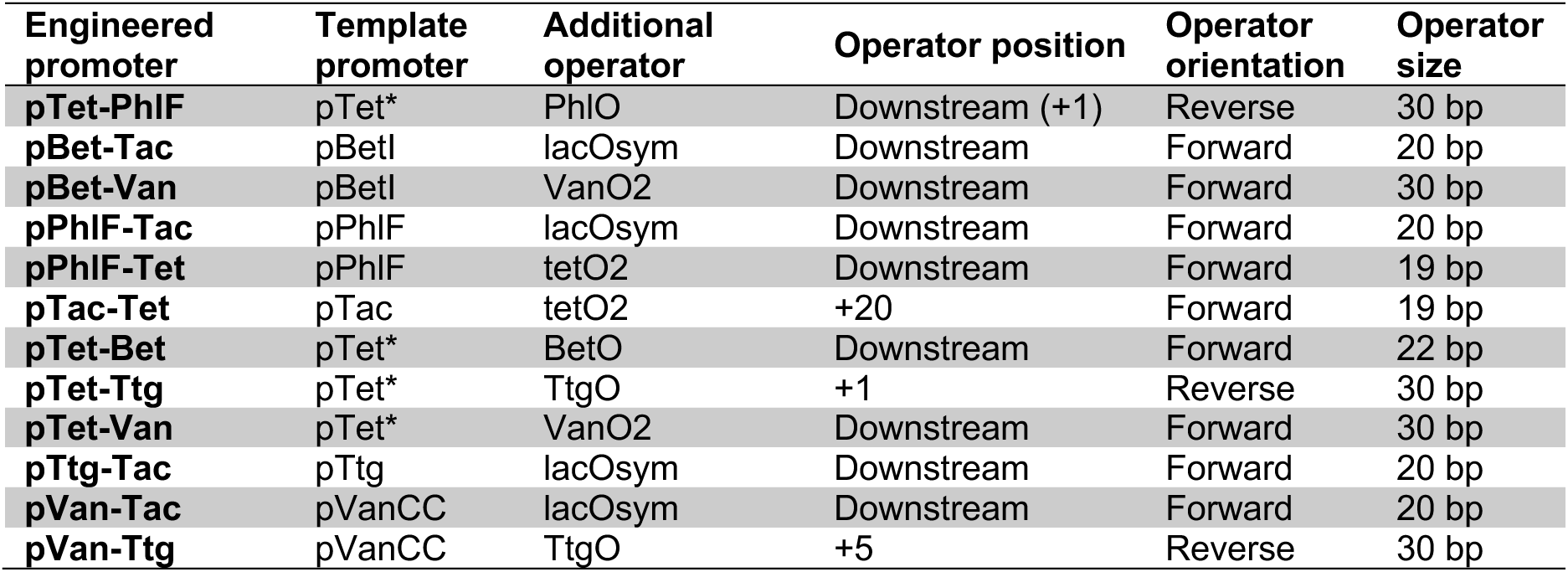
Architecture of the curated two-input promoter set. Template promoter names follow the original Marionette nomenclature13. pVanCC and pTet* are retained where applicable. The table summarises the structural variables used in the comparative analysis, whereas the exploratory pTac-Van construct is shown separately in Supplementary Figure S17 and Supplementary Table S7.

| Engineered promoter | Template promoter | Additional operator | Operator position | Operator orientation | Operator size |
| --- | --- | --- | --- | --- | --- |
| pTet-PhIF | pTet* | PhIO | Downstream (+1) | Reverse | 30 bp |
| pBet-Tac | pBetI | lacOsym | Downstream | Forward | 20 bp |
| pBet-Van | pBetI | VanO2 | Downstream | Forward | 30 bp |
| pPhIF-Tac | pPhIF | lacOsym | Downstream | Forward | 20 bp |
| pPhIF-Tet | pPhIF | tetO2 | Downstream | Forward | 19 bp |
| pTac-Tet | pTac | tetO2 | +20 | Forward | 19 bp |
| pTet-Bet | pTet* | BetO | Downstream | Forward | 22 bp |
| pTet-Ttg | pTet* | TtgO | +1 | Reverse | 30 bp |
| pTet-Van | pTet* | VanO2 | Downstream | Forward | 30 bp |
| pTtg-Tac | pTtg | lacOsym | Downstream | Forward | 20 bp |
| pVan-Tac | pVanCC | lacOsym | Downstream | Forward | 20 bp |
| pVan-Ttg | pVanCC | TtgO | +5 | Reverse | 30 bp |

For each AND truth table, the first digit denotes the template-promoter input and the second digit denotes the added-operator input. Figure 2 and Supplementary Table S2 summarise the full set of tested promoter-reporter truth tables. For each promoter-reporter combination, we selected the cleanest complete non-toxic grid, defined as a four-state [00]/[10]/[01]/[11] inducer set with acceptable growth and a valid fluorescence-versus-OD fit in every state. We then chose the highest non-zero inducer pair that preserved such a complete grid. This procedure yielded 19 curated promoter-reporter truth tables across the 12 comparative architectures.

**Fig. 2.**
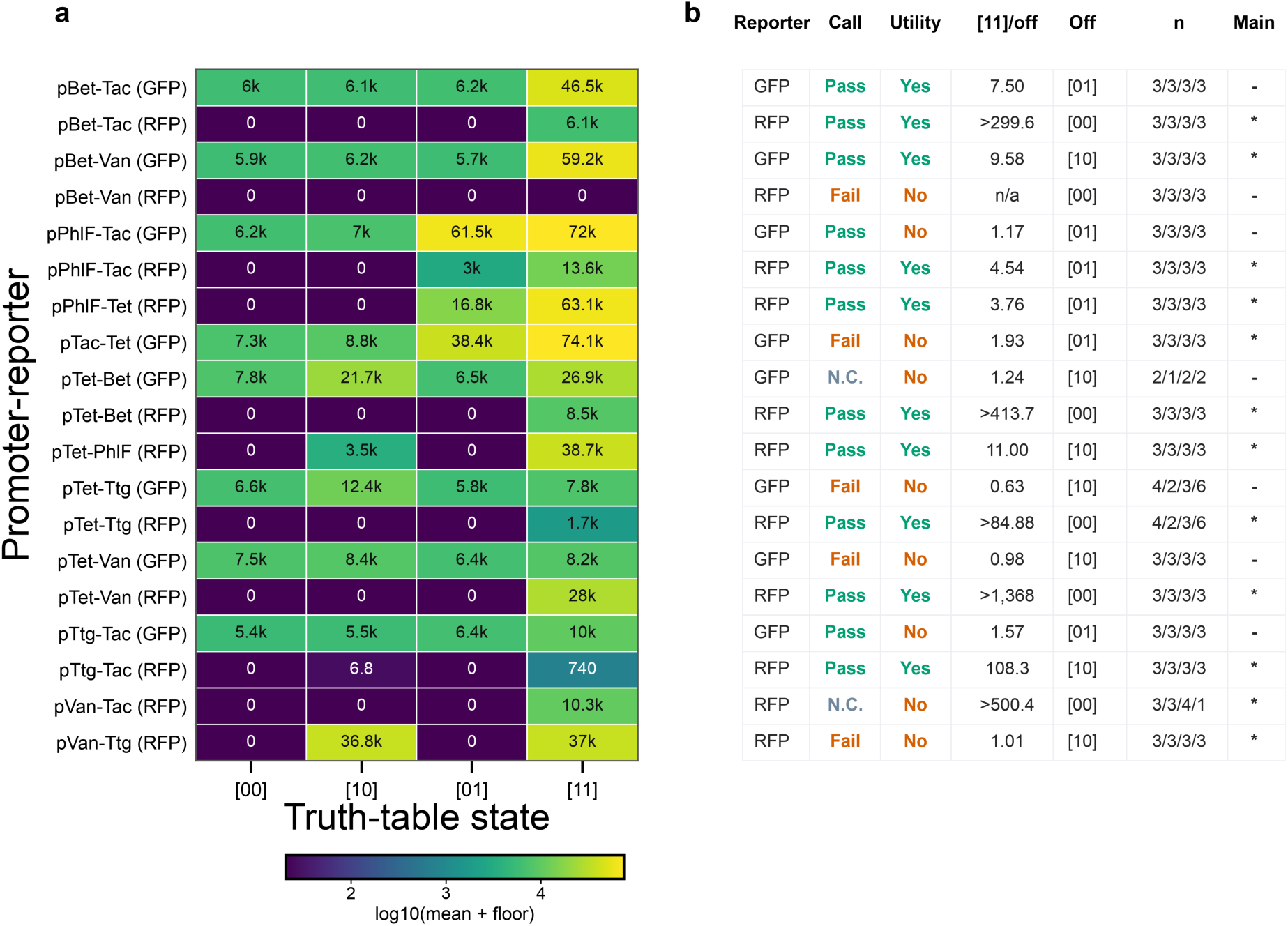
Dataset-wide summary of all curated promoter-reporter truth tables. **a**, Heat map of mean normalised fluorescence-versus-OD slopes for every curated promoter-reporter combination analysed in this study. The colour scale encodes log10(mean + empirical display floor), using the response floor described in Methods only for display and lower-bound ratios. **b,** Source-traceable annotation table reporting reporter identity, operational classification, Engineering Utility flag, [11]/max-off ratio, highest off state, biological replicate counts in [00], [10], [01], and [11] order, and the reporter row used as the architecture-level representative. Engineering Utility is True only for rows that pass the predefined operational AND criterion and have [11]/max-off at least 2.0. Source values are reported in Supplementary Table S2. Lower-bound ratios are marked with > where the maximum off-state mean was clipped to 0.00 a.u./OD. N.C. indicates not classifiable because one or more required state comparisons lacked sufficient biological-replicate support. For each promoter, the first digit corresponds to the template-promoter input and the second digit corresponds to the added-operator input. Inducer identities are listed in Supplementary Table S6. The two rows containing an n = 1 state, pTet-Bet(GFP) and pVan-Tac(RFP), are retained for transparency and are N.C.; neither supports a high-utility gate call.

We experimentally tested the exploratory pTac-Van construct and retained it in the Supplementary Information. However, we excluded it from the curated benchmark because it did not provide a comparably clean complete non-toxic four-state dataset. We used the Holm-adjusted one-sided Welch screen as a predefined operational criterion, but analysed engineering utility separately from that binary pass/fail call.

Several architectures produced clear high-utility chemical-input AND responses Figure 2 summarises all 19 curated promoter-reporter truth tables, including operational pass/fail calls, Engineering Utility flags, [11]/max-off values, highest off-state identity, selected-reporter markers, and biological replicate counts reported per state. Nine architecture-level representative rows passed the predefined operational screen and met the Engineering Utility threshold. The strongest selected rows were pTet-Van(RFP), pTet-Bet(RFP), and pBet-Tac(RFP), with [11]/max-off ratios of >1,367, >414, and >300, respectively, because all three off states were near the empirical response floor. Additional high-utility selected gates included pTtg-Tac(RFP), pTet-Ttg(RFP), pTet-PhlF(RFP), pBet-Van(GFP), pPhlF-Tac(RFP), and pPhlF-Tet(RFP), with [11]/max-off ratios from 3.76 to 108. Figure 4 shows these candidate Marionette-compatible two-input promoter parts as direct [00]/[10]/[01]/[11] truth tables with biological replicate points, per-state n values, inducer pairs, [11]/max-off ratios and highest off states.

### Leakage in the single-input states is the dominant engineering challenge

Quantitative comparisons across all reporter-specific truth tables emphasise that leakage suppression, not maximal [11] output, best separates strong and weak designs (Figure 3a,b). The same dataset also shows why statistical passing alone is insufficient. Several constructs generated large fully induced signals yet still performed poorly because [10] or [01] remained too high. pPhlF-Tac(GFP) is a statistical/operational pass but not a practical high-utility gate because the [01] state remains close to [11]. pTtg-Tac(GFP) similarly passed operationally but did not meet the Engineering Utility flag. In this dataset, engineering utility therefore tracks suppression of the highest off state more closely than absolute maximal brightness.

**Fig. 3.**
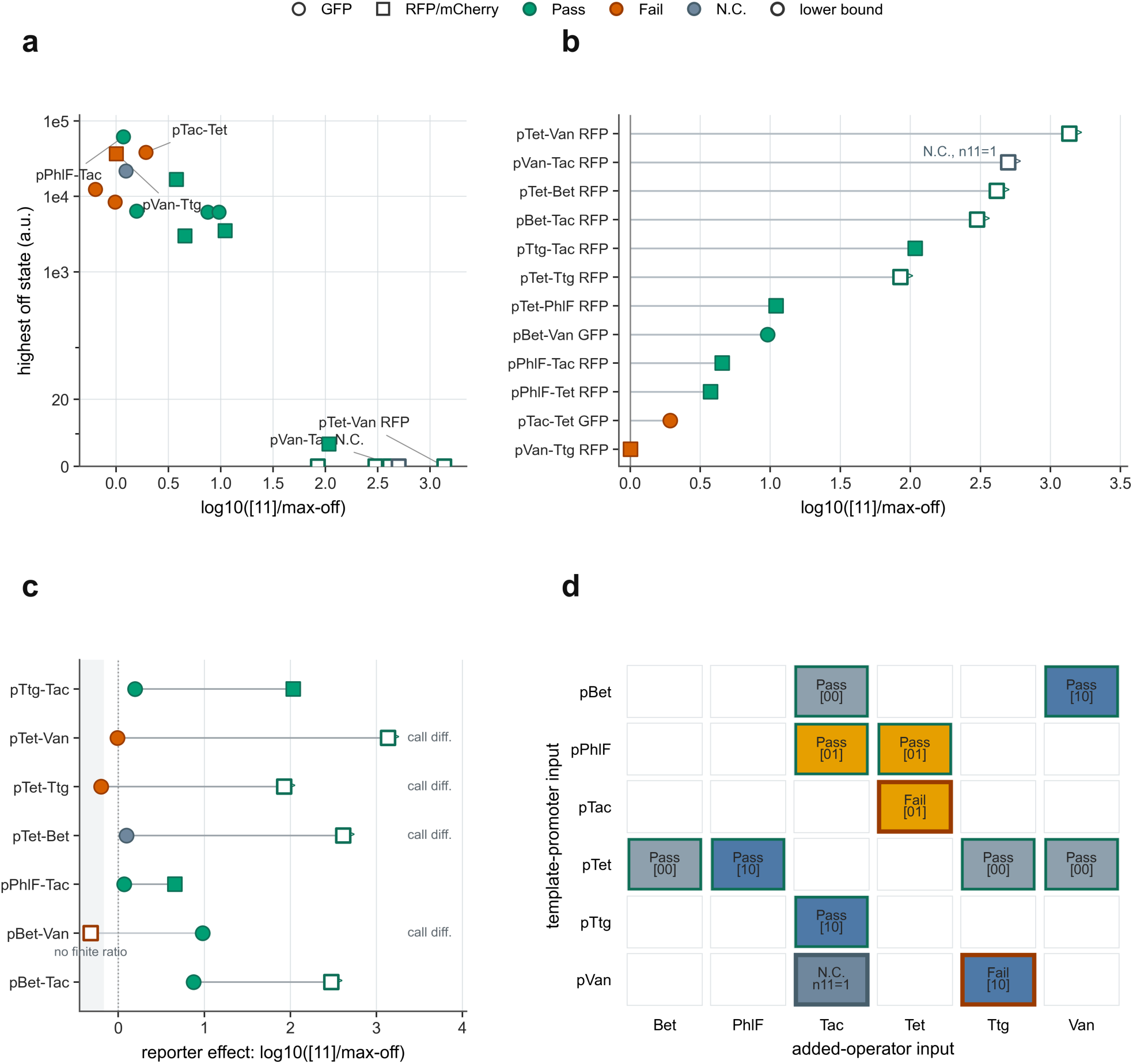
Quantitative constraints on AND-promoter utility across leakage, reporter and scaffold contexts. **a**, Leakage constraint map for reporter-specific truth tables in Supplementary Table S2. The x-axis shows log10([11]/max-off), where [11]/max-off is the fully induced mean divided by the brightest non-[11] state. The y-axis shows the brightest off-state mean. In panels a-**c,** circles denote GFP rows and squares denote RFP/mCherry rows. Green markers denote operational pass, red-orange markers denote fail, and grey-blue markers denote N.C. Open markers with > labels denote lower-bound ratios caused by max-off values clipped to 0.00 a.u./OD. **b,** Ranked log10([11]/max-off) for the reporter row used as the architecture-level representative for each promoter design. **c,** Dual-reporter architectures plotted as paired GFP and RFP log10([11]/max-off) values. Rows labelled call diff. have reporter-dependent operational calls, and pBet-Van RFP is labelled no finite ratio because [11]/max-off is undefined. **d,** Scaffold-by-added-input matrix for the reporter rows used as architecture-level representatives. The y-axis gives the template-promoter scaffold and the x-axis gives the added regulatory input. Each filled tile represents one tested architecture. Green tile borders mark pass, red-orange borders mark fail, and grey-blue borders mark N.C. Tile fill identifies the brightest off state for that architecture: grey-blue [00], blue [10], or orange [01]. Text inside each tile reports Pass, Fail, or N.C. and gives the limiting off state for failures. Together these panels separate operational classification from engineering utility and show that leakage, reporter context and scaffold/input direction affect practical outcomes.

### Reciprocal architectures and scaffold compatibility are not interchangeable

Reporter-normalised comparisons also revealed strong architectural asymmetry (Figure 3c,d). The reciprocal pair pTet-PhlF and pPhlF-Tet both passed, but their selected-reporter [11]/max-off ratios differed by about 2.9-fold (10.999 versus 3.756), showing that swapping scaffold and inserted input does not preserve performance. More broadly, all four selected pTet*-derived architectures passed the operational screen, as did both pBetI-derived and both pPhlF-derived designs, whereas neither pVanCC-derived architecture produced a classifiable pass: pVan-Tac was N.C. because only one [11] biological replicate passed QC, and pVan-Ttg failed because [10] rose to the [11] level. These counts are not intended as universal rankings, but within this Marionette benchmark they show that scaffold identity was a dominant variable rather than a neutral background. Figure 4 therefore focuses on high-utility selected gates, while weak, failed, and non-classifiable truth tables remain source-traceable in Figure 2, Figure 3, Supplementary Figs. S13-S16, and Supplementary Table S2.

**Fig. 4.**
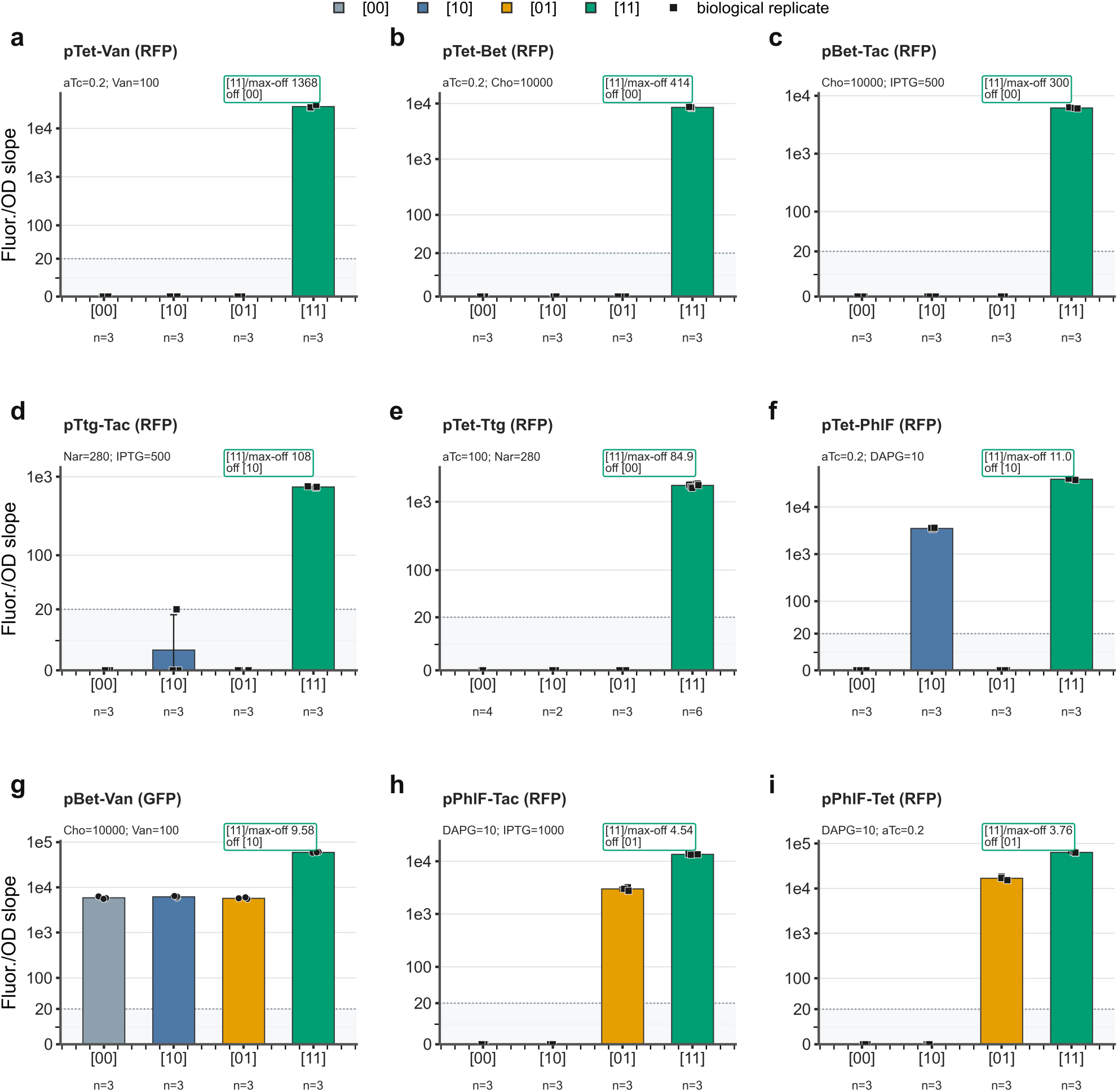
High-utility chemical-input AND-gate truth tables. Each panel shows one promoter-reporter construct for which the fully induced state [11] was significantly greater than [00], [10] and [01] by the predefined Holm-adjusted Welch operational screen and had [11]/max-off at least 2.0. These rows are the architecture-level representatives for the corresponding promoter designs: **a,** pTet-Van(RFP), **b,** pTet-Bet(RFP), **c,** pBet-Tac(RFP), **d,** pTtg-Tac(RFP), **e,** pTet-Ttg(RFP), **f,** pTet-PhlF(RFP), **g,** pBet-Van(GFP), **h,** pPhlF-Tac(RFP), and **i,** pPhlF-Tet(RFP). Bars show mean normalised fluorescence-versus-OD slope for [00], [10], [01] and [11]. Error bars denote s.d., and black symbols show biological-replicate means. Each panel reports the inducer pair, per-state n values, [11]/max-off ratio and highest off state. State colours are grey-blue [00], blue [10], orange [01] and green [11]. Panels use a symlog y-axis with a marked linear region around the empirical response floor so zero-clipped and low fitted slopes remain visible. Full Holm-adjusted one-sided Welch P values and complete reporter-specific truth tables, including weak, failed and non-classifiable rows, are reported in Supplementary Table S2 and Supplementary Figs. S13-S16.

### Sequence context and template controls define the usable design space

The architecture map and the supplementary sequence panel (Figure 1b and Supplementary Figure S19) highlight a recurrent sequence-level constraint among the long-operator designs. PhlO and TtgO were inserted downstream in reverse orientation when forward placement risked recreating promoter-like motifs within the early transcribed region. The present data do not prove that reverse orientation caused the successful behaviours, but they identify orientation as an actionable design variable for long binding sites.

To distinguish weak scaffold output from insertion-dependent effects, we compared each AND promoter with its corresponding single-input template promoter in the available reporter context (Figure 5 and Supplementary Table S12). The template-control source data provide the parent-template OFF/ON slopes, and the conditional activation-fold analysis compares template [1]/[0] with AND [11]/[01], holding the added-operator input ON while the template input is toggled. This comparison tests whether weak parent-promoter output alone explains the observed phenotypes. We also used the dual-channel measurements to diagnose whether biomass-associated fluorescence could affect the operational truth-table calls. For every well, the analysis fits fluorescence as F = β0 + β1 × OD_600_ over the exponential-growth window. OD-independent medium, optical, plate-reader, and well-background offsets enter β0, whereas the slope β1 reports the total biomass-normalised fluorescence-channel output of the complete promoter–reporter–host context. Therefore, any residual same-channel background that scales with OD600 would contribute to β1 rather than β0. Architectures measured in both EGFP and mCherry/RFP contexts allowed a same-channel alternate-reporter background diagnostic: the same optical channel measured in the alternate reporter construct estimates the OD-dependent same-channel background available under matched conditions.

**Fig. 5.**
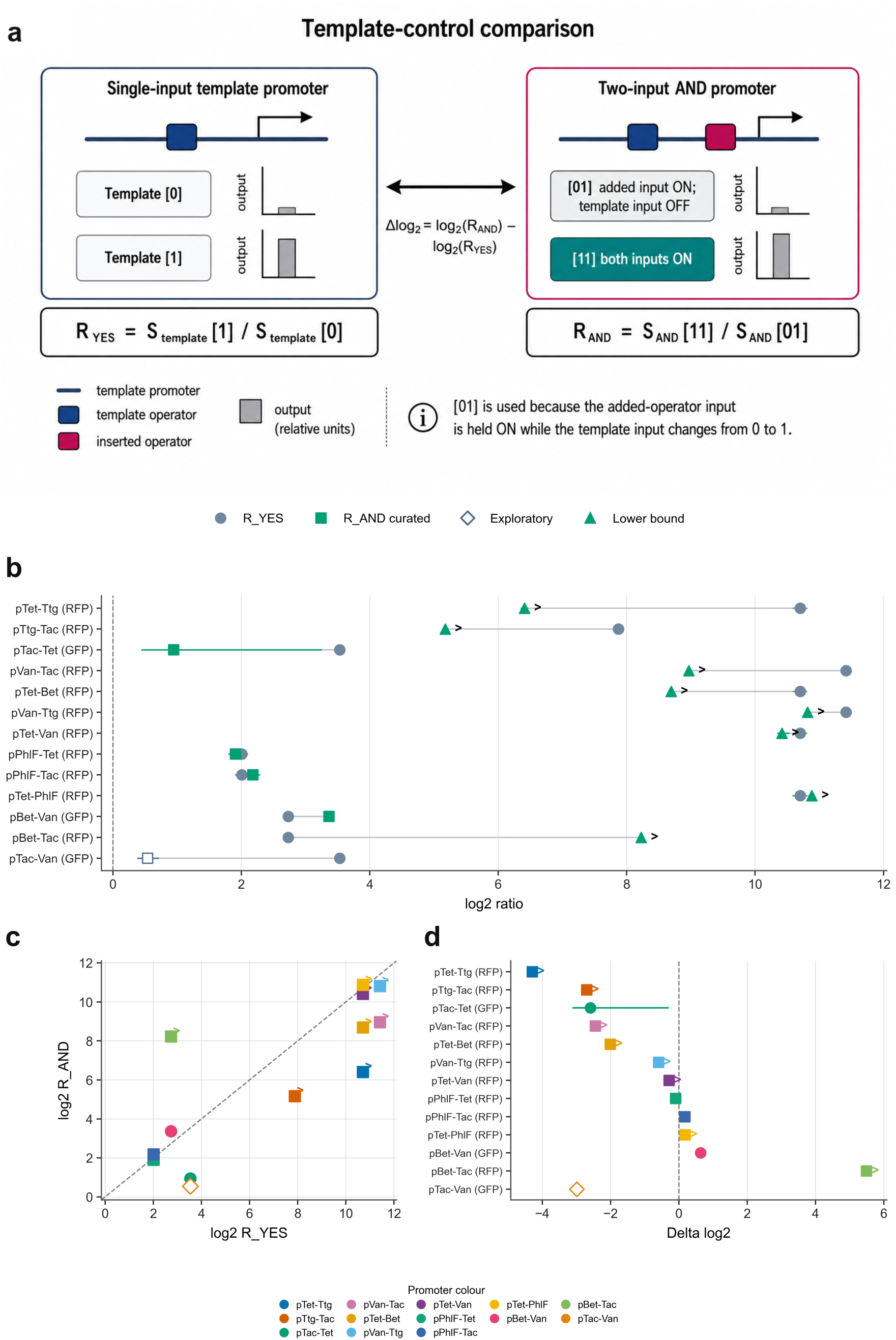
Template-promoter reference measurements and conditional activation-fold comparison. **a**, Conceptual definition of the template-promoter and AND-promoter ratios used for the conditional activation comparison. **b,** Paired log2 R_YES and log2 R_AND values for selected template/AND comparisons. Blue-grey circles show R_YES for the template promoter, green squares show curated R_AND values for the AND promoter, open diamonds mark exploratory pTac-Van, and green triangles with > labels mark lower-bound R_AND values. Horizontal lines show 95% bootstrap confidence intervals where available. **c,** Scatter plot of log2 R_YES against log2 R_AND for the same comparisons. Circles mark GFP rows, squares mark RFP/mCherry rows, open diamonds mark exploratory pTac-Van, and > labels mark lower-bound ratios. Marker colours in panels c and d identify individual promoter architectures, as shown in the shared promoter-colour legend below the panels. The dashed grey line marks y = **x,** where template and AND ratios are equal. **d,** Difference in conditional activation, Δlog2 = log2(R_AND) - log2(R_YES) = log2(AND [11]/[01]) - log2(template [1]/[0]). The y-axis lists the promoter-reporter comparisons, symbols use the same circle, square, open-diamond and > encodings as panel **c,** and values above zero indicate larger conditional activation in the AND-promoter context than in the template-promoter context. GFP and RFP are independent reporter outputs, not normalisation controls. For AND states, the first digit denotes the template-promoter input and the second digit denotes the added-operator input.

Supplementary Table S2b reports signed same-channel alternate-reporter β1 estimates in parentheses after the corresponding formal state means, and Supplementary Table S13 reports the state-level diagnostic and the matched margin checks. These diagnostics are not normalisation controls and were not subtracted from the truth tables. The operational benchmark therefore remains based on matched [00], [10], [01], and [11] comparisons within each complete promoter–reporter–host context. The template-control comparison does not assign mechanism, but it shows that several AND-promoter behaviours diverge from the corresponding parent scaffold after insertion of the second operator. No selected high-utility call changed after the same-channel background diagnostic.

## Discussion

This study engineered and quantitatively analysed a compact set of Marionette-derived two-input bacterial promoters, providing a source-traceable parts resource for chemical-input AND switching in *E. coli* and an empirical map of the constraints that determine which compact architectures remain useful after a second operator is inserted. Previous work established that bacterial combinatorial promoter behaviour depends on operator position, promoter geometry, regulatory context, and downstream sequence context^2–11^. Our contribution is to make those constraints directly comparable within a defined Marionette architecture set that retains successful, weak, failed, and non-classifiable designs. Several selected constructs produced clear high-utility truth tables, making them candidate Marionette-compatible two-input promoter parts for downstream circuit construction.

The benchmark also separates operational classification from engineering utility. The Holm-adjusted one-sided Welch screen provides a consistent way to identify promoter-reporter combinations in which [11] exceeds [00], [10], and [01]. However, that statistical criterion alone does not define a practically useful AND switch. Downstream circuits will impose their own thresholds for absolute [11] output, tolerated leakage, dynamic range, and cellular burden. We therefore report an Engineering Utility flag in Figure 2 and Supplementary Table S2, defined as operational Yes with [11]/max-off at least 2.0. This distinction is apparent in the wide range of [11]/max-off values among operational passes and in reporter-specific cases where [11] is statistically higher than the nearest off state but the separation remains modest (Figures 2 and 3 and Supplementary Table S2). For practical use, the full truth table and the highest off-state output are therefore more informative than a binary pass/fail label alone.

Within this benchmark, leakage in the single-input states was the main limitation. Several constructs produced high fully induced output but failed, or became weakly useful, because [10] or [01] approached [11]. This behaviour was especially informative because it showed that promoter strength and digitality can decouple. A strong scaffold can be unsuitable if the added operator does not suppress the relevant partially induced state, and a statistically passing promoter can remain a poor actuator if the nearest off state is too high. The most useful designs therefore suppress partial-state leakage before maximizing fully induced expression.

Scaffold identity and reciprocal architecture also had clear practical consequences. The pTet*-, pBetI-, and pPhlF-derived architectures tolerated added operators better than the pVanCC-derived architectures in the selected reporter contexts, whereas neither pVanCC-derived architecture produced a classifiable pass: pVan-Tac was N.C. because only one [11] biological replicate passed QC, and pVan-Ttg failed because [10] rose to the [11] level. This pattern should not be interpreted as a universal ranking of Marionette scaffolds, because the library is not exhaustive and the failed or N.C. designs might improve with altered spacing, operator choice, copy number, or expression context. It does show, however, that the template scaffold is not a neutral carrier of an added operator. Reciprocal designs must also be treated as distinct architectures: pTet-PhlF and pPhlF-Tet both passed, but their selected-reporter [11]/max-off ratios differed substantially, indicating that swapping the scaffold and inserted input does not preserve promoter behaviour.

Long operator insertions introduced an additional sequence-level constraint. In the final designs, PhlO and TtgO were placed in reverse orientation when forward placement risked recreating promoter-like motifs in or near the early transcribed region. The present dataset does not include matched forward-orientation controls, so it cannot prove that reverse orientation caused the successful behaviours. The data instead identify orientation as an actionable design variable and a testable hypothesis for long binding sites. Accordingly, this benchmark does not estimate an independent positional effect or claim an exhaustive positional rule. A direct follow-up would build matched forward and reverse PhlO and TtgO variants, scan operator position in single-base increments across at least one helical turn, and combine fluorescence outputs with RNAP and repressor occupancy measurements. Such experiments would test whether motif avoidance, steric interference, helical phasing, or transcriptional roadblocking explains the observed patterns.

The template-promoter controls help distinguish weak parent-scaffold output from effects introduced by the second operator (Figure 5 and Supplementary Table S12). These controls show that several AND-promoter behaviours cannot be explained simply by low template-promoter activation, without assigning a molecular mechanism to each construct. We also used the dual-channel, dual-reporter measurements to test whether biomass-associated same-channel background could bias the formal truth-table interpretation. This signal is the OD-dependent component that enters β1, in contrast to OD-independent medium, optical, plate-reader, and well offsets captured by β0. Supplementary Table S2b reports signed same-channel alternate-reporter β1 estimates in parentheses after each corresponding formal state mean where an independent alternate reporter comparison was available, and Supplementary Table S13 reports both the state-level diagnostic and the exact independent margin checks. For every selected high-utility architecture with an independent matched diagnostic, the corrected [11]-versus-off-state margin retained the same operational interpretation; the pTet-Bet(GFP) and pVan-Tac(RFP) cases remain N.C. because a required state has n = 1. These diagnostics are not reporter-free controls and cannot partition β1 into endogenous and reporter-derived molecular components. Interpreting them as background estimates assumes approximate spectral separation between the two reporters and does not correct for reporter-specific maturation, folding, brightness, burden, or transcript-context effects. We therefore use them only as robustness checks for state-dependent same-channel background, not as normalisation or subtraction terms. The operational benchmark remains defined as the total biomass-normalised channel output β1 of each complete promoter–reporter–host context, with leakage assessed by matched within-construct comparisons among [00], [10], [01], and [11]. Reporter-free plasmid controls would be needed only for absolute molecular attribution, not for the operational classification reported here. No selected high-utility call was changed by this diagnostic background diagnostic.

Reporter context was another important design variable. Four of seven dual-reporter architectures changed operational classification between reporter contexts, showing that promoter behaviour cannot be separated completely from the downstream genetic context. Possible explanations include changes in early transcribed-region sequence, transcript folding, RNAP clearance, local DNA supercoiling, or reporter-specific maturation and burden effects. Fluorescent-protein maturation kinetics are themselves reporter- and condition-dependent^10^. The present bulk fluorescence data do not distinguish among these mechanisms. We therefore do not rank GFP and RFP by a universal dynamic-range hierarchy; instead, we treat reporter identity as part of the promoter-gene context that must be validated for the intended application. A compact promoter validated upstream of one reporter should not be assumed to retain the same truth table upstream of another gene.

These results support a practical design workflow for compact two-input Marionette promoters. Beyond the specific parts reported here, the study shows that compact two-input promoters should be engineered as complete promoter-gene contexts rather than as modular sums of two single-input elements. Designers should begin from scaffolds with strong single-input repression, place the second operator with attention to local sequence context and long-operator orientation, test reciprocal arrangements as independent designs, quantify all four truth-table states rather than only fold induction, and validate the final construct upstream of the intended gene. The parts reported here provide an immediate starting point for Marionette-based circuits, while the benchmark defines the measurements needed to transfer the strategy to other hosts, regulators, and circuit loads.

## Methods

### Promoter framework and sensor context

We analysed engineered two-input combinatorial promoters derived from an extensively optimised small-molecule sensor set^13^. In the *E. coli* experimental strain used here (originally designated “Marionette”), the corresponding transcription-factor modules resided integrated into the chromosome. Therefore, the plasmids we characterised in this study carried the engineered output promoters and reporter cassettes rather than separate plasmid-encoded sensor regulators. Within the sensor set relevant to the present work, several regulator-promoter pairs corresponded to evolved variants, including PhlF-AM/pPhlF, VanR-AM/pVanCC, LacI-AM/pTac, BetI-AM/pBetI, and TtgR-AM/pTtg; the corresponding tet module is denoted pTet* in the original platform nomenclature^13^. Because previous optimisation acted on regulator sequence, regulator expression elements, and output promoter sequences, we interpreted the present constructs as combinations of optimised sensor modules rather than as combinations of entirely unmodified natural parts.

### Operator sequences and nomenclature

We used the original template names throughout: pVanCC, pBetI, pTtg, pPhlF, pTac, and pTet* ^13^. Engineered promoter names describe the template scaffold followed by the added regulatory input. Our curated comparative dataset comprised the architectures pTet-PhlF, pBet-Tac, pBet-Van, pPhlF-Tac, pPhlF-Tet, pTac-Tet, pTet-Bet, pTet-Ttg, pTet-Van, pTtg-Tac, pVan-Tac, and pVan-Ttg. Additional operator sequences included the 19 bp tetO2 site ^13^; the 30 bp VanO2 site containing two half-sites for cooperative VanR binding ^14^; the 22 bp BetO site identified by *de novo* screening ^15^; the 30 bp PhlO site; the 30 bp TtgO site ^16^; and the symmetric lacOsym site (20 bp) for stronger lac repressor binding ^17^.

### Plasmid construction

We assembled plasmids by Gibson assembly ^18^ using gene synthesis and PCR amplification with Q5 Master Mix DNA Polymerase (New England Biolabs, USA) and 40-bp homology domains. We purchased DNA from Integrated DNA Technologies (USA). We designed the plasmids P1 and P2 to have the same total size (4,774 bp) and a common backbone comprising a medium-copy-number pMB1 origin, an ampicillin-resistance cassette from pSEVA191 ^19^, and the spy terminator ^20^ to isolate the promoter. We amplified the pMB1 replicon, a ColE1 derivative, from pWT018e (Addgene #107886) ^21^ and introduced a T-to-C mutation in RNAII to lower copy number ^22, 23^. To create the memregulon libraries, we inserted a promoter-specific spacer followed by the promoter into the common backbone of both P1 and P2, except for the hybrid memregulon, in which the two plasmids carried different promoters. We introduced a terminator when plasmid-size constraints allowed it (Supplementary Information). Each promoter was followed by a promoter-specific spacer, the RiboJ ribozyme insulator ^24^, a weak ribosome-binding site (BBa_B0064), and then either mCherry or EGFP. The mCherry sequence derived from plasmid PAJ310 ^25^ and carried a C-terminal Gly-Ser-Gly extension to match the size of EGFP from pWT018a. After the fluorescent protein genes, we added the antibiotic-resistance cassettes CmR-dKanR and dCmR-KanR from pWT018e and pWT018a polycistronically using BBa_B0064 in P1 and P2, respectively. CmR-dKanR contains a chloramphenicol-resistance gene fused through a Gly-Ser-Gly-Ser-Gly-Ser linker to a kanamycin-resistance gene carrying the Asp208Ala inactivating mutation. dCmR-KanR contains a chloramphenicol-resistance gene carrying the His195Ala inactivating mutation, together with the Ala29Thr substitution that arose during cloning, fused to a kanamycin-resistance gene through the same linker.

### Promoter design

We generated two-input combinatorial promoters by modifying established sensor-template promoters with an additional regulatory operator. In most architectures, we introduced the added site in the downstream transcribed region at a defined position relative to the transcription start site, for example tetO2 at +20 in pTac-Tet, TtgO at +1 in pTet-Ttg, and TtgO at +5 in pVan-Ttg. Across the curated library, we therefore compared template-dependent effects of operator identity, position, orientation, and length. We inserted the long PhlO and TtgO sites in reverse orientation when sequence inspection indicated that their forward orientation could reconstruct promoter-like motifs, especially partial -10 or -35 elements, in the downstream region. Throughout the manuscript, engineered promoter names refer to the template scaffold first and the added regulatory input second, whereas template promoters retain the original nomenclature ^13^.

### Strains and culture conditions

We used *Escherichia coli* Top10 for all cloning steps. The strain carried the DH10B genotype[F− mcrA Δ(mrr-hsdRMS-mcrBC) Φ80lacZΔM15 ΔlacX74 recA1 endA1 araΔ139 Δ(ara, leu)7697 galU galK λ− rpsL (StrR) nupG]. We grew strains in LB medium at 37 °C with shaking at 200 rpm and supplemented the medium, as required, with carbenicillin (80 μg mL⁻¹), kanamycin (50 μg mL⁻¹), ampicillin (100 μg mL⁻¹), or chloramphenicol (34 μg mL⁻¹). For combinatorial promoter characterisation, we used *E. coli* Marionette strains ^13^ in the DH10B background. For these assays, we used M9 minimal medium containing 1× M9 salts, 100 μM CaCl2, 2 mM MgSO4, 10 μM FeSO4, 0.8% (v/v) glycerol, 0.2% casamino acids, 1 μg mL⁻¹ thiamine, 20 μg mL⁻¹ uracil, and 30 μg mL⁻¹ leucine, adjusted to pH 7.4. We used ampicillin for overnight growth and carbenicillin during characterisation experiments.

### Fluorescence characterisation of combinatorial promoters

We characterised plasmids carrying combinatorial promoters fused to either EGFP or mCherry in chloramphenicol-cured Marionette DH10B strains. We inoculated each strain from a single colony, either from a freshly transformed LB agar plate or from a glycerol stock, and grew cultures overnight at 37 °C in LB medium containing ampicillin (100 μg mL⁻¹). We then diluted overnight cultures 1:2000 into freshly prepared M9 medium containing the cognate inducer combination (Supplementary Table S6) and carbenicillin (80 μg mL⁻¹), and incubated them for a further 3 h. After growth, we transferred 200 μL of each culture to wells of a 96-well plate (Custom Corning Costar) and included both biological and technical replicates. We measured OD_600_ and fluorescence in an Infinite F500 plate reader (Tecan) at 37 °C with shaking, using repeated acquisitions every 15 min. For EGFP, we used a 465/35 nm excitation filter and a 530/25 nm emission filter; for mCherry, we used a 580/20 nm excitation filter and a 635/35 nm emission filter. For every well, we acquired OD_600_, GFP-channel fluorescence, and RFP-channel fluorescence at each time point; the construct’s cognate channel was used for the formal truth-table call, whereas independent alternate reporter constructs, when available under matched conditions, were used for same-channel background sensitivity checks. Raw time-course data were then analysed with a custom Python workflow implementing growth-model fitting and fluorescence-versus-OD regression.

### Truth-table analysis

We fitted growth dynamics with the Baranyi model and extracted reporter-specific fluorescence-versus-OD slopes over the OD_600_ window 0.15-0.70. Specifically, we fitted F = β0 + β1 × OD_600_ and used β1 as the operational reporter output, following the established exponential-phase fluorescence-versus-turbidity approach of Alper et al.^26^ These slopes were averaged per biological replicate and then assembled into complete four-state truth tables [00], [10], [01], and [11]. A complete non-toxicity grid required acceptable growth and a valid fluorescence-versus-OD fit for every state in the four-state set. When multiple non-zero dose pairs were available, we prioritised the highest pair that retained such a complete grid. Error bars in the truth-table plots denote the s.d. across biological replicate means, and state-specific replicate counts are reported in Figure 2 and Supplementary Table S2. Promoter-reporter rows with fewer than two QC-passing biological-replicate means in any state required for a Welch comparison were labelled N.C. rather than pass or fail. Engineering Utility was defined as True only for rows with an operational Yes call and [11]/max-off at least 2.0. We classified a construct as passing the predefined operational criterion when the fully induced [11] state exceeded each non-fully induced state after Holm correction of one-sided Welch comparisons (P ≤ 0.05). Because we clipped fitted fluorescence-versus-OD slopes at zero when negative, an off-state mean of 0.00 a.u./OD indicates a non-positive fitted response after normalisation rather than literal zero fluorescence. Accordingly, for [11]/max-off calculations, we reported any construct with max off = 0.00 a.u./OD as a lower bound using an empirical response floor of 20.49 a.u./OD, corresponding to the smallest positive off-state biological replicate response observed in the selected RFP dataset. OD-independent medium, optical, plate-reader, and well-background offsets contribute to β0 rather than β1. β1 reports the total biomass-normalised fluorescence-channel output of the complete promoter–reporter–host context. Both GFP and RFP channels were acquired in every assay, although only the cognate reporter channel was used for each formal truth-table call. Same-channel alternate-reporter background sensitivity checks are described below. We define leakage by matched within-construct comparisons among [00], [10], [01], and [11].

### Same-channel alternate-reporter background diagnostic

We define biomass-associated same-channel background as the OD-dependent signal in the queried optical channel that remains when the cognate reporter for that channel is absent. For promoter architectures measured in independent EGFP and mCherry/RFP contexts, we used the same optical channel in the alternate reporter construct to estimate this component for each formal reporter measurement. Thus, for EGFP constructs, the GFP-channel slope measured from the corresponding mCherry/RFP construct provided the same-channel background estimate, whereas for mCherry/RFP constructs, the RFP-channel slope measured from the corresponding EGFP construct provided the same-channel background estimate. We applied the same fluorescence-versus-OD regression, truth-table state definitions, OD_600_ window, and biological-replicate grouping used for the formal analysis. We required the same promoter architecture, truth-table state, inducer identities and concentrations, optical channel, and slope-extraction procedure for an exact independent match. We did not use same-culture non-cognate channel readings as independent alternate-reporter background estimates because those readings are spectral cross-channel checks rather than independent same-channel measurements; such cases were flagged separately in the source data. Supplementary Table S2b reports signed same-channel alternate-reporter β1 estimates in parentheses after the corresponding formal state means where independent alternate reporter comparisons were available. For exact matched states with n > 1 and a positive formal reading, we also expressed the positive background estimate as a fraction of the corresponding formal same-state β1 value and tested whether state-dependent background differences could affect the [11] versus off-state margins. These diagnostics were interpreted only as sensitivity checks because they assume approximate spectral separation and cannot correct for reporter-specific maturation, brightness, folding, burden, or transcript-context effects. They were not used to normalise, subtract, or reclassify the formal truth tables.

### Template-promoter control and conditional activation-fold analysis

We analysed six single-input template promoters corresponding to the scaffolds used in the AND-promoter panel: pTet* for pTet-PhlF, pTet-Bet, pTet-Ttg, and pTet-Van; pBetI for pBet-Tac and pBet-Van; pPhlF for pPhlF-Tac and pPhlF-Tet; pTac for pTac-Tet and exploratory pTac-Van; pTtg for pTtg-Tac; and pVanCC for pVan-Tac and pVan-Ttg. For each template/AND pair, the first digit in the AND truth table denotes the template-promoter input and the second digit denotes the added-operator input. We calculated the single-promoter YES-gate activation ratio as R_YES = S_template[1]/S_template[0] and the AND-promoter conditional activation ratio as R_AND = S_AND[11]/S_AND[01]. We used [01] rather than [00] because the added-operator input is held ON, so the added repressor is relieved while only the template input changes from 0 to 1. Template ON states were matched to the same template-inducer concentration used in each AND truth table whenever available; otherwise the nearest available non-toxic ON condition was used and flagged in Supplementary Table S12. Uncertainty was estimated by 10,000 bootstrap resamples of biological-replicate fluorescence-versus-OD slopes within each state using random seed 1001. Ratios are reported as log2 values with 95% bootstrap confidence intervals. If a denominator mean was non-positive because fitted slopes had been clipped at zero, the ratio was reported as a lower bound; for plotting lower bounds on log axes only, we used a reporter-specific response floor equal to the smallest strictly positive biological-replicate slope in the combined included dataset for that reporter. The same measurement definition applies to the template-control analysis. Regression places OD-independent plate-reader, medium, optical, and well-background offsets in β0, whereas β1 reports the total biomass-normalised fluorescence-channel output. Because both optical channels were acquired, the non-cognate channel provides an internal check on combined channel background and possible spectral bleed-through under the same assay conditions. These values are not used for normalisation or subtraction; the analysis compares matched within-reporter, within-construct states. Reporter-free controls would be needed only to partition the cognate-channel slope into endogenous and reporter-derived molecular components.

## Availability of data and materials

The datasets supporting the conclusions of this article are included within the article and its additional files. The submitted source-data workbook is source_data.xlsx. The reproducibility scripts for the quantitative analyses are included as code_ocean_scripts.zip, and the SnapGene plasmid files are included as snapgene_files.zip. Plasmids and strains are available from the corresponding author on reasonable request.

## Acknowledgments

AJ was funded by the Spanish Agencia Estatal de Investigación (AEI) through grant PID2023-151174NB-I00 funded by MICIU/AEI/10.13039/501100011033 and grant PID2020-118436GB-I00 funded by MCIN/AEI/10.13039/501100011033; AJ and SP were funded by the Biotechnology and Biological Sciences Research Council [BB/P020615/1] (EVO-ENGINE) and by the Biotechnology and Biological Sciences Research Council and the Engineering and Physical Sciences Research Council [BB/M017982/1] (Warwick Integrative Synthetic Biology Centre, WISB); AJ was funded by the European Union Seventh Framework Programme (FET Proactive) under grant agreement No. 610730 (EVOPROG); by the European Union under the Horizon Europe research and innovation programme through the European Innovation Council (EIC Pathfinder) under grant agreement No. 101070948 (PhotoSynH2); by the Department of the Navy, Office of Naval Research, under award No. N62909-23-1-2008 (Office of Naval Research Global, ONRG); by the ”la Caixa” Foundation under the project code HR22-00405 [EvoPunch]; and by a departmental allocation from the School of Life Sciences, University of Warwick.

## Author contributions

**A.J.:** Conceptualization, Methodology, Software, Formal analysis, Visualization, Writing – original draft, Writing – review & editing, Funding acquisition, Supervision. **S.P.:** Investigation, Validation, Formal analysis, Writing – review & editing. Both authors approved the final manuscript.

