## Supplementary Information for "Engineering compact bacterial combinatorial promoters for two-input chemical AND switching"

#### Supplementary Figures

Supplementary Fig. S1 | Dynamics and fit diagnostics for pTet-PhlF.
Supplementary Fig. S2 | Dynamics and fit diagnostics for pBet-Tac.
Supplementary Fig. S3 | Dynamics and fit diagnostics for pBet-Van.
Supplementary Fig. S4 | Dynamics and fit diagnostics for pPhlF-Tac.
Supplementary Fig. S5 | Dynamics and fit diagnostics for pPhlF-Tet.
Supplementary Fig. S6 | Dynamics and fit diagnostics for pTac-Tet.
Supplementary Fig. S7 | Dynamics and fit diagnostics for pTet-Bet.
Supplementary Fig. S8 | Dynamics and fit diagnostics for pTet-Ttg.
Supplementary Fig. S9 | Dynamics and fit diagnostics for pTet-Van.
Supplementary Fig. S10 | Dynamics and fit diagnostics for pTtg-Tac.
Supplementary Fig. S11 | Dynamics and fit diagnostics for pVan-Tac.
Supplementary Fig. S12 | Dynamics and fit diagnostics for pVan-Ttg.
Supplementary Fig. S13 | Reporter-specific truth tables matching Supplementary Table S2.
Supplementary Fig. S14 | Reporter-specific truth tables matching Supplementary Table S2.
Supplementary Fig. S15 | Reporter-specific truth tables matching Supplementary Table S2.
Supplementary Fig. S16 | Reporter-specific truth tables matching Supplementary Table S2.
Supplementary Fig. S17 | Architecture of the experimentally tested pTac-Van combinatorial promoter.
Supplementary Fig. S18 | Coverage of the scaffold-by-input design space used in this study.
Supplementary Fig. S19 | Sequence alignment around the -10 region and +1 site in reverse-operator examples.

Supplementary Tables

Supplementary Table S1 | Distributed source-data and reproducibility files.
Supplementary Table S2 | Statistics for all 19 curated two-input combinatorial promoter-reporter truth tables, including the Engineering Utility flag.
Supplementary Table S3 | Excluded entries and quality-control summary for the primary analysis.
Supplementary Table S4 | Primers used in this work.
Supplementary Table S5 | Promoter sequences.
Supplementary Table S6 | Selected inducer concentrations and truth-table digit mapping.
Supplementary Table S7 | Compact promoter sequence annotations for engineered promoter architectures.
Supplementary Table S8 | Cross-reference between characterised promoter/reporter constructs.
Supplementary Table S9 | Representative mCherry plasmid sequence.
Supplementary Table S10 | Representative EGFP plasmid sequence.
Supplementary Table S11 | Overview of the engineered promoter architectures.
Supplementary Table S12 | Template-promoter YES-gate activation ratios and AND-promoter conditional activation ratios.
Supplementary Table S13 | Same-channel alternate-reporter background diagnostic and margin checks.

#### Supplementary Text

This Supplementary Information documents the inducer-conditioned truth-table analysis of the Tecan plate-reader datasets used to evaluate the engineered two-input combinatorial promoters. The response variable is the fitted normalised fluorescence (a.u./OD), extracted separately for GFP and RFP reporters. As in the main text, truth-table language is operational in inducer space: the experimentally controlled inputs are inducer concentrations, whereas promoter regulation in the stricter biophysical sense depends on intracellular concentrations of active transcription factors and promoter occupancy. We restrict the curated comparative analysis to the 12 pre-specified architectures that yielded complete four-state truth tables, while also documenting the experimentally tested exploratory pTac-Van construct separately. Supplementary Figures S1-S12 provide compact dynamics and fit diagnostics, Supplementary Figures S13-S19 provide reporter-specific truth tables, exploratory architecture, coverage, and sequence-context views, and Supplementary Tables S1-S13 provide replicate provenance, summary statistics, annotated promoter architectures, primer sequences, representative plasmid sequences, and construct-to-figure mapping. Promoter names follow the nomenclature used in the main text: engineered promoters are named by scaffold followed by the added input, whereas template promoters retain their original designations (pVanCC, pBetI, pTtg, pPhlF, pTac, and pTet*)13.

Dynamics and fit diagnostics for curated combinatorial designs. Only the 12 curated combinatorial promoters selected for truth-table evaluation are shown here. Each promoter is condensed to a single page with $OD_{600}$, GFP, and RFP trajectories overlaid by inducer state ([00], [10], [01], [11]), the fitted growth model overlaid on $OD_{600}$, and a fluorescence-versus-OD diagnostic panel for the reporter used in the promoter call. Fluorescence normalisation is assessed against measured $OD_{600}$ within the OD window 0.15-0.70. Single-input template controls and exploratory constructs are summarised separately in the revised main-text figures and supplementary overview panels.

#### Supplementary Figures

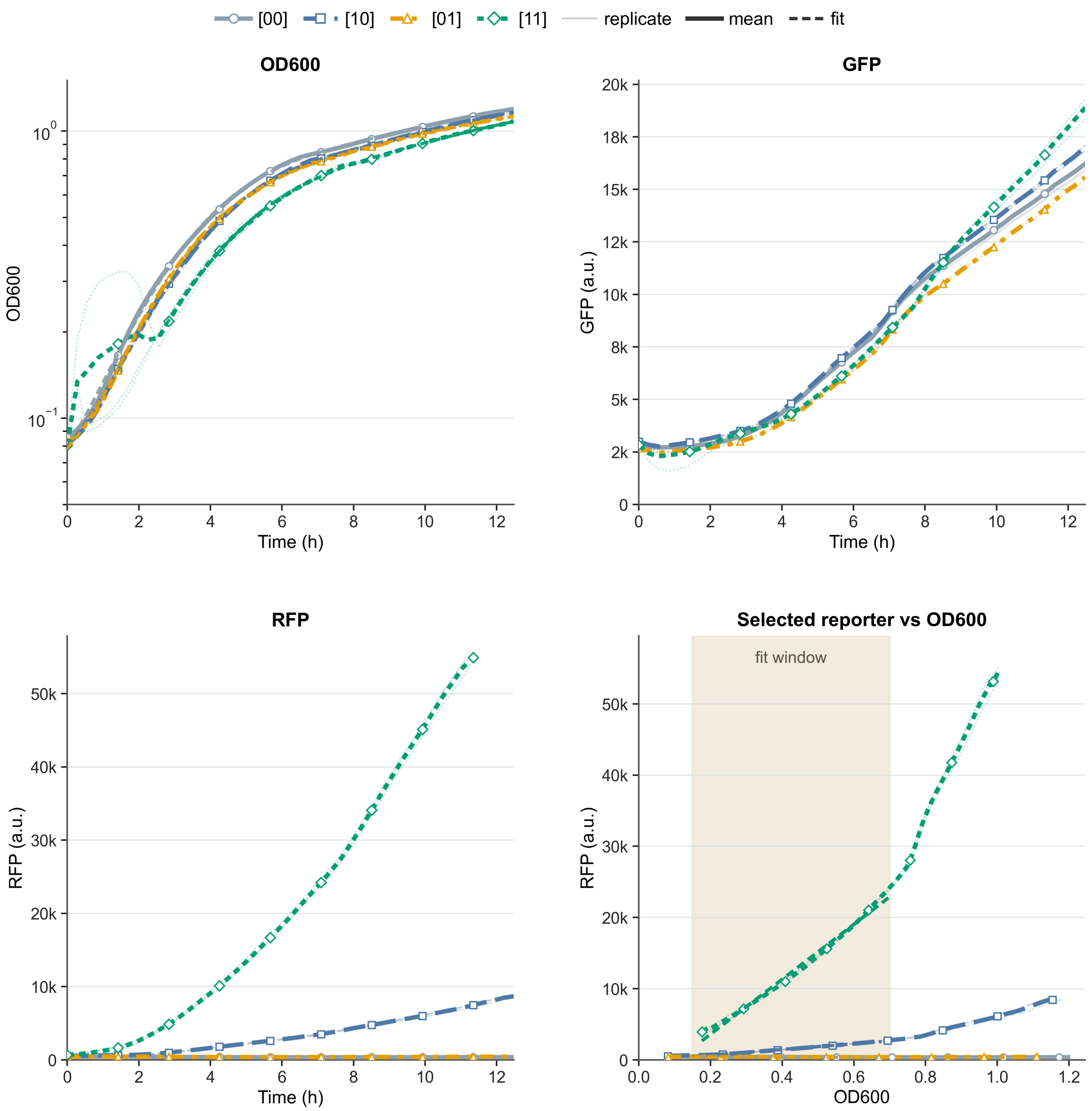

**Supplementary Fig. S1 | Dynamics and fit diagnostics for pTet-PhlF.** The four subpanels show $OD_{600}$ versus time with the growth fit, GFP versus time, RFP versus time, and selected-reporter fluorescence versus $OD_{600}$. Selected reporter: RFP. Inducer pair, using the source grid label with units in Supplementary Table S6: aTc=0.2; DAPG=10. Biological replicates per state: [00] n = 3, [10] n = 3, [01] n = 3, [11] n = 3. GFP and RFP are independent reporter outputs, not normalisation controls. The fluorescence-versus-OD diagnostic is the panel used to derive the normalised fluorescence for the selected reporter.

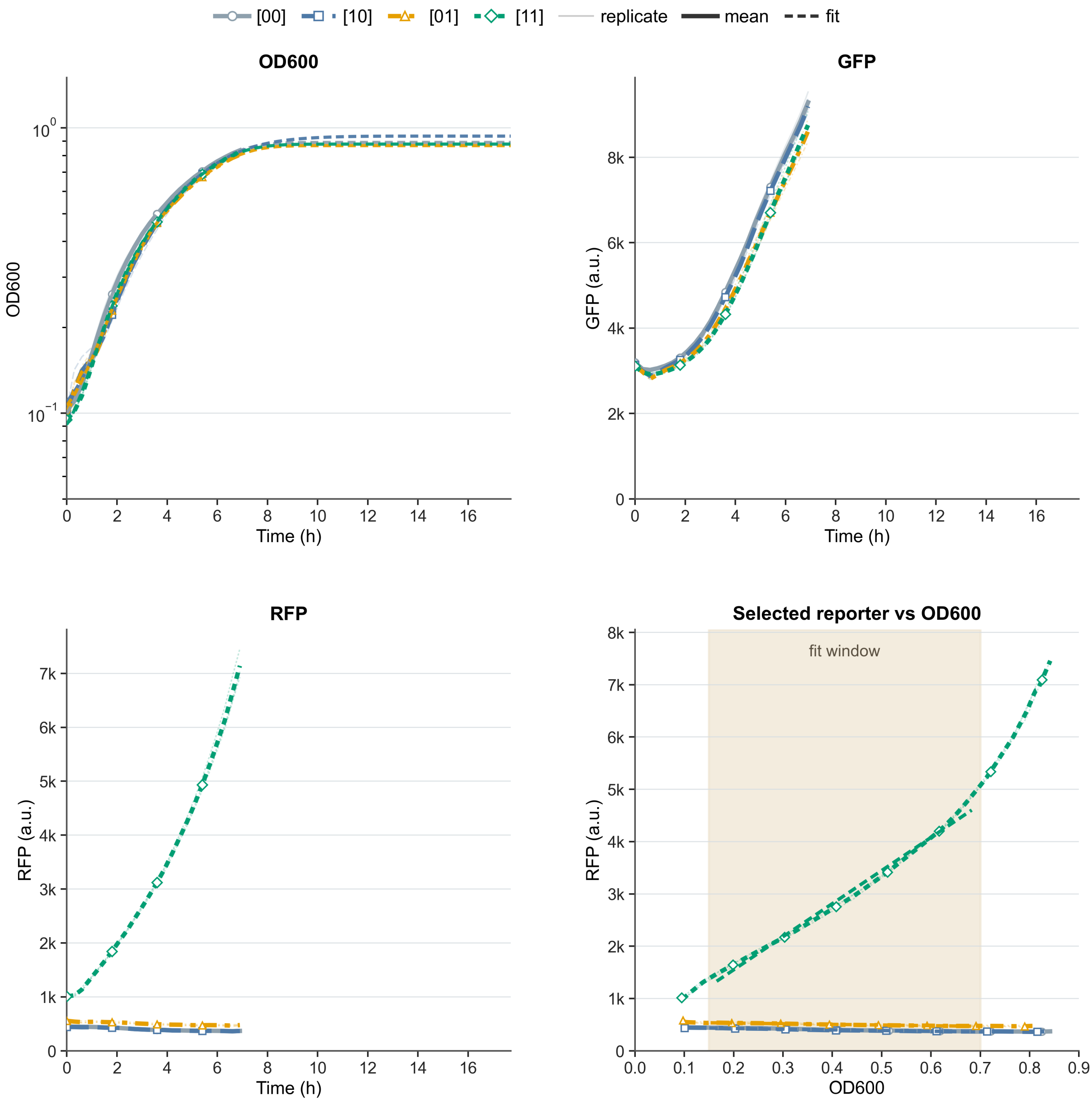

**Supplementary Fig. S2 | Dynamics and fit diagnostics for pBet-Tac.** The four subpanels show $OD_{600}$ versus time with the growth fit, GFP versus time, RFP versus time, and selected-reporter fluorescence versus $OD_{600}$. Selected reporter: RFP. Inducer pair, using the source grid label with units in Supplementary Table S6: Cho=10000; IPTG=500. Biological replicates per state: [00] n = 3, [10] n = 3, [01] n = 3, [11] n = 3. GFP and RFP are independent reporter outputs, not normalisation controls. The fluorescence-versus-OD diagnostic is the panel used to derive the normalised fluorescence for the selected reporter.

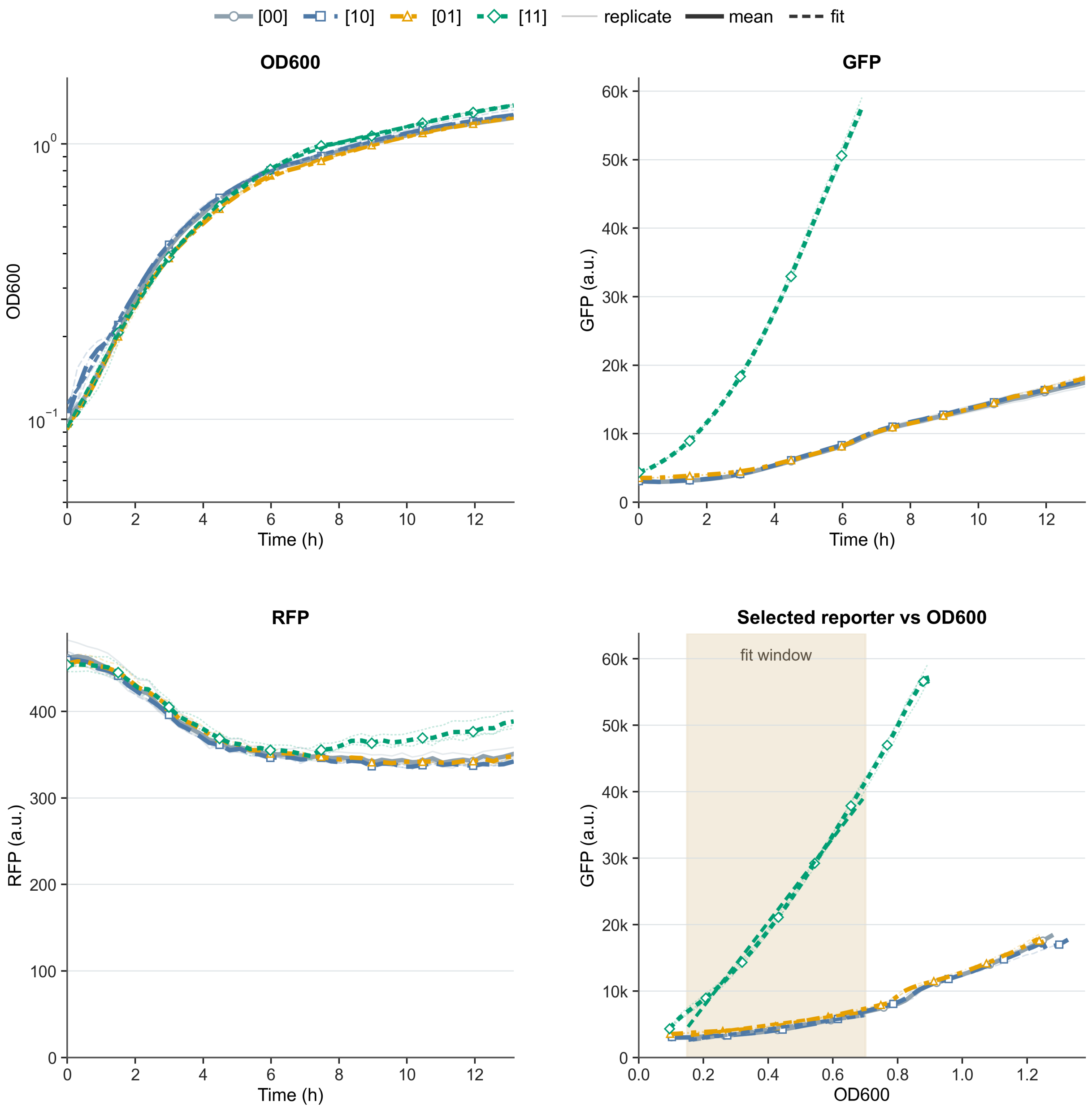

**Supplementary Fig. S3 | Dynamics and fit diagnostics for pBet-Van.** The four subpanels show $OD_{600}$ versus time with the growth fit, GFP versus time, RFP versus time, and selected-reporter fluorescence versus $OD_{600}$. Selected reporter: GFP. Inducer pair, using the source grid label with units in Supplementary Table S6: Cho=10000; Van=100. Biological replicates per state: [00] n = 3, [10] n = 3, [01] n = 3, [11] n = 3. GFP and RFP are independent reporter outputs, not normalisation controls. The fluorescence-versus-OD diagnostic is the panel used to derive the normalised fluorescence for the selected reporter.

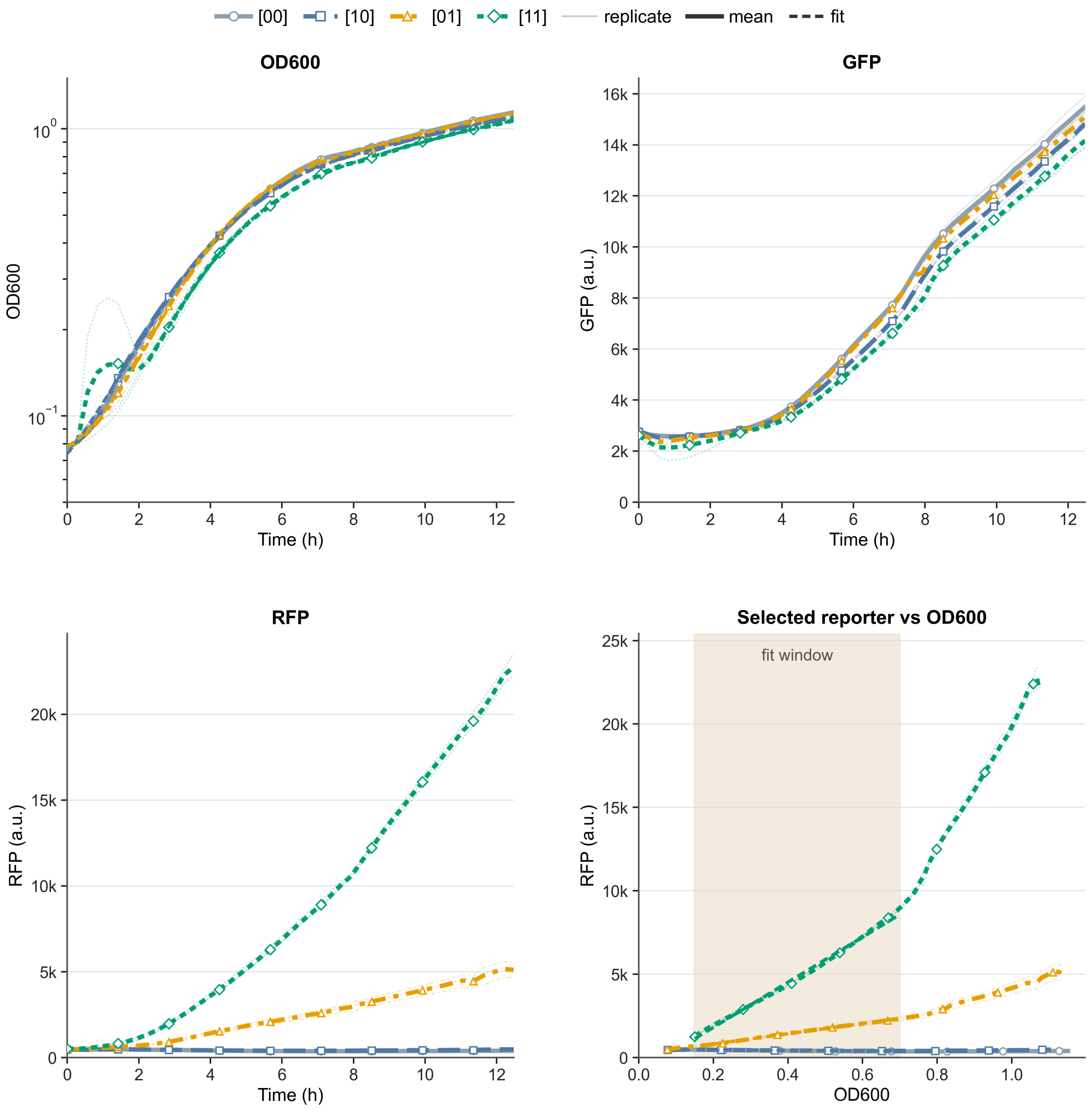

**Supplementary Fig. S4 | Dynamics and fit diagnostics for pPhlF-Tac.** The four subpanels show $OD_{600}$ versus time with the growth fit, GFP versus time, RFP versus time, and selected-reporter fluorescence versus $OD_{600}$. Selected reporter: RFP. Inducer pair, using the source grid label with units in Supplementary Table S6: DAPG=10; IPTG=1000. Biological replicates per state: [00] n = 3, [10] n = 3, [01] n = 3, [11] n = 3. GFP and RFP are independent reporter outputs, not normalisation controls. The fluorescence-versus-OD diagnostic is the panel used to derive the normalised fluorescence for the selected reporter.

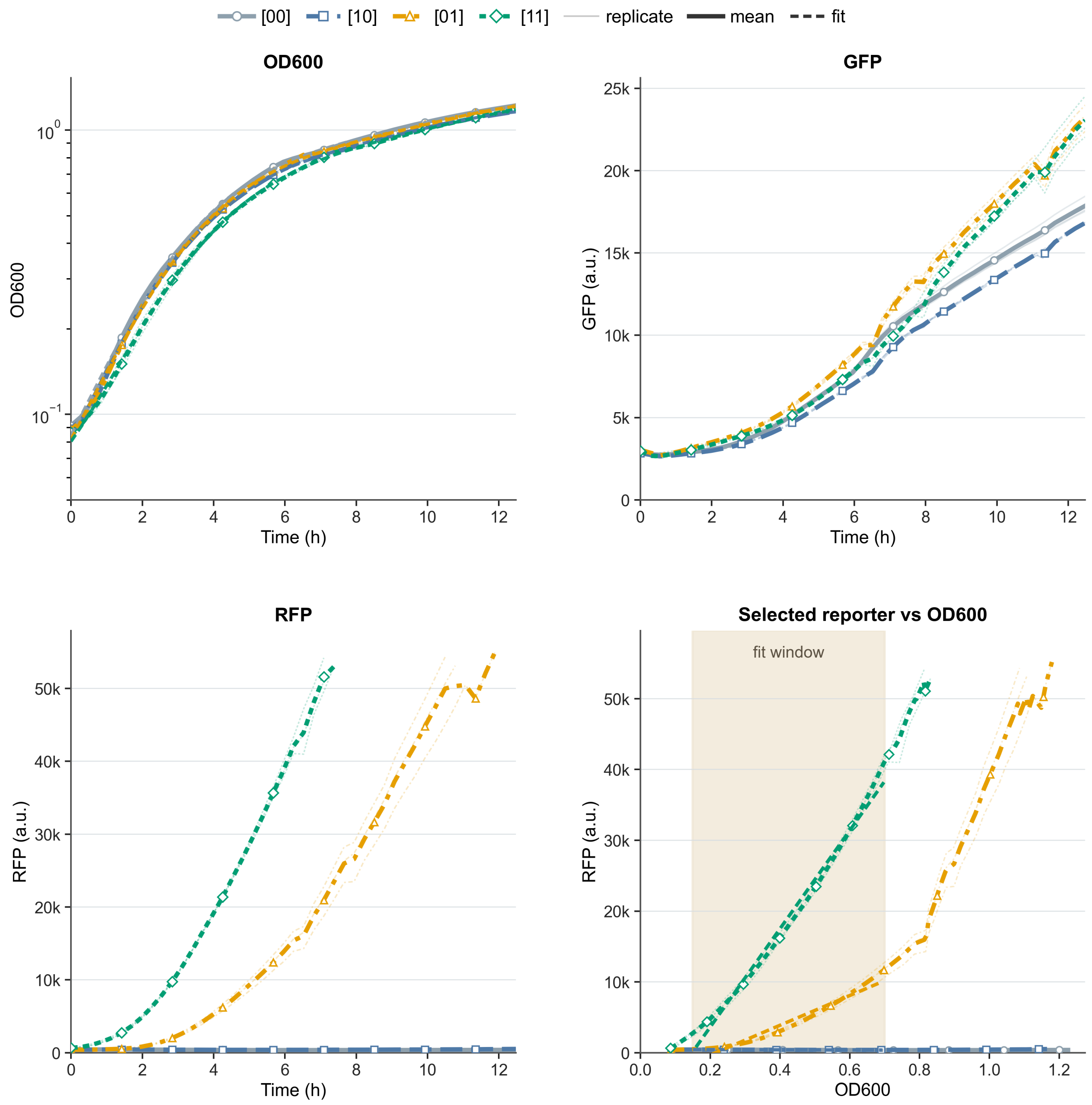

**Supplementary Fig. S5 | Dynamics and fit diagnostics for pPhlF-Tet.** The four subpanels show $OD_{600}$ versus time with the growth fit, GFP versus time, RFP versus time, and selected-reporter fluorescence versus $OD_{600}$. Selected reporter: RFP. Inducer pair, using the source grid label with units in Supplementary Table S6: DAPG=10; aTc=0.2. Biological replicates per state: [00] n = 3, [10] n = 3, [01] n = 3, [11] n = 3. GFP and RFP are independent reporter outputs, not normalisation controls. The fluorescence-versus-OD diagnostic is the panel used to derive the normalised fluorescence for the selected reporter.

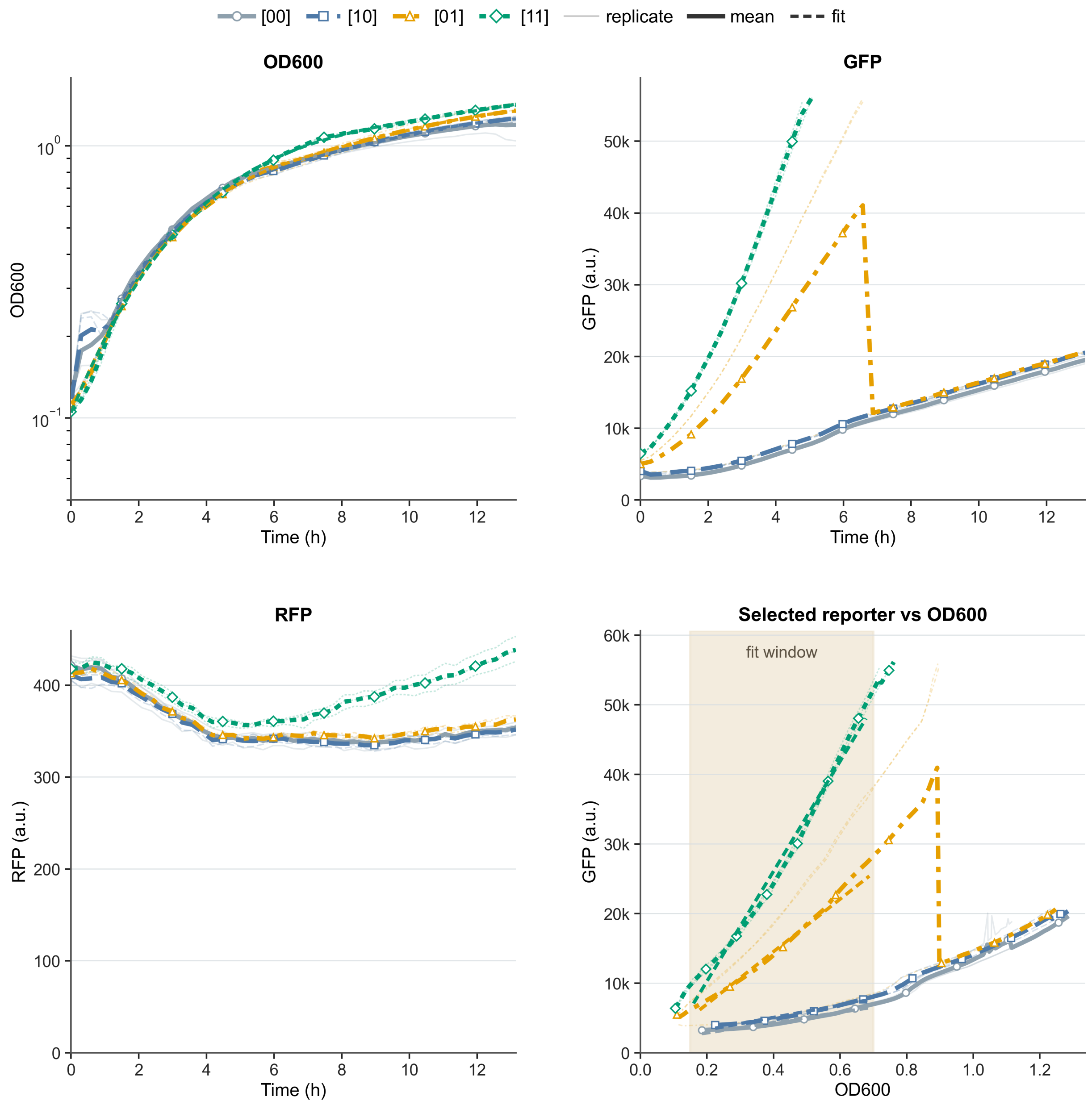

**Supplementary Fig. S6 | Dynamics and fit diagnostics for pTac-Tet.** The four subpanels show $OD_{600}$ versus time with the growth fit, GFP versus time, RFP versus time, and selected-reporter fluorescence versus $OD_{600}$. Selected reporter: GFP. Inducer pair, using the source grid label with units in Supplementary Table S6: IPTG=1000; aTc=0.2. Biological replicates per state: [00] n = 3, [10] n = 3, [01] n = 3, [11] n = 3. GFP and RFP are independent reporter outputs, not normalisation controls. The fluorescence-versus-OD diagnostic is the panel used to derive the normalised fluorescence for the selected reporter.

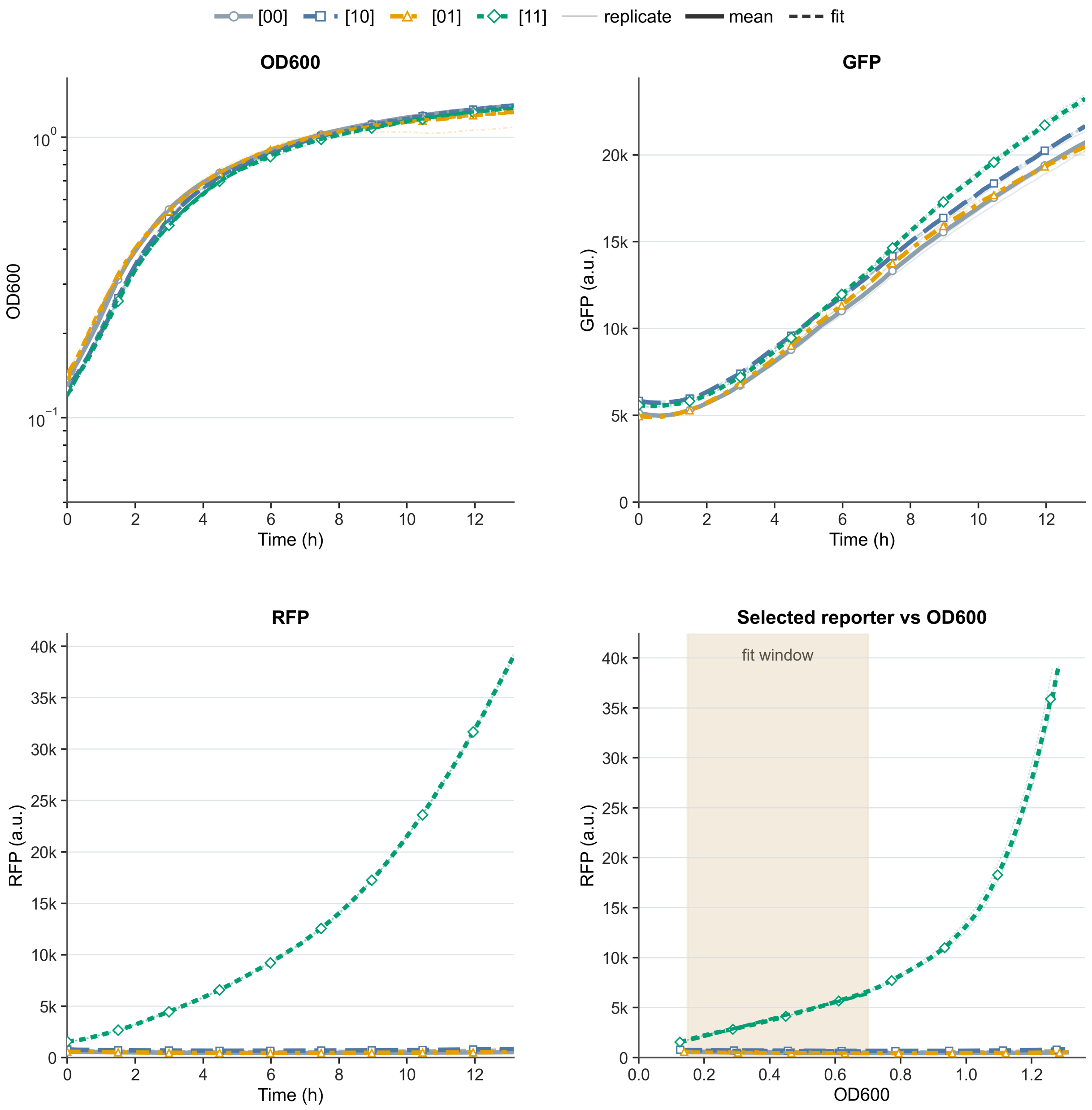

**Supplementary Fig. S7 | Dynamics and fit diagnostics for pTet-Bet.** The four subpanels show $OD_{600}$ versus time with the growth fit, GFP versus time, RFP versus time, and selected-reporter fluorescence versus $OD_{600}$. Selected reporter: RFP. Inducer pair, using the source grid label with units in Supplementary Table S6: aTc=0.2; Cho=10000. Biological replicates per state: [00] n = 3, [10] n = 3, [01] n = 3, [11] n = 3. GFP and RFP are independent reporter outputs, not normalisation controls. The fluorescence-versus-OD diagnostic is the panel used to derive the normalised fluorescence for the selected reporter.

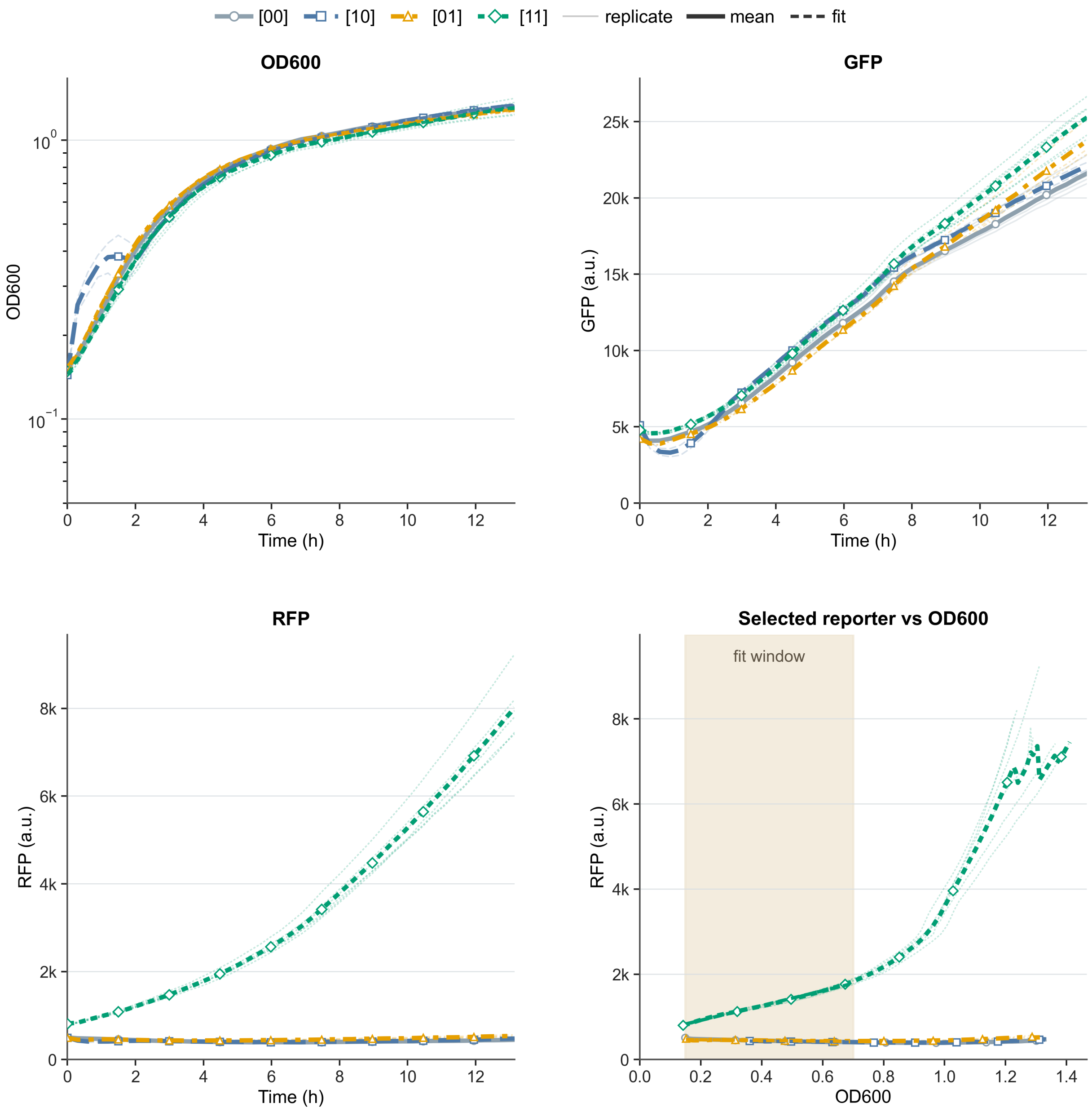

**Supplementary Fig. S8 | Dynamics and fit diagnostics for pTet-Ttg.** The four subpanels show $OD_{600}$ versus time with the growth fit, GFP versus time, RFP versus time, and selected-reporter fluorescence versus $OD_{600}$. Selected reporter: RFP. Inducer pair, using the source grid label with units in Supplementary Table S6: aTc=100; Nar=280. Biological replicates per state: [00] n = 4, [10] n = 2, [01] n = 3, [11] n = 6. GFP and RFP are independent reporter outputs, not normalisation controls. The fluorescence-versus-OD diagnostic is the panel used to derive the normalised fluorescence for the selected reporter.

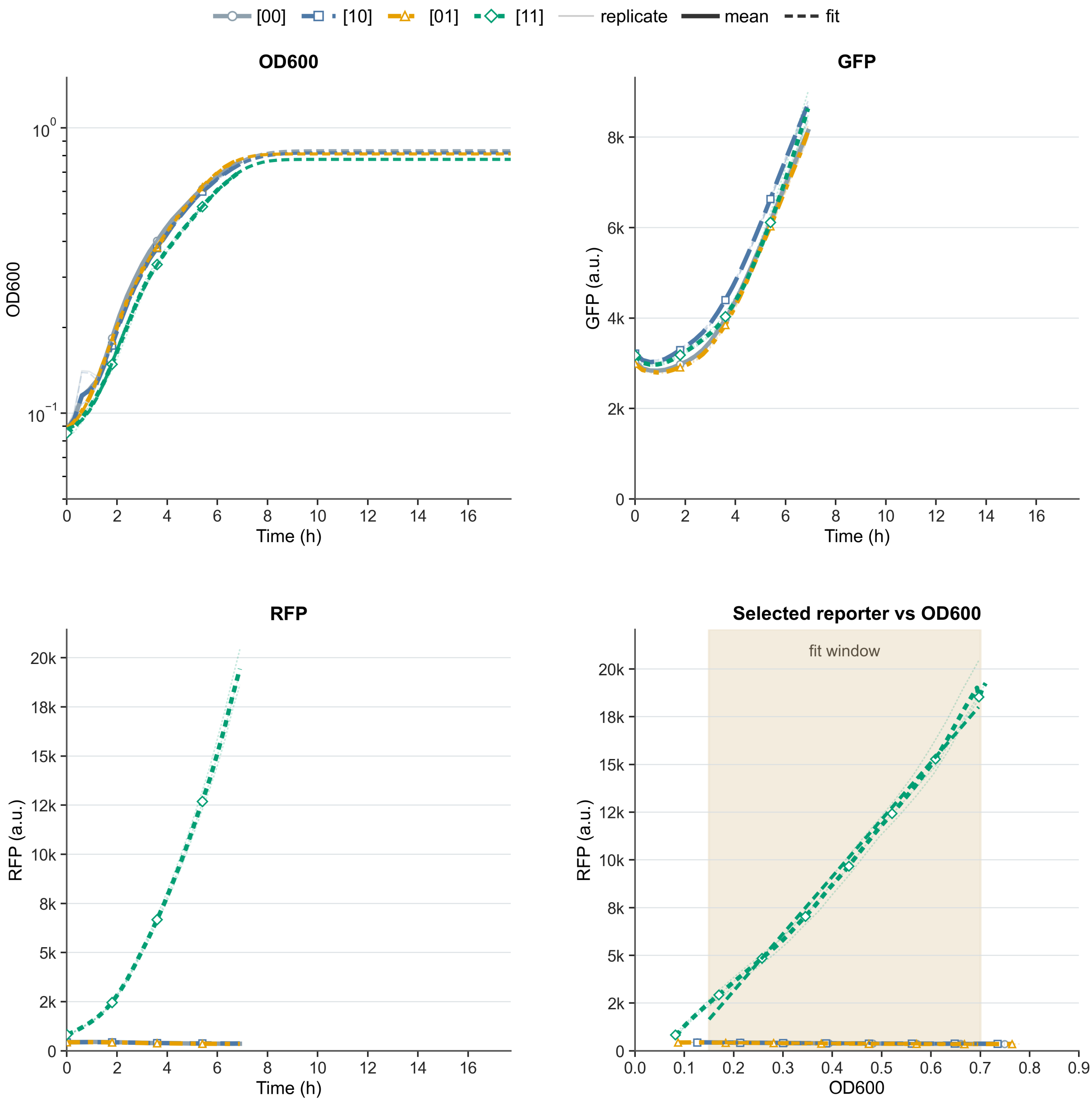

**Supplementary Fig. S9 | Dynamics and fit diagnostics for pTet-Van.** The four subpanels show $OD_{600}$ versus time with the growth fit, GFP versus time, RFP versus time, and selected-reporter fluorescence versus $OD_{600}$. Selected reporter: RFP. Inducer pair, using the source grid label with units in Supplementary Table S6: aTc=0.2; Van=100. Biological replicates per state: [00] n = 3, [10] n = 3, [01] n = 3, [11] n = 3. GFP and RFP are independent reporter outputs, not normalisation controls. The fluorescence-versus-OD diagnostic is the panel used to derive the normalised fluorescence for the selected reporter.

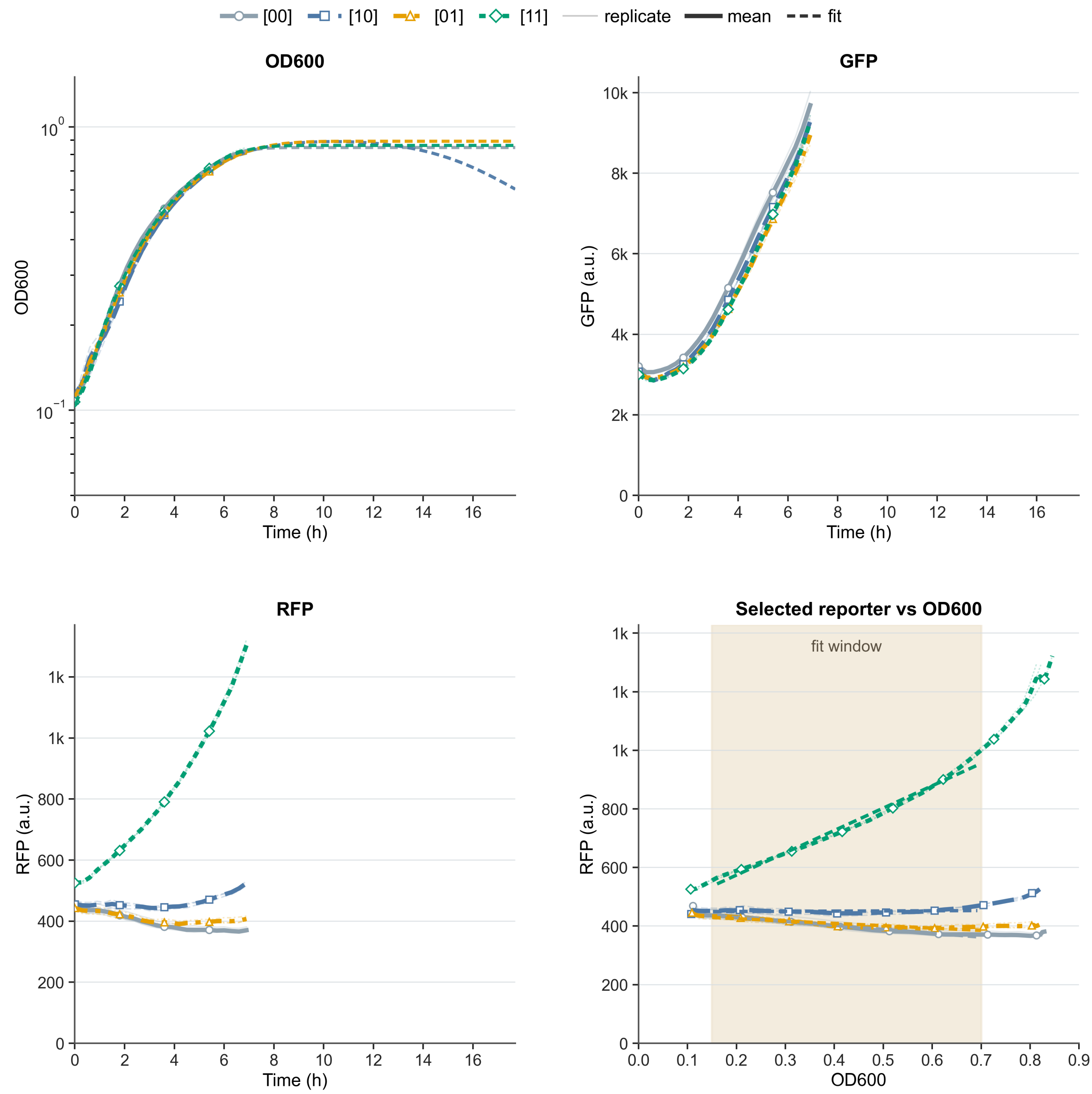

**Supplementary Fig. S10 | Dynamics and fit diagnostics for pTtg-Tac.** The four subpanels show $OD_{600}$ versus time with the growth fit, GFP versus time, RFP versus time, and selected-reporter fluorescence versus $OD_{600}$. Selected reporter: RFP. Inducer pair, using the source grid label with units in Supplementary Table S6: Nar=280; IPTG=500. Biological replicates per state: [00] n = 3, [10] n = 3, [01] n = 3, [11] n = 3. GFP and RFP are independent reporter outputs, not normalisation controls. The fluorescence-versus-OD diagnostic is the panel used to derive the normalised fluorescence for the selected reporter.

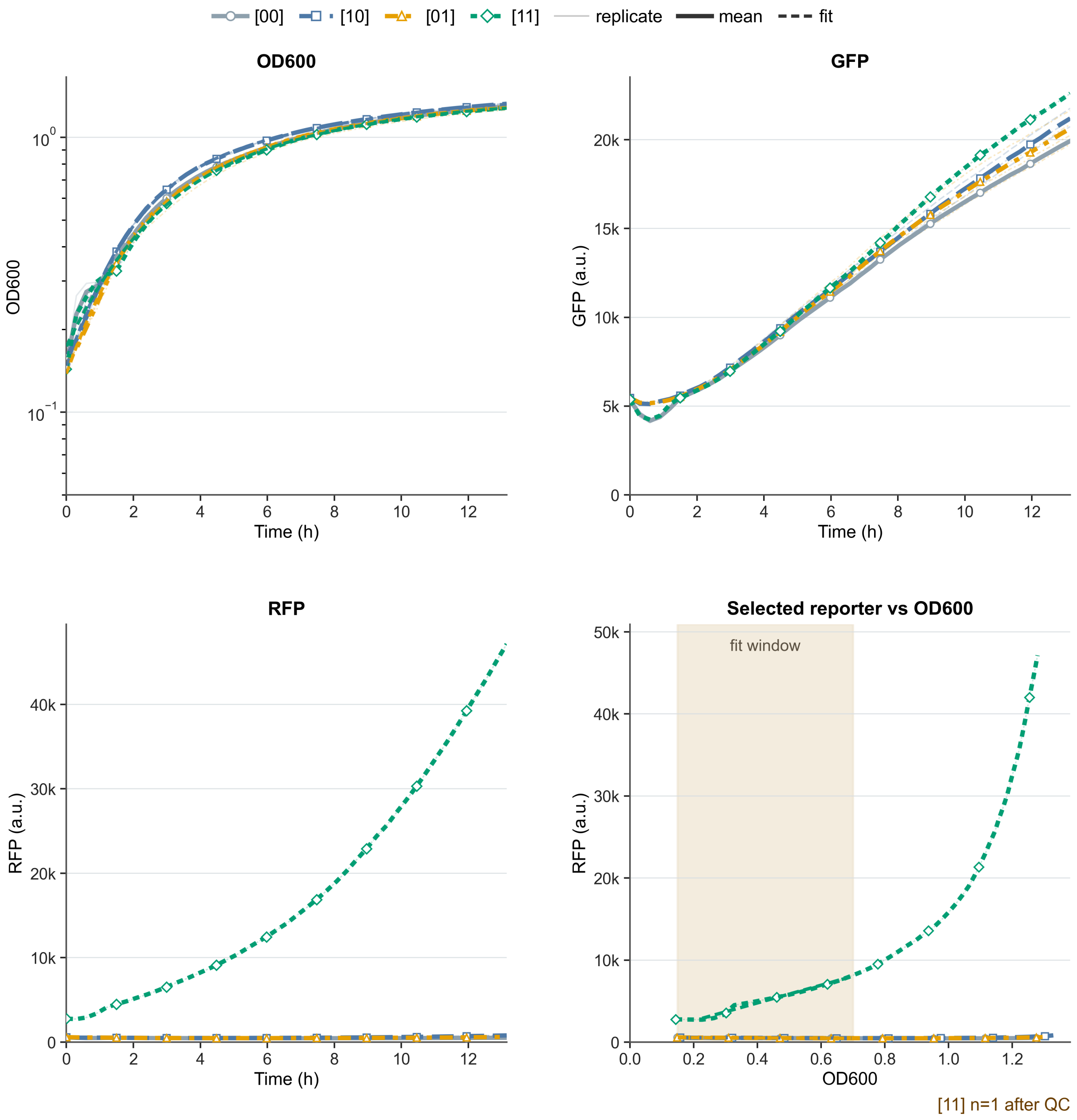

**Supplementary Fig. S11 | Dynamics and fit diagnostics for pVan-Tac.** The four subpanels show $OD_{600}$ versus time with the growth fit, GFP versus time, RFP versus time, and selected-reporter fluorescence versus $OD_{600}$. Selected reporter: RFP. Inducer pair, using the source grid label with units in Supplementary Table S6: Van=21; IPTG=1000. Biological replicates per state: [00] n = 3, [10] n = 3, [01] n = 4, [11] n = 1. The [11] output was high, but this row is N.C. because the [11] state has insufficient biological-replicate support after QC. GFP and RFP are independent reporter outputs, not normalisation controls. The fluorescence-versus-OD diagnostic is the panel used to derive the normalised fluorescence for the selected reporter.

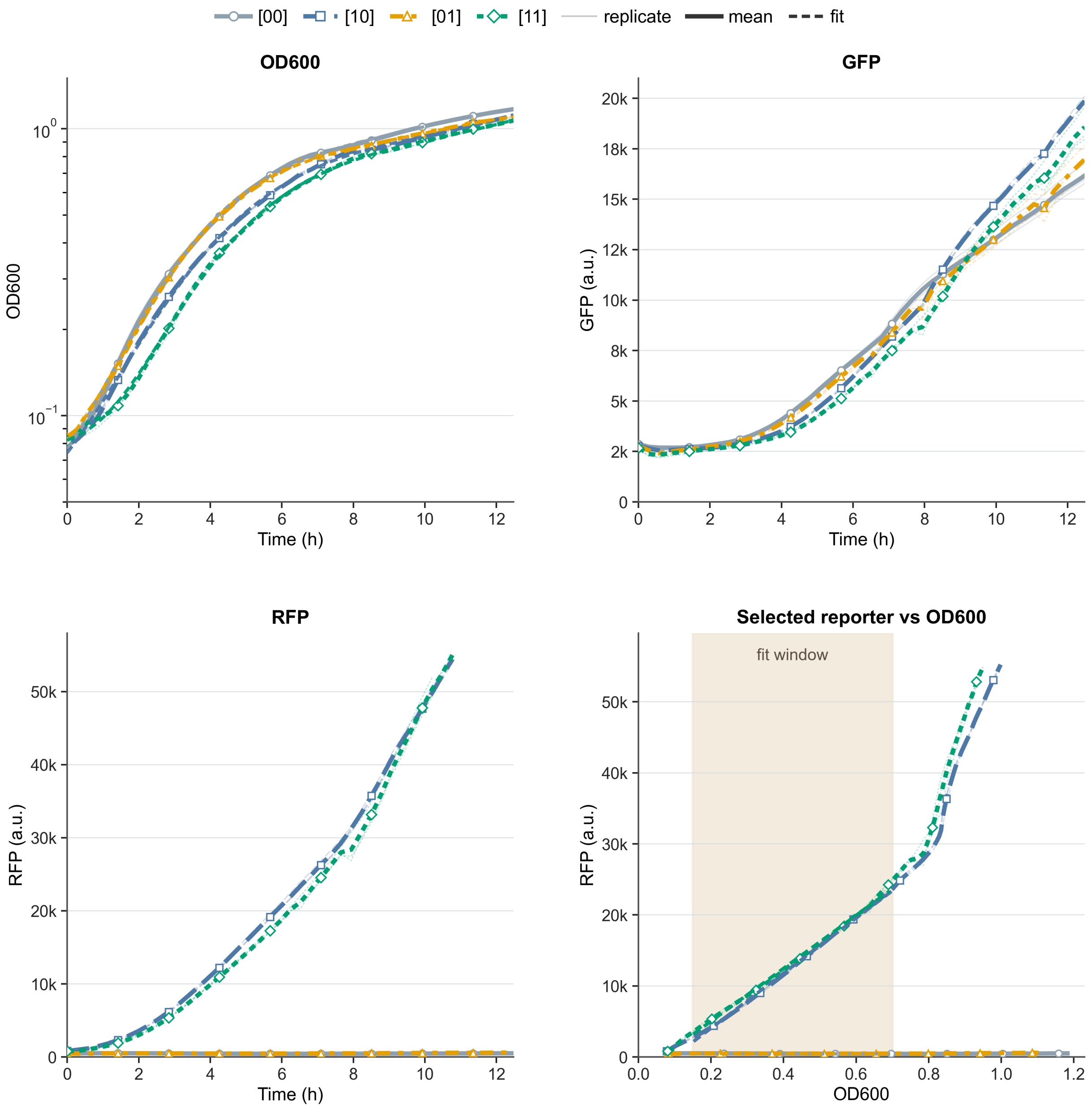

**Supplementary Fig. S12 | Dynamics and fit diagnostics for pVan-Ttg.** The four subpanels show $OD_{600}$ versus time with the growth fit, GFP versus time, RFP versus time, and selected-reporter fluorescence versus $OD_{600}$. Selected reporter: RFP. Inducer pair, using the source grid label with units in Supplementary Table S6: Van=100; Nar=1000. Biological replicates per state: [00] n = 3, [10] n = 3, [01] n = 3, [11] n = 3. GFP and RFP are independent reporter outputs, not normalisation controls. The fluorescence-versus-OD diagnostic is the panel used to derive the normalised fluorescence for the selected reporter.

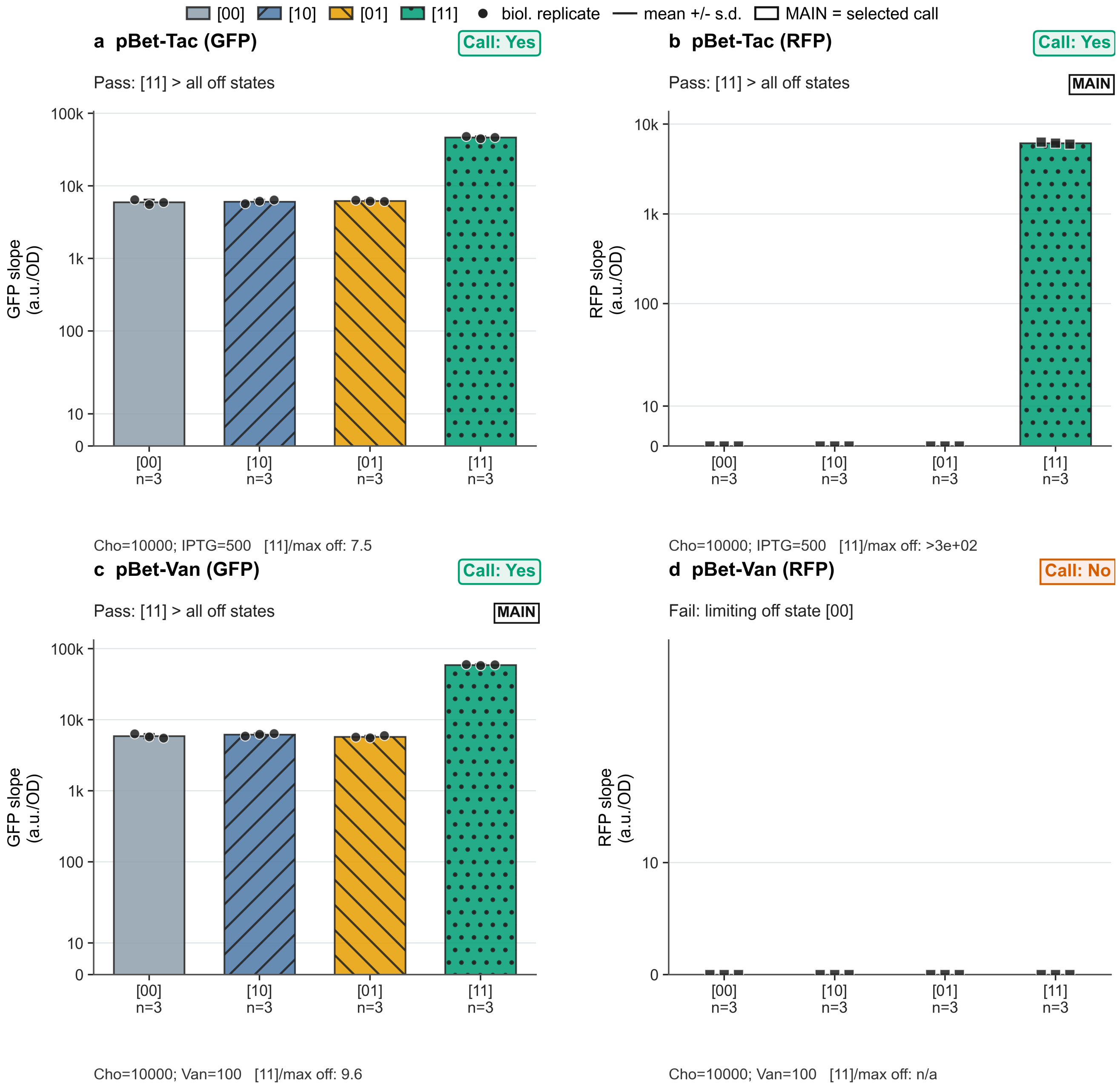

**Supplementary Fig. S13 | Reporter-specific truth tables for pBet-derived architectures matching Supplementary Table S2. a,** pBet-Tac (GFP), **b,** pBet-Tac (RFP), **c,** pBet-Van (GFP), **d,** pBet-Van (RFP). Bars show biological-replicate mean fluorescence-versus-OD slopes, error bars denote s.d., and symbols show individual biological-replicate values. Bars use fixed [00]/[10]/[01]/[11] state colours and hatching. The MAIN badge marks the reporter row used as the architecture-level representative in the main figures. Call badges show Yes for rows where [11] passes the predefined Holm-adjusted Welch operational screen against all off states, No for rows that fail that screen, and N.C. for rows that cannot be classified because one or more required comparisons lack sufficient biological-replicate support. Limiting-state text identifies the off state that restricts separation, while full Holm-adjusted one-sided Welch P values remain in Supplementary Table S2. The y-axis uses symlog scaling with a linear region from 0 to 20 a.u./OD so zero and low values remain visible. GFP and RFP/mCherry are independent output reporters, not normalisation controls.

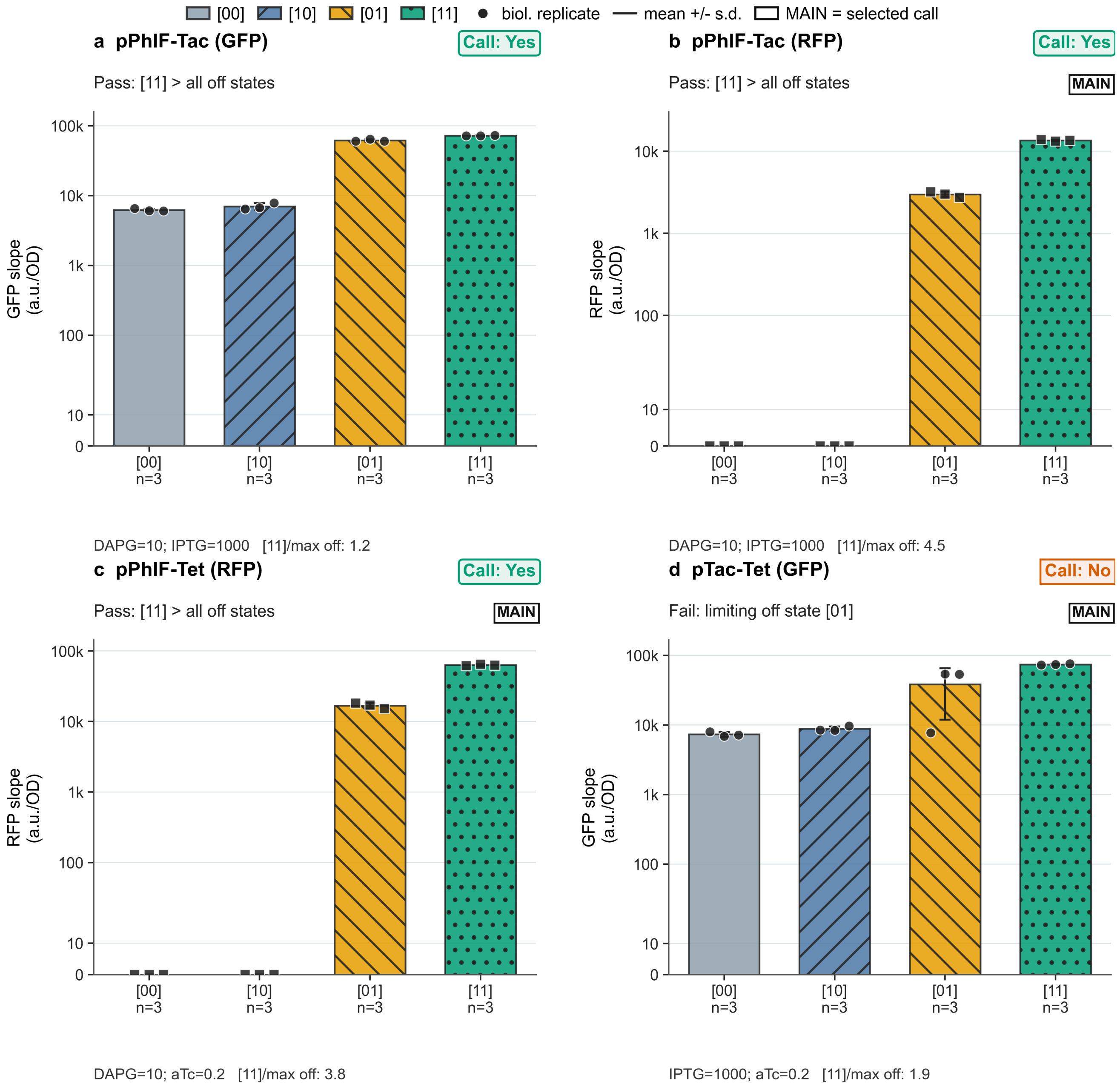

**Supplementary Fig. S14 | Reporter-specific truth tables for pPhlF- and pTac-derived architectures matching Supplementary Table S2. a,** pPhlF-Tac (GFP), **b,** pPhlF-Tac (RFP), **c,** pPhlF-Tet (RFP/mCherry), **d,** pTac-Tet (GFP). pPhlF-Tac(GFP) is an operational pass but not a practical high-utility gate because the [01] state remains close to [11]. Bars show biological-replicate mean fluorescence-versus-OD slopes, error bars denote s.d., and symbols show individual biological-replicate values. Bars use fixed [00]/[10]/[01]/[11] state colours and hatching. The MAIN badge marks the reporter row used as the architecture-level representative in the main figures. Call badges show Yes for rows where [11] passes the predefined Holm-adjusted Welch operational screen against all off states, No for rows that fail that screen, and N.C. for rows that cannot be classified because one or more required comparisons lack sufficient biological-replicate support. Limiting-state text identifies the off state that restricts separation, while full Holm-adjusted one-sided Welch P values remain in Supplementary Table S2. The y-axis uses symlog scaling with a linear region from 0 to 20 a.u./OD so zero and low values remain visible. GFP and RFP/mCherry are independent output reporters, not normalisation controls.

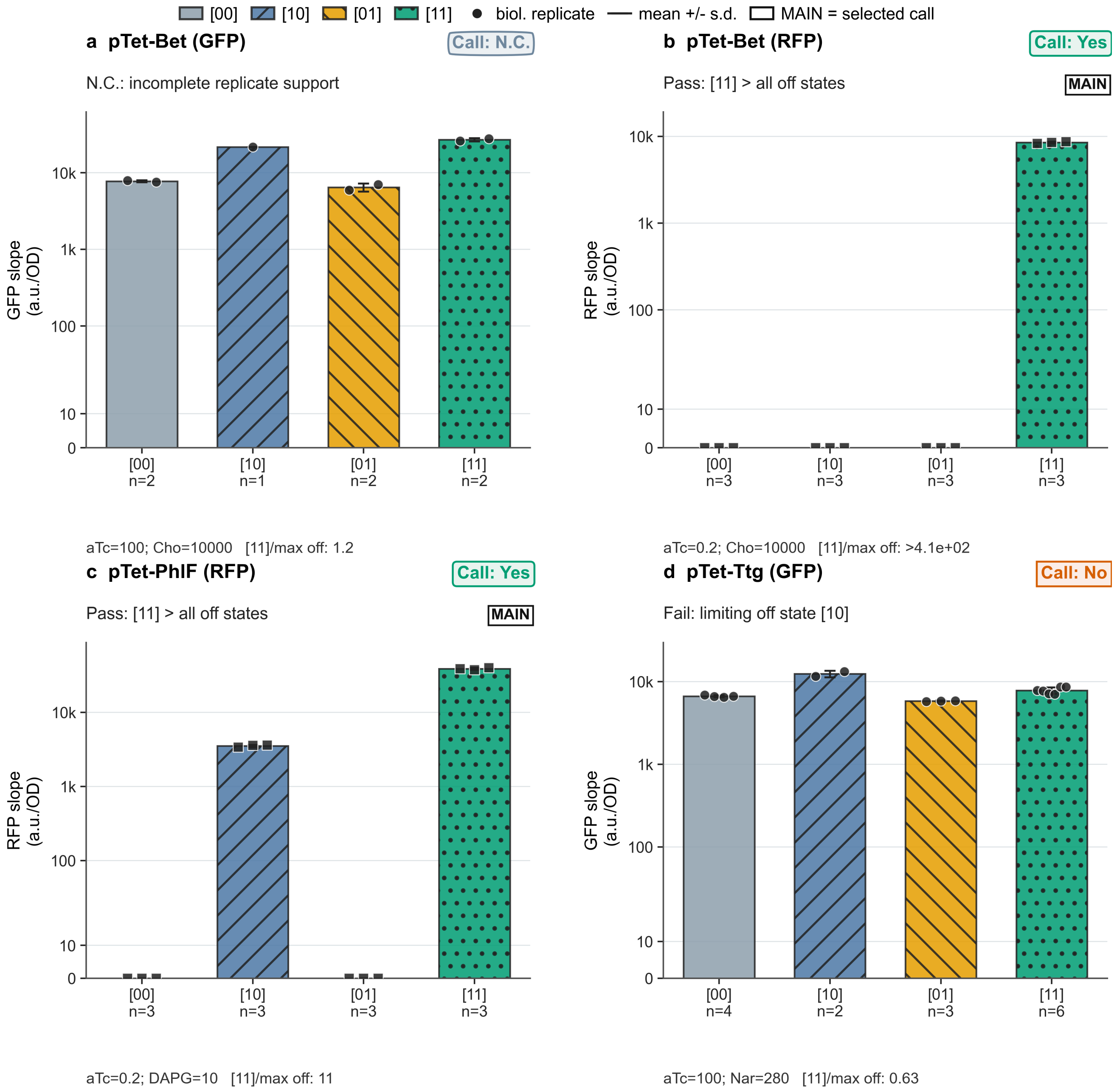

**Supplementary Fig. S15 | Reporter-specific truth tables for pTet-derived architectures matching Supplementary Table S2. a,** pTet-Bet (GFP), **b,** pTet-Bet (RFP/mCherry), **c,** pTet-PhlF (RFP/mCherry), **d,** pTet-Ttg (GFP). Bars show biological-replicate mean fluorescence-versus-OD slopes, error bars denote s.d., and symbols show individual biological-replicate values. Bars use fixed [00]/[10]/[01]/[11] state colours and hatching. The MAIN badge marks the reporter row used as the architecture-level representative in the main figures. Call badges show Yes for rows where [11] passes the predefined Holm-adjusted Welch operational screen against all off states, No for rows that fail that screen, and N.C. for rows that cannot be classified because one or more required comparisons lack sufficient biological-replicate support. Limiting-state text identifies the off state that restricts separation, while full Holm-adjusted one-sided Welch P values remain in Supplementary Table S2. The y-axis uses symlog scaling with a linear region from 0 to 20 a.u./OD so zero and low values remain visible. GFP and RFP/mCherry are independent output reporters, not normalisation controls.

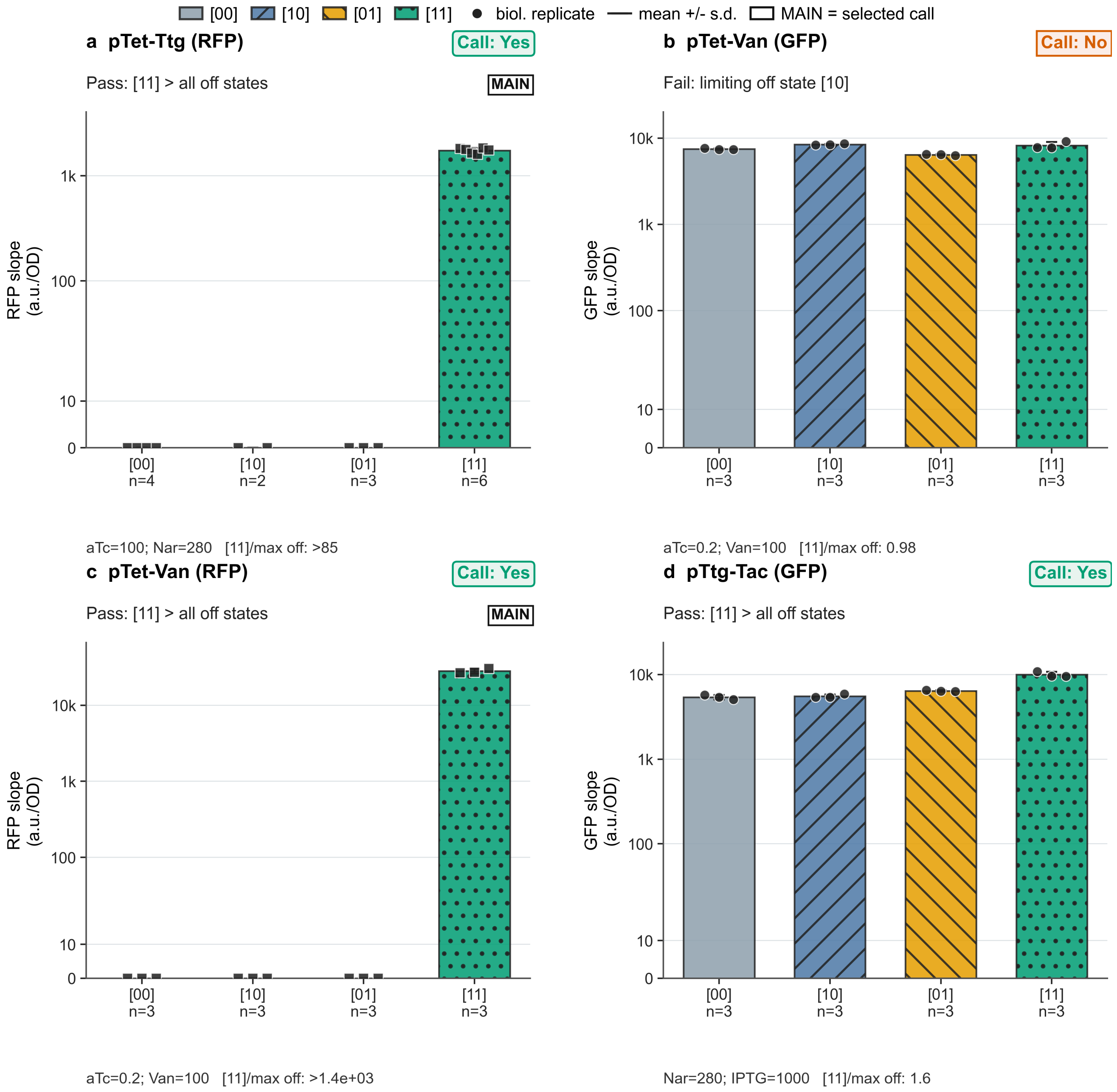
[00] [10] [01] [11] biol. replicate mean +/- s.d. MAIN = selected call
a pTet-Ttg (RFP)
Call: Yes
Pass: [11] > all off states
MAIN
RFP slope (a.u./OD)
1k
100
10
0
[00] n=4
[10] n=2
[01] n=3
[11] n=6
aTc=100; Nar=280 [11]/max off: >85
b pTet-Van (GFP)
Call: No
Fail: limiting off state [10]
GFP slope (a.u./OD)
10k
1k
100
10
0
[00] n=3
[10] n=3
[01] n=3
[11] n=3
aTc=0.2; Van=100 [11]/max off: 0.98
c pTet-Van (RFP)
Call: Yes
Pass: [11] > all off states
MAIN
RFP slope (a.u./OD)
10k
1k
100
10
0
[00] n=3
[10] n=3
[01] n=3
[11] n=3
aTc=0.2; Van=100 [11]/max off: >1.4e+03
d pTtg-Tac (GFP)
Call: Yes
Pass: [11] > all off states
GFP slope (a.u./OD)
10k
1k
100
10
0
[00] n=3
[10] n=3
[01] n=3
[11] n=3
Nar=280; IPTG=1000 [11]/max off: 1.6

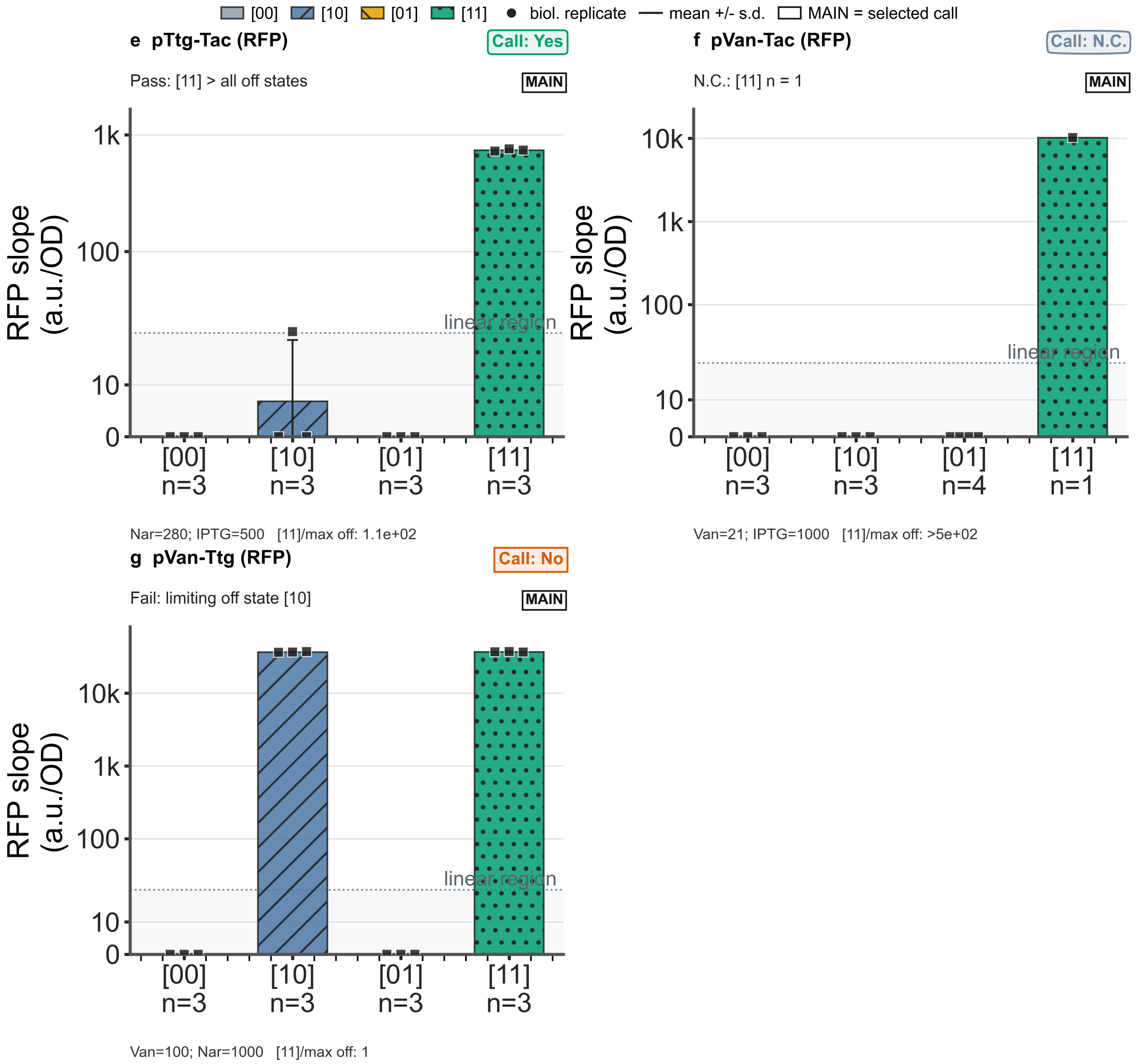

**Supplementary Fig. S16 | Reporter-specific truth tables for additional pTet-, pTtg- and pVan-derived architectures matching Supplementary Table S2. a,** pTet-Ttg (RFP/mCherry), **b,** pTet-Van (GFP), **c,** pTet-Van (RFP/mCherry), **d,** pTtg-Tac (GFP), **e,** pTtg-Tac (RFP/mCherry), **f,** pVan-Tac (RFP/mCherry), **g,** pVan-Ttg (RFP/mCherry). Bars show biological-replicate mean fluorescence-versus-OD slopes, error bars denote s.d., and symbols show individual biological-replicate values. Bars use fixed [00]/[10]/[01]/[11] state colours and hatching. The MAIN badge marks the reporter row used as the architecture-level representative in the main figures. Call badges show Yes for rows where [11] passes the predefined Holm-adjusted Welch operational screen against all off states, No for rows that fail that screen, and N.C. for rows that cannot be classified because one or more required comparisons lack sufficient biological-replicate support. Limiting-state text identifies the off state that restricts separation, while full Holm-adjusted one-sided Welch P values remain in Supplementary Table S2. The y-axis uses symlog scaling with a linear region from 0 to 20 a.u./OD so zero and low values remain visible. GFP and RFP/mCherry are independent output reporters, not normalisation controls. pVan-Tac is N.C. because [11] has n = 1 after QC. pVan-Ttg is a No/failed truth table with [10] as the limiting off state.

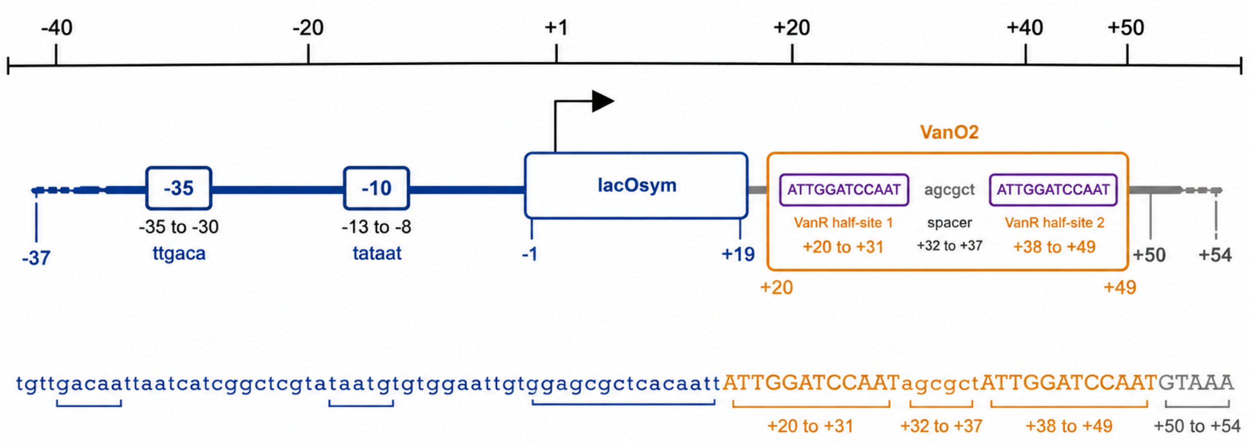

| Feature | Coordinates |
|---|---|
| pTac template | -37 to +19 |
| lacOsym | -1 to +19 |
| VanO2 | +20 to +49 |
| downstream flank | +50 to +54 |

**Supplementary Fig. S17 | Architecture of the experimentally tested pTac-Van combinatorial promoter.** The compact promoter region is drawn to scale and positions are reported relative to the transcription start site (+1). The pTac template, lacOsym site and VanO2 site are coloured consistently with main-text Fig. 1b and Supplementary Table S7. This architecture was experimentally tested and is documented in Supplementary Table S7, but it was not included in the curated main-text architecture set shown in main-text Fig. 1b.

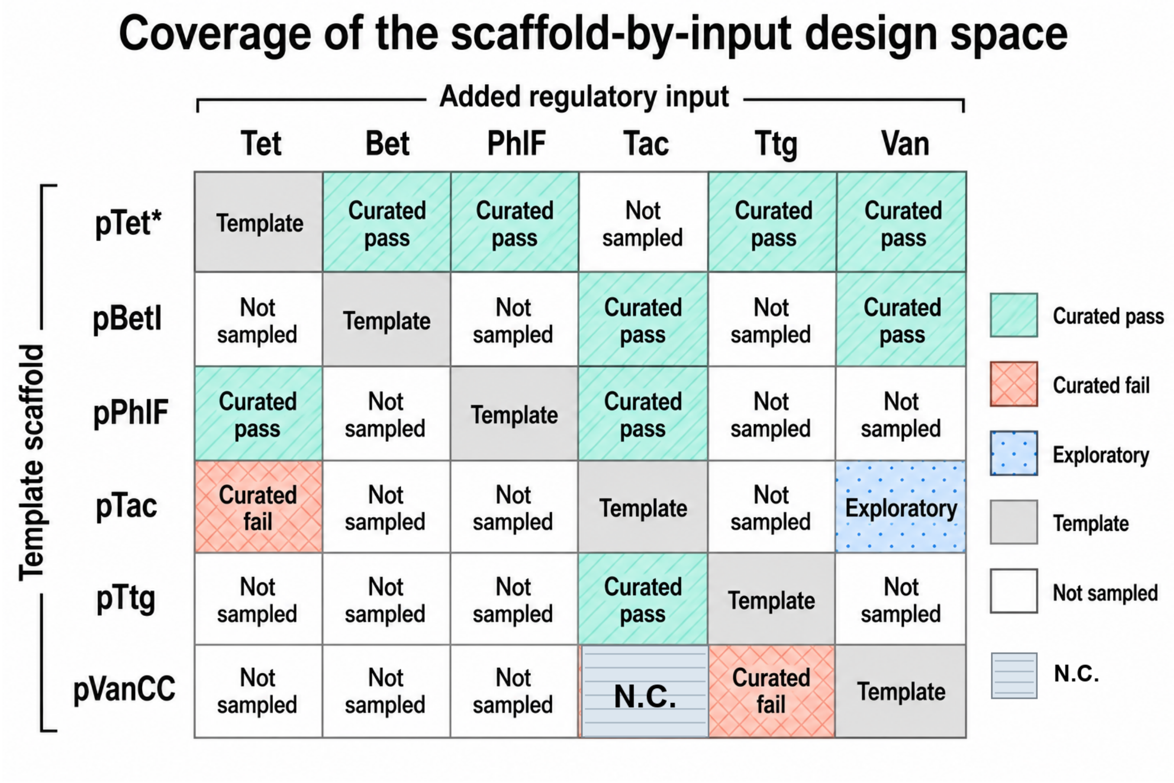

**Supplementary Fig. S18 | Coverage of the scaffold-by-input design space used in this study.** The matrix uses colour and text labels to distinguish curated architectures that passed the operational screen (green), curated architectures that failed (red), the not-classifiable pVan-Tac case (grey-blue N.C.), the exploratory pTac-Van construct (blue), template-only entries (grey), and scaffold-input combinations that were not sampled in the present benchmark (white).

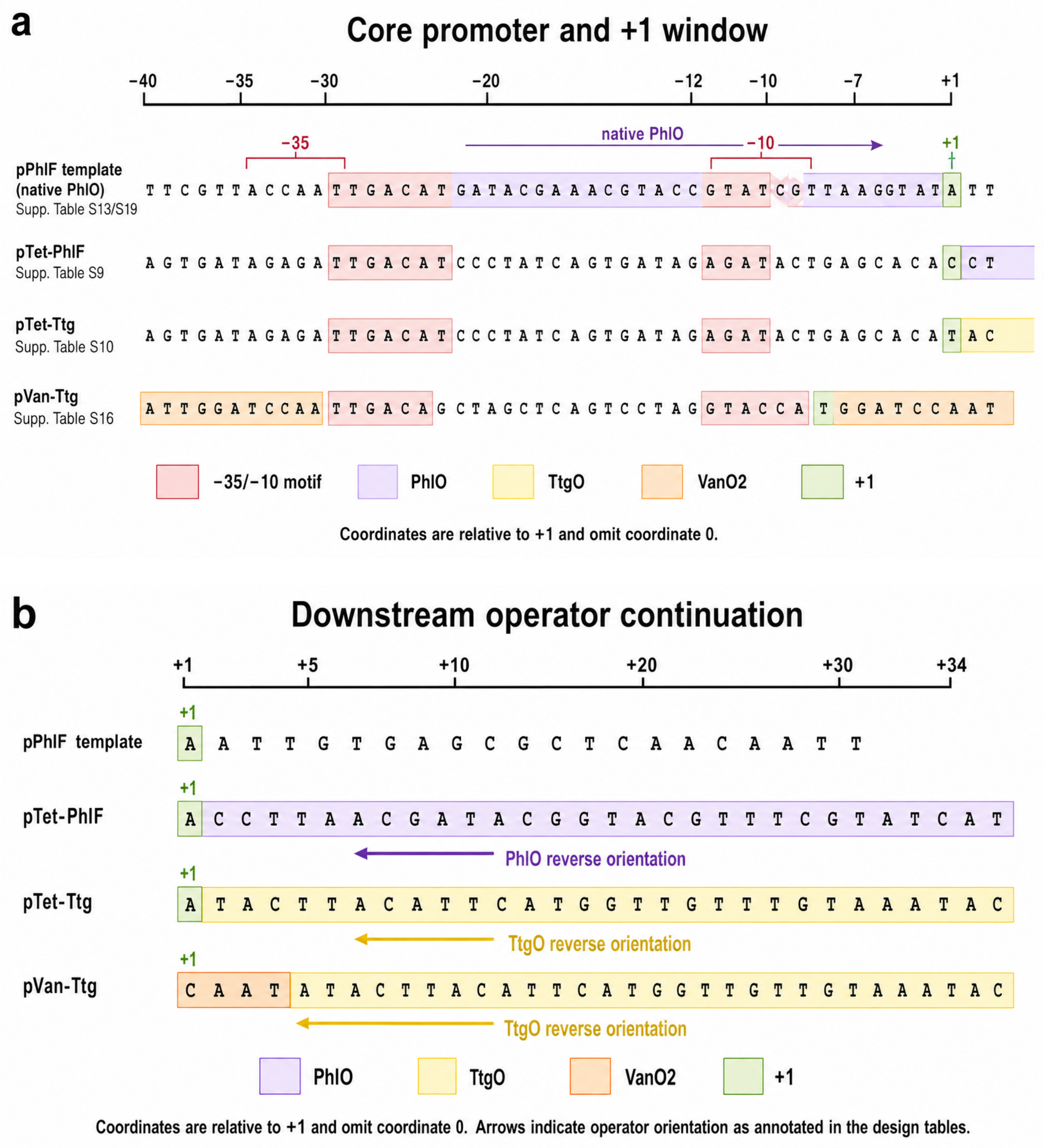

**Supplementary Fig. S19 | Sequence alignment around the -10 region and +1 site in reverse-operator examples. a,** Core-promoter/+1 window. **b,** Downstream operator continuation. Rows show pPhlF template/native PhlO, pTet-PhlF, pTet-Ttg and pVan-Ttg. DNA bases are monospaced. Coordinates are relative to +1 and omit coordinate 0. Colours mark PhlO (purple), TtgO (yellow), VanO2 (orange), annotated -35/-10 motifs (pink) and +1 (green). Reverse orientation is indicated for inserted PhlO and TtgO. The figure illustrates sequence-aware placement used to avoid promoter-like motif arrangements in compact downstream-operator designs. It does not demonstrate that orientation caused promoter behaviour.

### Supplementary Tables

| Distributed file | Contents |
|---|---|
| source_data.xlsx | Source-data workbook distributed with the submission. It contains source data for retained main and supplementary figures and tables. |
| code_ocean_scripts.zip | Python scripts for reproducing the processed analyses and regenerated quantitative charts from the submitted source-data workbook. |
| snapgene_files.zip | SnapGene DNA files for the promoter-reporter plasmid constructs. |

**Supplementary Table S1 | Distributed source-data and reproducibility files.**

**a**

| Promoter | Reporter | Template | Added input | Template inducer | Added inducer | Inducer pair | Selected main call | Operational call | Engineering Utility |
|---|---|---|---|---|---|---|---|---|---|
| pBet-Tac | GFP | pBetI | Tac | Cho | IPTG | Cho=10000; IPTG=500 | No | Yes | True |
| pBet-Tac | RFP | pBetI | Tac | Cho | IPTG | Cho=10000; IPTG=500 | Yes | Yes | True |
| pBet-Van | GFP | pBetI | Van | Cho | Van | Cho=10000; Van=100 | Yes | Yes | True |
| pBet-Van | RFP | pBetI | Van | Cho | Van | Cho=10000; Van=100 | No | No | False |
| pPhlF-Tac | GFP | pPhlF | Tac | DAPG | IPTG | DAPG=10; IPTG=1000 | No | Yes | False |
| pPhlF-Tac | RFP | pPhlF | Tac | DAPG | IPTG | DAPG=10; IPTG=1000 | Yes | Yes | True |
| pPhlF-Tet | RFP | pPhlF | Tet | DAPG | aTc | DAPG=10; aTc=0.2 | Yes | Yes | True |
| pTac-Tet | GFP | pTac | Tet | IPTG | aTc | IPTG=1000; aTc=0.2 | Yes | No | False |
| pTet-Bet | GFP | pTet* | Bet | aTc | Cho | aTc=100; Cho=10000 | No | N.C. | False |
| pTet-Bet | RFP | pTet* | Bet | aTc | Cho | aTc=0.2; Cho=10000 | Yes | Yes | True |
| pTet-PhlF | RFP | pTet* | PhlF | aTc | DAPG | aTc=0.2; DAPG=10 | Yes | Yes | True |
| pTet-Ttg | GFP | pTet* | Ttg | aTc | Nar | aTc=100; Nar=280 | No | No | False |
| pTet-Ttg | RFP | pTet* | Ttg | aTc | Nar | aTc=100; Nar=280 | Yes | Yes | True |
| pTet-Van | GFP | pTet* | Van | aTc | Van | aTc=0.2; Van=100 | No | No | False |
| pTet-Van | RFP | pTet* | Van | aTc | Van | aTc=0.2; Van=100 | Yes | Yes | True |
| pTtg-Tac | GFP | pTtg | Tac | Nar | IPTG | Nar=280; IPTG=1000 | No | Yes | False |
| pTtg-Tac | RFP | pTtg | Tac | Nar | IPTG | Nar=280; IPTG=500 | Yes | Yes | True |
| pVan-Tac | RFP | pVanCC | Tac | Van | IPTG | Van=21; IPTG=1000 | Yes | N.C. | False |
| pVan-Ttg | RFP | pVanCC | Ttg | Van | Nar | Van=100; Nar=1000 | Yes | No | False |

**b**

| Promoter | Reporter | [00] mean | [10] mean | [01] mean | [11] mean |
|---|---|---|---|---|---|
| pBet-Tac | GFP | 5973 (6472) | 6075 (6858) | 6206 (6398) | 46516 (5710) |
| pBet-Tac | RFP | 0.000 (0.000) | 0.000 (0.000) | 0.000 (0.000) | 6139 (0.000) |
| pBet-Van | GFP | 5876 | 6180 | 5741 | 59190 |
| pBet-Van | RFP | 0.000 | 0.000 | 0.000 | 0.000 |
| pPhlF-Tac | GFP | 6200 (5807) | 6986 (5361) | 61486 (5746) | 71953 (7336) |
| pPhlF-Tac | RFP | 0.000 (0.000) | 0.000 (0.000) | 2988 (0.000) | 13564 (0.000) |
| pPhlF-Tet | RFP | 0.000 | 0.000 | 16800 | 63107 |

| | | | | | |
|---|---|---|---|---|---|
| pTac-Tet | GFP | 7339 | 8818 | 38428 | 74064 |
| pTet-Bet | GFP | 7753 (5253) | 21701 (5957) | 6455 (6100) | 26924 (6583) |
| pTet-Bet | RFP | 0.000 (0.000) | 0.000 (1063) | 0.000 (0.000) | 8479 (2188) |
| pTet-PhlF | RFP | 0.000 | 3517 | 0.000 | 38690 |
| pTet-Ttg | GFP | 6649 | 12377 | 5806 | 7813 |
| pTet-Ttg | RFP | 0.000 | 0.000 | 0.000 | 1740 |
| pTet-Van | GFP | 7451 | 8425 | 6395 | 8216 |
| pTet-Van | RFP | 0.000 | 0.000 | 0.000 | 28030 |
| pTtg-Tac | GFP | 5386 (6858) | 5547 (6899) | 6389 (6304) | 10002 (6092) |
| pTtg-Tac | RFP | 0.000 (0.000) | 6.831 (0.000) | 0.000 (0.000) | 739.87 (0.000) |
| pVan-Tac | RFP | 0.000 | 0.000 | 0.000 | 10256 |
| pVan-Ttg | RFP | 0.000 | 36787 | 0.000 | 36976 |

**c**

| Promoter | Reporter | [00] sd | [10] sd | [01] sd | [11] sd | n [00] | n [10] | n [01] | n [11] |
|---|---|---|---|---|---|---|---|---|---|
| pBet-Tac | GFP | 465.69 | 357.55 | 141.36 | 1651 | 3 | 3 | 3 | 3 |
| pBet-Tac | RFP | 0.000 | 0.000 | 0.000 | 143.08 | 3 | 3 | 3 | 3 |
| pBet-Van | GFP | 406.05 | 252.59 | 217.03 | 1099 | 3 | 3 | 3 | 3 |
| pBet-Van | RFP | 0.000 | 0.000 | 0.000 | 0.000 | 3 | 3 | 3 | 3 |
| pPhlF-Tac | GFP | 279.05 | 761.17 | 2243 | 678.67 | 3 | 3 | 3 | 3 |
| pPhlF-Tac | RFP | 0.000 | 0.000 | 228.75 | 258.88 | 3 | 3 | 3 | 3 |
| pPhlF-Tet | RFP | 0.000 | 0.000 | 1509 | 1653 | 3 | 3 | 3 | 3 |
| pTac-Tet | GFP | 585.32 | 691.41 | 26605 | 1406 | 3 | 3 | 3 | 3 |
| pTet-Bet | GFP | 245.17 | 0.000 | 761.93 | 1297 | 2 | 1 | 2 | 2 |
| pTet-Bet | RFP | 0.000 | 0.000 | 0.000 | 204.60 | 3 | 3 | 3 | 3 |
| pTet-PhlF | RFP | 0.000 | 122.34 | 0.000 | 1190 | 3 | 3 | 3 | 3 |
| pTet-Ttg | GFP | 189.48 | 1142 | 74.51 | 679.44 | 4 | 2 | 3 | 6 |
| pTet-Ttg | RFP | 0.000 | 0.000 | 0.000 | 94.78 | 4 | 2 | 3 | 6 |
| pTet-Van | GFP | 129.07 | 139.60 | 99.68 | 785.88 | 3 | 3 | 3 | 3 |
| pTet-Van | RFP | 0.000 | 0.000 | 0.000 | 2023 | 3 | 3 | 3 | 3 |
| pTtg-Tac | GFP | 334.90 | 280.64 | 129.11 | 754.13 | 3 | 3 | 3 | 3 |
| pTtg-Tac | RFP | 0.000 | 11.83 | 0.000 | 15.17 | 3 | 3 | 3 | 3 |
| pVan-Tac | RFP | 0.000 | 0.000 | 0.000 | 0.000 | 3 | 3 | 4 | 1 |
| pVan-Ttg | RFP | 0.000 | 509.59 | 0.000 | 236.54 | 3 | 3 | 3 | 3 |

**d**

| Promoter | Reporter | Max off | Highest off | [11]/max off | Lower bound | Holm P [11]>[00] | Holm P [11]>[10] | Holm P [11]>[01] |
|---|---|---|---|---|---|---|---|---|
| pBet-Tac | GFP | 6206 | [01] | 7.495 | No | <0.001 | <0.001 | <0.001 |
| pBet-Tac | RFP | 0.000 | [00] | >299.55 | Yes | <0.001 | <0.001 | <0.001 |
| pBet-Van | GFP | 6180 | [10] | 9.578 | No | <0.001 | <0.001 | <0.001 |
| pBet-Van | RFP | 0.000 | [00] | | Yes | 1.000 | 1.000 | 1.000 |
| pPhlF-Tac | GFP | 61486 | [01] | 1.170 | No | <0.001 | <0.001 | 0.005 |
| pPhlF-Tac | RFP | 2988 | [01] | 4.539 | No | <0.001 | <0.001 | <0.001 |

| | | | | | | | | |
|---|---|---|---|---|---|---|---|---|
| pPhlF-Tet | RFP | 16800 | [01] | 3.756 | No | <0.001 | <0.001 | <0.001 |
| pTac-Tet | GFP | 38428 | [01] | 1.927 | No | <0.001 | <0.001 | 0.073 |
| pTet-Bet | GFP | 21701 | [10] | 1.241 | No | | | |
| pTet-Bet | RFP | 0.000 | [00] | >413.74 | Yes | <0.001 | <0.001 | <0.001 |
| pTet-PhlF | RFP | 3517 | [10] | 11.00 | No | <0.001 | <0.001 | <0.001 |
| pTet-Ttg | GFP | 12377 | [10] | 0.631 | No | 0.007 | 0.959 | 0.001 |
| pTet-Ttg | RFP | 0.000 | [00] | >84.88 | Yes | <0.001 | <0.001 | <0.001 |
| pTet-Van | GFP | 8425 | [10] | 0.975 | No | 0.232 | 0.654 | 0.082 |
| pTet-Van | RFP | 0.000 | [00] | >1368 | Yes | 0.003 | 0.003 | 0.003 |
| pTtg-Tac | GFP | 6389 | [01] | 1.566 | No | 0.005 | 0.005 | 0.006 |
| pTtg-Tac | RFP | 6.831 | [10] | 108.30 | No | <0.001 | <0.001 | <0.001 |
| pVan-Tac | RFP | 0.000 | [00] | >500.44 | Yes | | | |
| pVan-Ttg | RFP | 36787 | [10] | 1.005 | No | <0.001 | 0.302 | <0.001 |

**Supplementary Table S2 | Statistics for all 19 curated two-input combinatorial promoter-reporter truth tables.** The table reports all curated reporter-specific truth tables rather than only the selected architecture-level rows. **a,** Construct identity, operational classification and engineering-utility flag. **b,** State means; parenthetical values report same-channel alternate-reporter diagnostics where an independent matched alternate reporter context is available. **c,** State standard deviations and biological replicate counts. **d,** Off-state ratio and Holm-adjusted comparisons. The selected main call column identifies the reporter used for the main architecture-level call. Ratios with non-positive maximum off-state means are reported as lower bounds rather than infinity. N.C. or not_classifiable indicates that one or more required state comparisons lacked sufficient biological-replicate support.

| Promoter | Reporter | Reason if not selected |
|---|---|---|
| pBet-Tac | GFP | not selected as architecture-level representative |
| pBet-Tac | RFP | |
| pBet-Van | GFP | |
| pBet-Van | RFP | not selected as architecture-level representative |
| pPhlF-Tac | GFP | not selected as architecture-level representative |
| pPhlF-Tac | RFP | |
| pPhlF-Tet | RFP | |
| pTac-Tet | GFP | |
| pTet-Bet | GFP | not selected as architecture-level representative |
| pTet-Bet | RFP | |
| pTet-PhlF | RFP | |
| pTet-Ttg | GFP | not selected as architecture-level representative |
| pTet-Ttg | RFP | |
| pTet-Van | GFP | not selected as architecture-level representative |
| pTet-Van | RFP | |
| pTtg-Tac | GFP | not selected as architecture-level representative |
| pTtg-Tac | RFP | |
| pVan-Tac | RFP | |
| pVan-Ttg | RFP | |

**Supplementary Table S3 | Selection notes and provenance.**

| # | Primer name | Sequence |
|---|---|---|
| 1 | AND pTet-Ttg (aTc+NAR)151 FW | agcacATACTTACATTCATGGTTGTTTGTAAATACACagctgtcaccggatgtgctttc |
|  | AND pTet-Ttg (aTc+NAR)152 RW | gctGTGTATTTACAAACAACCATGAATGTAAGTATgtgctcagtatctctatcactgatagggatg |
| 2 | AND pVan-Ttg (Van+NAR) 155 FW | ATACTTACATTCATGGTTGTTTGTAAATACTGTAAACagctgtcaccggatgtgctttc |
|  | AND pVan-Ttg (Van+NAR) 156 RW | TTTACATTGGATCCAATagcgctATTGGATCCAATaattgtgagcgctcacaattccacac |
| 3 | AND pBetI-Tac (cho-IPTG)161) FW | ataatgctagcaattgtgagcgctcacaattAAGATCGGGTGTAAACagctgtcac |
|  | AND pBetI-Tac (cho-IPTG)162) FW | CCCGATCTTaattgtgagcgctcacaattgctagcattatattgaacgtccaatcaattg |
| 4 | AND pPhlF-Tac(DAPG- IPTG)165 FW | gtatcgttaaggtaattgtgagcgctcacaattTCGGGTGTAAACagctgtcaccg |
|  | AND pPhlF-Tac(DAPG- IPTG)166 RW | CACCCGAaattgtgagcgctcacaattaccttaacgatacggtacgtttcgtatca |
| 5 | AND pTtg-pTac (NAR-IPTG)167 FW | tatattccttagcaaaattgtgagcgctcacaattGGGTGTAAACagctgtcaccggat |
|  | AND pTtg-pTac (NAR-IPTG)168 FW | CACCCaattgtgagcgctcacaattttgctaaggaatatacttacattcatggttgtttg |
| 6 | AND pVan-Tac (Van-IPTG)169 FW | tggatccaataattgtgagcgctcacaattTAAGATCGGGTGTAAACagctgtcac |
|  | AND pVan-Tac (Van-IPTG)170 FW | CCCGATCTTAaattgtgagcgctcacaattattggatccaatggtacctaggactg |
| 7 | AND pTet -Bet (aTc-Cho)173 FW | atccggtgacagctGTTTACACCCattgattggacgttcaatataagtgctcagtatc |
|  | AND pTet- Bet (aTc-Cho)174 RW | TACACCCattgattggacgttcaatataagtgctcagtatctctatcactgatagggat |
| 8 | AND pTet-PhiF (aTc-DAPG)175 FW | ctgagcacaccttaacgatacggtacgtttcgtatcatACagctgtcaccggatgtgctttc |
|  | AND pTet-PhiF (aTc-DAPG)176 RW | GTatgatacgaaacgtaccgtatcgttaaggtgtgctcagtatctctatcactgatagggatg |
| 9 | AND pTet- Van (aTc-VAN)177 FW | gagcacATTGGATCCAATagcgctATTGGATCCAATACagctgtcaccggatgtgctttc |
|  | AND pTet- Van (aTc-VAN)178 RW | ctGTATTGGATCCAATagcgctATTGGATCCAATgtgctcagtatctctatcactgatagggatg |
| 10 | AND pTac-Van (IPTG+VAN)179 FW | acaattATTGGATCCAATagcgctATTGGATCCAATGTAAACagctgtcaccggatgtgc |
|  | AND pTac-Van (IPTG+VAN)180 RW | TTACATTGGATCCAATagcgctATTGGATCCAATaattgtgagcgctcacaattccacac |
| 11 | AND pTac-Tet (IPTG-aTc)194 FW | ctatcagtgatagagaTTGTGAGCGGATAACAAagctgtcaccggatgtgctttcc |
|  | AND pTac-Tet (IPTG-aTc)194 FW | gacagctTTGTTATCCGCTCACAAtctctatcactgatagggaaattgtgagcgc |

**Supplementary Table S4 | Primers used in this work.**

| Promoter | Sequence |
|---|---|
| pTac-Tet | tgttgacaattaatcatcggctcgtataatgtgtggaattgtgagcgctcacaatttccctatcagtgatag agaTAAGATCGGGTGTAAA |
| pTac-Van | tgttgacaattaatcatcggctcgtataatgtgtggaattgtgagcgctcacaattATTGGATCCAATagcg ctATTGGATCCAATGTAAA |
| pTet-Van | ttttcagcaggacgcactgacctccctatcagtgatagagattgacatccctatcagtgatagagatactga gcacATTGGATCCAATagcgctATTGGATCCAATA |
| pTet-Bet | ttttcagcaggacgcactgacctccctatcagtgatagagattgacatccctatcagtgatagagatactga gcacttatattgaacgtccaatcaatGGGTGTAAA |
| pTet-Ttg | ttttcagcaggacgcactgacctccctatcagtgatagagattgacatccctatcagtgatagagatactga gcacATACTTACATTCATGGTTGTTTGTAAATAC |
| pTet-PhlF | ttttcagcaggacgcactgacctccctatcagtgatagagattgacatccctatcagtgatagagatactga gcacaccttaacgatacggtacgtttcgtatcatA |
| pVan-Ttg | ttggatccaattgacagctagctcagtcctaggtaccattggatccaatATACTTACATTCATGGTTGTTTG TAAATACTGTAAA |
| pVan-Tac | attggatccaattgacagctagctcagtcctaggtaccattggatccaataattgtgagcgctcacaattTA AGATCGGGTGTAAA |
| pBet-Tac | agcgcgggtgagagggattcgttaccaatagacaattgattggacgttcaatataatgctagcaattgtgag cgctcacaattAAGATCGGGTGTAAA |
| pTtg-Tac | cacccagcagtatttacaaacaaccatgaatgtaagtatattccttagcaaaattgtgagcgctcacaattG GGTGTAAA |
| pPhlF-Tac | cgacgtacggtggaatctgattcgttaccaattgacatgatacgaaacgtaccgtatcgttaaggtaattgt gagcgctcacaattTCGGGTGTAAA |

**Supplementary Table S5 | Promoter sequences.**

a

| Inducer | Full name | Unit used in concentration table |
|---|---|---|
| DAPG | 2,4-diacetylphloroglucinol | μM |
| IPTG | isopropyl β-D-1-thiogalactopyranoside | μM |
| aTc | anhydrotetracycline hydrochloride | 0.2 μM or 100 ng mL−1, matching the source grid label |
| Cho | choline chloride | μM |
| Nar | naringenin | μM |
| Van | vanillic acid | μM |

b

| Promoter-reporter and source grid label | [10] first/template input ON | [01] second/added input ON | [11] both inputs ON |
|---|---|---|---|
| pBet-Tac (GFP) Cho=10000; IPTG=500 | pBetI input; Cho 10000 μM | Tac input; IPTG 500 μM | Cho 10000 μM + IPTG 500 μM |
| pBet-Tac (RFP) Cho=10000; IPTG=500 | pBetI input; Cho 10000 μM | Tac input; IPTG 500 μM | Cho 10000 μM + IPTG 500 μM |
| pBet-Van (GFP) Cho=10000; Van=100 | pBetI input; Cho 10000 μM | Van input; Van 100 μM | Cho 10000 μM + Van 100 μM |
| pBet-Van (RFP) Cho=10000; Van=100 | pBetI input; Cho 10000 μM | Van input; Van 100 μM | Cho 10000 μM + Van 100 μM |
| pPhlF-Tac (GFP) DAPG=10; IPTG=1000 | pPhlF input; DAPG 10 μM | Tac input; IPTG 1000 μM | DAPG 10 μM + IPTG 1000 μM |
| pPhlF-Tac (RFP) DAPG=10; IPTG=1000 | pPhlF input; DAPG 10 μM | Tac input; IPTG 1000 μM | DAPG 10 μM + IPTG 1000 μM |
| pPhlF-Tet (RFP) DAPG=10; aTc=0.2 | pPhlF input; DAPG 10 μM | Tet input; aTc 0.2 μM | DAPG 10 μM + aTc 0.2 μM |
| pTac-Tet (GFP) IPTG=1000; aTc=0.2 | pTac input; IPTG 1000 μM | Tet input; aTc 0.2 μM | IPTG 1000 μM + aTc 0.2 μM |
| pTet-Bet (GFP) aTc=100; Cho=10000 | pTet* input; aTc 100 ng mL−1 | Bet input; Cho 10000 μM | aTc 100 ng mL−1 + Cho 10000 μM |

c

| Promoter-reporter and source grid label | [10] first/template input ON | [01] second/added input ON | [11] both inputs ON |
|---|---|---|---|
| pTet-Bet (RFP) aTc=0.2; Cho=10000 | pTet* input; aTc 0.2 μM | Bet input; Cho 10000 μM | aTc 0.2 μM + Cho 10000 μM |
| pTet-PhlF (RFP) aTc=0.2; DAPG=10 | pTet* input; aTc 0.2 μM | PhlF input; DAPG 10 μM | aTc 0.2 μM + DAPG 10 μM |
| pTet-Ttg (GFP) aTc=100; Nar=280 | pTet* input; aTc 100 ng mL−1 | Ttg input; Nar 280 μM | aTc 100 ng mL−1 + Nar 280 μM |
| pTet-Ttg (RFP) aTc=100; Nar=280 | pTet* input; aTc 100 ng mL−1 | Ttg input; Nar 280 μM | aTc 100 ng mL−1 + Nar 280 μM |
| pTet-Van (GFP) aTc=0.2; Van=100 | pTet* input; aTc 0.2 μM | Van input; Van 100 μM | aTc 0.2 μM + Van 100 μM |
| pTet-Van (RFP) aTc=0.2; Van=100 | pTet* input; aTc 0.2 μM | Van input; Van 100 μM | aTc 0.2 μM + Van 100 μM |
| pTtg-Tac (GFP) Nar=280; IPTG=1000 | pTtg input; Nar 280 μM | Tac input; IPTG 1000 μM | Nar 280 μM + IPTG 1000 μM |
| pTtg-Tac (RFP) Nar=280; IPTG=500 | pTtg input; Nar 280 μM | Tac input; IPTG 500 μM | Nar 280 μM + IPTG 500 μM |
| pVan-Tac (RFP) Van=21; IPTG=1000 | pVanCC input; Van 21 μM | Tac input; IPTG 1000 μM | Van 21 μM + IPTG 1000 μM |

| Promoter-reporter and source grid label | [10] first/template input ON | [01] second/added input ON | [11] both inputs ON |
|---|---|---|---|
| pVan-Ttg (RFP) Van=100; Nar=1000 | pVanCC input; Van 100 μM | Ttg input; Nar 1000 μM | Van 100 μM + Nar 1000 μM |

**Supplementary Table S6 | Selected inducer concentrations and truth-table digit mapping. a,** Inducer identity and units. **b,** Selected non-zero working concentrations for the analysed truth-table grids. **c,** Continuation of selected non-zero working concentrations for the analysed truth-table grids. Concentrations are the selected non-zero working concentrations for the analysed four-state grid, not stock concentrations. The source grid label is preserved exactly as used in Supplementary Table S2 and the source workbooks. State [00] contains neither inducer, [10] contains only the first/template-input inducer, [01] contains only the second/added-input inducer, and [11] contains both. For aTc, the source workbooks use two notations: rows labelled aTc=0.2 are reported as 0.2 μM, whereas rows labelled aTc=100 are reported as 100 ng $mL^{-1}$; this table keeps those source-specific unit conventions explicit.

| Promoter Name | Template Region (-35/-10) | Added Operator Position | Full Sequence (5' to 3') |
|---|---|---|---|
| pTac-Van | **pTac template**: -37 to +19; -35: -35 to -30; -10: -13 to -8; +1: +1<br>Underlined overlap: **lacOsym** -1 to +19 within pTac template; underlined bases use the larger template-feature colour. | **VanO2**: +20 to +49 | tgttgacaattaatcatcggctcgtataatgtgtggaattgtgagcgctcacaattATTGGATCCAATagcgctATTGGATCCAAT |
| pTet-Bet | **pTet* template**: -76 to -1; -35: -35 to -30; -10: -12 to -7<br>Underlined overlap: **tetO2** -54 to -36 within pTet* template; underlined bases use the larger template-feature colour.<br>Underlined overlap: **tetO2** -29 to -11 within pTet* template; underlined bases use the larger template-feature colour. | **BetO**: +1 to +22 | ttttcagcaggacgcactgacctccctatcagtgatagagattgacatccctatcagtgatagagatactgagcacttatattgaacgtccaatcaat |
| pTet-PhlF | **pTet* template**: -76 to -1; -35: -35 to -30; -10: -12 to -7<br>Underlined overlap: **tetO2** -54 to -36 within pTet* template; underlined bases use the larger template-feature colour.<br>Underlined overlap: **tetO2** -29 to -11 within pTet* template; underlined bases use the larger template-feature colour. | **PhlO**: +1 to +30 | ttttcagcaggacgcactgacctccctatcagtgatagagattgacatccctatcagtgatagagatactgagcacaccttaacgatacggtacgtttcgtatcat |
| pTet-Ttg | **pTet* template**: -76 to -1; -35: -35 to -30; -10: -12 to -7<br>Underlined overlap: **tetO2** -54 to -36 within pTet* template; underlined bases use the larger template-feature colour.<br>Underlined overlap: **tetO2** -29 to -11 within pTet* template; underlined bases use the larger template-feature colour. | **TtgO**: +1 to +30 | ttttcagcaggacgcactgacctccctatcagtgatagagattgacatccctatcagtgatagagatactgagcacATACTTACATTCATGGTTGTTTGTAAATAC |
| pTet-Van | **pTet* template**: -76 to -1; -35: -35 to -30; -10: -12 to -7<br>Underlined overlap: **tetO2** -54 to -36 within pTet* template; underlined bases use the larger template-feature colour.<br>Underlined overlap: **tetO2** -29 to -11 within pTet* template; underlined bases use the larger template-feature colour. | **VanO2**: +1 to +30 | ttttcagcaggacgcactgacctccctatcagtgatagagattgacatccctatcagtgatagagatactgagcacATTGGATCCAATagcgctATTGGATCCAAT |
| pBet-Tac | **pBetI template**: -63 to -1; -35: -35 to -30; -10: -12 to -7<br>Underlined overlap: **BetO** -29 to -8 within pBetI template; underlined bases use the larger template-feature colour. | **lacOsym**: +1 to +20 | agcgcgggtgagagggattcgttaccaatagacaattgattggacgttcaatataatgctagcaattgtgagcgctcacaatt |
| pPhlF-Tac | **pPhlF template**: -66 to -1; -35: -35 to -31; -10: -12 to -7<br>Underlined overlap: **native PhlO** -30 to -1 within pPhlF template; underlined bases use the larger template-feature colour. | **lacOsym**: +1 to +20 | cgacgtacggtggaatctgattcgttaccaattgacatgatacgaaacgtaccgtatcgttaaggtaattgtgagcgctcacaatt |
| pTtg-Tac | **pTtg template**: -48 to +3; -35: -36 to -31; -10: -10 to -1; +1: +1<br>Underlined overlap: **native TtgO** -30 to -11 within pTtg template; underlined bases use the larger template-feature colour. | **lacOsym**: +4 to +23 | cacccagcagtatttacaaacaaccatgaatgtaagtatattccttagcaaaattgtgagcgctcacaatt |

| | | | |
|---|---|---|---|
| pVan-Tac | **pVanCC template**: -46 to +4; -35: -35 to -30; -10: -12 to -7; +1: +1<br>Underlined overlap: **VanO2 half-site** -46 to -36 within pVanCC template; underlined bases use the larger template-feature colour.<br>Underlined overlap: **VanO2 half-site** -8 to +3 within pVanCC template; underlined bases use the larger template-feature colour. | **lacOsym**: +5 to +24 | attggatccaattgacagctagctcagtcctaggtaccattggatccaataattgtgagcgctcacaatt |
| pVan-Ttg | **pVanCC template**: -46 to +4; -35: -35 to -30; -10: -12 to -7; +1: +1<br>Underlined overlap: **VanO2 half-site** -46 to -36 within pVanCC template; underlined bases use the larger template-feature colour.<br>Underlined overlap: **VanO2 half-site** -8 to +3 within pVanCC template; underlined bases use the larger template-feature colour. | **TtgO**: +5 to +34 | attggatccaattgacagctagctcagtcctaggtaccattggatccaatATACTTACATTCATGGTTGTTTGTAAATAC |
| pTac-Tet | **pTac template**: -37 to +19; -35: -35 to -30; -10: -13 to -8; +1: +1<br>Underlined overlap: **lacOsym** -1 to +19 within pTac template; underlined bases use the larger template-feature colour. | **tetO2**: +20 to +38; **lacO (native)**: +39 to +55 | tgttgacaattaatcatcggctcgtataatgtgtggaattgtgagcgctcacaatttccctatcagtgatagagaTTGTGAGCGGATAACAA |
| pBet-Van | **pBetI template**: -63 to -1; -35: -35 to -30; -10: -12 to -7<br>Underlined overlap: **BetO** -29 to -8 within pBetI template; underlined bases use the larger template-feature colour. | **VanO2**: +1 to +30 | agcgcgggtgagagggattcgttaccaatagacaattgattggacgttcaatataatgctagcATTGGATCCAATagcgctATTGGATCCAAT |
| pPhlF-Tet | **pPhlF template**: -66 to -1; -35: -35 to -31; -10: -12 to -7<br>Underlined overlap: **native PhlO** -30 to -1 within pPhlF template; underlined bases use the larger template-feature colour. | **tetO2**: +1 to +19 | cgacgtacggtggaatctgattcgttaccaattgacatgatacgaaacgtaccgtatcgttaaggttccctatcagtgatagaga |

**Supplementary Table S7 | Compact promoter sequence annotations for engineered promoter architectures.** This consolidated table reports promoter name, annotated template -35/-10 region, added-operator position relative to +1, and full 5' to 3' sequence.

| Promoter | Reporter | Main figures | Supplementary figures |
|---|---|---|---|
| pBet-Tac | GFP | Fig. 1b; Fig. 2; Fig. 3 | Fig. S13a |
| pBet-Tac | RFP | Fig. 1b; Fig. 2; Fig. 3; Fig. 4; Fig. 5 | Fig. S13b; Fig. S2 |
| pBet-Van | GFP | Fig. 1b; Fig. 2; Fig. 3; Fig. 4; Fig. 5 | Fig. S13c; Fig. S3 |
| pBet-Van | RFP | Fig. 1b; Fig. 2; Fig. 3 | Fig. S13d |
| pPhlF-Tac | GFP | Fig. 1b; Fig. 2; Fig. 3 | Fig. S14a |
| pPhlF-Tac | RFP | Fig. 1b; Fig. 2; Fig. 3; Fig. 4; Fig. 5 | Fig. S14b; Fig. S4 |
| pPhlF-Tet | RFP | Fig. 1b; Fig. 2; Fig. 3; Fig. 4; Fig. 5 | Fig. S14c; Fig. S5 |
| pTac-Tet | GFP | Fig. 1b; Fig. 2; Fig. 3; Fig. 5 | Fig. S14d; Fig. S6 |
| pTet-Bet | GFP | Fig. 1b; Fig. 2; Fig. 3 | Fig. S15a |
| pTet-Bet | RFP | Fig. 1b; Fig. 2; Fig. 3; Fig. 4; Fig. 5 | Fig. S15b; Fig. S7 |
| pTet-PhlF | RFP | Fig. 1b; Fig. 2; Fig. 3; Fig. 4; Fig. 5 | Fig. S15c; Fig. S1 |
| pTet-Ttg | GFP | Fig. 1b; Fig. 2; Fig. 3 | Fig. S15d |

| | | | |
|---|---|---|---|
| pTet-Ttg | RFP | Fig. 1b; Fig. 2; Fig. 3; Fig. 4; Fig. 5 | Fig. S16a; Fig. S8 |
| pTet-Van | GFP | Fig. 1b; Fig. 2; Fig. 3 | Fig. S16b |
| pTet-Van | RFP | Fig. 1b; Fig. 2; Fig. 3; Fig. 4; Fig. 5 | Fig. S16c; Fig. S9 |
| pTtg-Tac | GFP | Fig. 1b; Fig. 2; Fig. 3 | Fig. S16d |
| pTtg-Tac | RFP | Fig. 1b; Fig. 2; Fig. 3; Fig. 4; Fig. 5 | Fig. S16e; Fig. S10 |
| pVan-Tac | RFP | Fig. 1b; Fig. 2; Fig. 3; Fig. 5 | Fig. S16f; Fig. S11 |
| pVan-Ttg | RFP | Fig. 1b; Fig. 2; Fig. 3; Fig. 5 | Fig. S16g; Fig. S12 |
| pTac-Van | GFP | Fig. 5 | Fig. S17 |
| pTac-Van | RFP | not shown | Fig. S17 |

**Supplementary Table S8 | Cross-reference between characterised promoter/reporter constructs and the current manuscript and supplementary figure panels.** Supplementary Table S2 contains all 19 curated promoter-reporter truth tables; Supplementary Table S12 contains the template-promoter conditional activation-fold control analysis.

```
cggcctttttacggttcctggccttttgctggccttttgctcacatgttctttcctgcgttatcccctgattctgtggat
aaccgtattaccgcctttgagtgagctgataccgctcgccgcagccgaacgaccgagcgcagcgagtaagaggacatccg
gtcaaataaaacgaaaggctcagtcgaaagactgggcctttcgttttagacTTCAGCCAAAAAACTTAAGACCGCCGGTC
TTGTCCACTACCTTGCAGTAATGCGGTGGACAGGATCGGCGGTTTTCTTTTCTCTTCTCAAAATGCCaGAGACCcggaaG
GCGCGCCaagattGGTCTCaCCGCAAATCGCGGCCTTTTTTATTGATAACAAAAAGGCGCTACTACTAAACGCCAGAGTa
gcgcgggtgagagggattcgttaccaatagacaattgattggacgttcaatataatgctagcaattgtgagcgctcacaa
ttAAGATCGGGTGTAAACagctgtcaccggatgtgctttccggtctgatgagtccgtgaggacgaaacagcctctacaaa
taattttgtttaaTgatAGAGAAAGAGGGGAAAgtaTAGatggtgagcaagggcgaggaggataacatggccatcatcaa
ggagttcatgcgcttcaaggtgcacatggagggctccgtgaacggccacgagttcgagatcgagggcgagggcgagggcc
gcccctacgagggcacccagaccgccaagctgaaggtgaccaagggtggccccctgcccttcgcctgggacatcctgtcc
cctcagttcatgtacggctccaaggcctacgtgaagcaccccgccgacatccccgactacttgaagctgtccttccccga
gggcttcaagtgggagcgcgtgatgaacttcgaggacggcggcgtggtgaccgtgacccaggactcctccttgcaggacg
gcgagttcatctacaaggtgaagctgcgcggcaccaacttcccctccgacggccccgtaatgcagaagaagaccatgggc
tgggaggcctcctccgagcggatgtaccccgaggacggcgccctgaagggcgagatcaagcagaggctgaagctgaagga
cggcggccactacgacgctgaggtcaagaccacctacaaggccaagaagcccgtgcagctgcccggcgcctacaacgtca
acatcaagttggacatcacctcccacaacgaggactacaccatcgtggaacagtacgaacgcgccgagggccgccactcc
accggcggcatggacgagctgtacaagGGCTCGGGATAATAACCTAGAGAAAGAGGGGAAATACTAGATGgagaaaaaaa
tcactggatataccaccgttgatatatcccaatggcatcgtaaagaacattttgaggcatttcagtcagttgctcaatgt
acctataaccagaccgttcagctggatattacggcctttttaaagaccgtaaagaaaaataagcacaagttttatccggc
ctttattcacattcttgcccgcctgatgaatgctcatccggagttccgtatggcaatgaaagacggtgagctggtgatat
gggatagtgttcacccttgttacaccgttttccatgagcaaactgaaacgttttcatcgctctggagtgaataccacgac
gatttccggcagtttctacacatatattcgcaagatgtggcgtgttacggtgaaaacctggcctatttccctaaagggtt
tattgagaatatgtttttcgtctcagccaatccctgggtgagtttcaccagttttgatttaaacgtggccaatatggaca
acttcttcgcccccgttttcactatgggcaaatattatacgcaaggcgacaaggtgctgatgccgctggcgattcaggtt
catcatgccgtttgtgatggcttccatgtcggcagaatgcttaatgaattacaacagtactgcgatgagtggcagggcgg
ggcgggcagcggttctggctccagccatattcaacgggaaacgtcttgctcgaggccgcgattaaattccaacatggatg
ctgatttatatgggtataaatgggctcgcgataatgtcgggcaatcaggtgcgacaatctatcgattgtatgggaagccc
gatgcgccagagttgtttctgaaacatggcaaaggtagcgttgccaatgatgttacagatgagatggtcagactaaactg
gctgacggaatttatgcctcttccgaccatcaagcattttatccgtactcctgatgatgcatggttactcaccactgcga
tccccgggaaaacagcattccaggtattagaagaatatcctgattcaggtgaaaatattgttgatgcgctggcagtgttc
ctgcgccggttgcattcgattcctgtttgtaattgtccttttaacagcgatcgcgtatttcgtctcgctcaggcgcaatc
acgaatgaataacggtttggttgatgcgagtgattttgatgacgagcgtaatggctggcctgttgaacaagtctggaaag
aaatgcataagcttttgccattctcaccggattcagtcgtcactcatggtgatttctcacttgataaccttatttttgac
gaggggaaattaataggttgtattgctgttggacgagtcggaatcgcagaccgataccaggatcttgccatcctatggaa
ctgcctcggtgagttttctccttcattacagaaacggctttttcaaaaatatggtattgataatcctgatatgaataaat
tgcagtttcatttgatgctcgatgagtttttctaactgtcagaccaagtttactcatatatactttagattgatttaaaa
cttcatttttaatttcttggactcctgttgatagatccagtaatgacctcagaactccatctggatttgttcagaacgct
cggttgccgccgggcgttttttattggtgagaatccaggggtccccaataattacgatttaaattagtagcccgcctaat
gagcgggcttttttttaattcccctatttgtttatttttctaaatacattcaaatatgtatccgctcatgagacaataac
cctgataaatgcttcaataatattgaaaaaggaagagtatgagcattcagcattttcgtgtggcgctgattccgtttttt
gcggcgttttgcctgccggtgtttgcgcatccggaaaccctggtgaaagtgaaagatgcggaagatcaactgggtgcgcg
cgtgggctatattgaactggatctgaacagcggcaaaattctggaatcttttcgtccggaagaacgttttccgatgatga
gcacctttaaagtgctgctgtgcggtgcggttctgagccgtgtggatgcgggccaggaacaactgggccgtcgtattcat
tatagccagaacgatctggtggaatatagcccggtgaccgaaaaacatctgaccgatggcatgaccgtgcgtgaactgtg
cagcgcggcgattaccatgagcgataacaccgcggcgaacctgctgctgacgaccattggcggtccgaaagaactgaccg
cgtttctgcataacatgggcgatcatgtgacccgtctggatcgttgggaaccggaactgaacgaagcgattccgaacgat
gaacgtgataccaccatgccggcagcaatggcgaccaccctgcgtaaactgctgacgggtgagctgctgaccctggcaag
ccgccagcaactgattgattggatggaagcggataaagtggcgggtccgctgctgcgtagcgcgctgccggctggctggt
ttattgcggataaaagcggtgcgggcgaacgtggcagccgtggcattattgcggcgctgggcccggatggtaaaccgagc
cgtattgtggtgatttataccaccggcagccaggcgacgatggatgaacgtaaccgtcagattgcggaaattggcgcgag
cctgattaaacattggtaaccgatacaattaaaggctccttttggagcctttttttttggaaaaaggatctaggtgaag
atcctttttgataatctcatgaccaaaatcccttaacgtgagttttcgttccactgagcgtcagaccccgtagaaaagat
caaaggatcttcttgagatcctttttttCctgcgcgtaatctgctgcttgcaaacaaaaaaaccaccgctaccagcggtgg
tttgtttgccggatcaagagctaccaactctttttccgaaggtaactggcttcagcagagcgcagataccaaatactgtC
cttctagtgtagccgtagttaggccaccacttcaagaactctgtagcaccgcctacatacctcgctctgctaatcctgtt
accagtggctgctgccagtggcgataagtcgtgtcttaccgggttggactcaagacgatagttaccggataaggcgcagc
ggtcgggctgaacggggggttcgtgcacacagcccagcttggagcgaacgacctacaccgaactgagatacctacagcgt
gagctatgagaaagcgccacgcttcccgaagggagaaaggcggacaggtatccggtaagcggcagggtcggaacaggaga
gcgcacgagggagcttccagggggaaacgcctggtatctttatagtcctgtcgggtttcgccacctctgacttgagcgtc
gatttttgtgatgctcgtcaggggggcggagcctatggaaaaacgccagcaacg
```

**Supplementary Table S9 | Representative mCherry plasmid sequence.** The complete plasmid sequence is shown as a single continuous entry, with one consistent colour assigned to each annotated feature across all of its segments. Feature key: rrnB T1 terminator; spy strong terminator; 18a-AAN-ONE gBlock; L3S1P22 strong terminator; gBlock 8xgRNA; pBetI promoter (Cho); lac operator (symmetric); RiboJ ribozyme insulator; RBS-64; mCherry; gBlock AAN-mCherry-R4-TWO-B; cat (CmR) CDS chloramphenicol resistance protein; 3GS linker; dKanR; TAMPC cassette from SEVA191; lambda t0 terminator; AmpR promoter; AmpR; fd terminator; ColE1 promoter; RNAII ColE1; ColE1 P1 RNA I promoter; pMB1 mutated to medium copy number.

```
cggcctttttacggttcctggccttttgctggccttttgctcacatgttctttcctgcgttatcccctgattctgtgg
ataaccgtattaccgcctttgagtgagctgataccgctcgccgcagccgaacgaccgagcgcagcgagtaagaggaca
tccggtcaaataaaacgaaaggctcagtcgaaagactgggcctttcgttttagacTTCAGCCAAAAAACTTAAGACCG
CCGGTCTTGTCCACTACCTTGCAGTAATGCGGTGGACAGGATCGGCGGTTTTCTTTTCTCTTCTCAAAATGCCaGAGA
CCcggaaGGCGCGCCaagattGGTCTCaCCGCAAATCGCGGCCTTTTTTATTGATAACAAAAAGGCGCTACTACTAAA
CGCCAGAGTagcgcgggtgagagggattcgttaccaatagacaattgattggacgttcaatataatgctagcaattgt
gagcgctcacaattAAGATCGGGTGTAAACagctgtcaccggatgtgctttccggtctgatgagtccgtgaggacgaa
acagcctctacaaataattttgtttaaTgatAGAGAAAGAGGGGAAAgtaTAGatggtgagcaagggcgaggagctgt
tcaccggggtggtgcccatcctggtcgagctggacggcgacgtaaacggccacaagttcagcgtgtccggcgagggcg
agggcgatgccacctacggcaagctgaccctgaagttcatctgcaccaccggcaagctgcccgtgccctggcccaccc
tcgtgaccaccctgacctacggcgtgcagtgcttcagccgctaccccgaccacatgaagcagcacgacttcttcaagt
ccgccatgcccgaaggctacgtccaggagcgcaccatcttcttcaaggacgacggcaactacaagacccgcgccgagg
tgaagttcgagggcgacaccctggtgaaccgcatcgagctgaagggcatcgacttcaaggaggacggcaacatcctgg
ggcacaagctggagtacaactacaacagccacaacgtctatatcatggccgacaagcagaagaacggcatcaaggtga
acttcaagatccgccacaacatcgaggacggcagcgtgcagctcgccgaccactaccagcagaacacccccatcggcg
acggccccgtgctgctgcccgacaaccactacctgagcacccagtccgccctgagcaaagaccccaacgagaagcgcg
atcacatggtcctActggagttcgtgaccgccgccgggatcactctcggcatggacgagctgtacaagTAATAACCTA
GAGAAAGAGGGGAAATACTAGATGgagaaaaaaatcactggatataccaccgttgatatatcccaatggcatcgtaaa
gaacattttgaggcatttcagtcagttactcaatgtacctataaccagaccgttcagctggatattacggcctttta
aagaccgtaaagaaaaataagcacaagttttatccggcctttattcacattcttgcccgcctgatgaatgctcatccg
gagttccgtatggcaatgaaagacggtgagctggtgatatgggatagtgttcacccttgttacaccgttttccatgag
caaactgaaacgttttcatcgctctggagtgaataccacgacgatttccggcagtttctacacatatattcgcaagat
gtggcgtgttacggtgaaaacctggcctatttccctaaagggtttattgagaatatgtttttcgtctcagccaatccc
tgggtgagtttcaccagttttgatttaaacgtggccaatatggacaacttcttcgcccccgttttcactatgggcaaa
tattatacgcaaggcgacaaggtgctgatgccgctggcgattcaggttcatgctgccgtttgtgatggcttccatgtc
ggcagaatgcttaatgaattacaacagtactgcgatgagtggcagggcggggcgggcagcggttctggctccagccat
attcaacgggaaacgtcttgctctaggccgcgattaaattccaacatggatgctgatttatatgggtataaatgggct
cgcgataatgtcgggcaatcaggtgcgacaatctatcgattgtatgggaagcccgatgcgccagagttgtttctgaaa
catggcaaaggtagcgttgccaatgatgttacagatgagatggtcagactaaactggctgacggaatttatgcctctt
ccgaccatcaagcattttatccgtactcctgatgatgcatggttactcaccactgcgatccccgggaaaacagcattc
caggtattagaagaatatcctgattcaggtgaaaatattgttgatgcgctggcagtgttcctgcgccggttgcattcg
attcctgtttgtaattgtccttttaacagcgaccgcgtatttcgtctcgctcaggcgcaatcacgaatgaataacggt
ttggttgatgcgagtgattttgatgacgagcgtaatggctggcctgttgaacaagtctggaaagaaatgcataaactt
ttgccattctcaccggattcagtcgtcactcatggtgatttctcacttgataaccttatttttgacgaggggaaatta
ataggttgtattgatgttggacgagtcggaatcgcagaccgataccaggatcttgccatcctatggaactgcctcggt
gagttttctccttcattacagaaacggctttttcaaaaatatggtattgataatcctgatatgaataaattgcagttt
catttgatgctcgatgagtttttctaactgtcagaccaagtttactcatatatactttagattgatttaaaacttcat
ttttaatttcttggactcctgttgatagatccagtaatgacctcagaactccatctggatttgttcagaacgctcggt
tgccgccgggcgttttttattggtgagaatccaggggtccccaataattacgatttaaattagtagcccgcctaatga
gcgggcttttttttaattcccctatttgtttatttttctaaatacattcaaatatgtatccgctcatgagacaataac
cctgataaatgcttcaataatattgaaaaaggaagagtatgagcattcagcatttcgtgtggcgctgattccgtttt
ttgcggcgttttgcctgccggtgtttgcgcatccggaaaccctggtgaaagtgaaagatgcggaagatcaactgggtg
cgcgcgtgggctatattgaactggatctgaacagcggcaaaattctggaatcttttcgtccggaagaacgttttccga
tgatgagcacctttaaagtgctgctgtgcggtgcggttctgagccgtgtggatgcgggccaggaacaactgggccgtc
gtattcattatagccagaacgatctggtggaatatagcccggtgaccgaaaaacatctgaccgatggcatgaccgtgc
gtgaactgtgcagcgcggcgattaccatgagcgataacaccgcggcgaacctgctgctgacgaccattggcggtccga
aagaactgaccgcgtttctgcataacatgggcgatcatgtgacccgtctggatcgttgggaaccggaactgaacgaag
cgattccgaacgatgaacgtgataccaccatgccggcagcaatggcgaccaccctgcgtaaactgctgacgggtgagc
tgctgaccctggcaagccgccagcaactgattgattggatggaagcggataaagtggcgggtccgctgctgcgtagcg
cgctgccggctggctggtttattgcggataaaagcggtgcgggcgaacgtggcagccgtggcattattgcggcgctgg
gcccggatggtaaaccgagccgtattgtggtgatttataccaccggcagccaggcgacgatggatgaacgtaaccgtc
agattgcggaaattggcgcgagcctgattaaacattggtaaaccgatacaattaaaggctcctttggagcctttttt
tttggaaaaaggatctaggtgaagatcctttttgataatctcatgaccaaaatcccttaacgtgagttttcgttccac
tgagcgtcagaccccgtagaaaagatcaaaggatcttcttgagatcctttttttCctgcgcgtaatctgctgcttgcaa
acaaaaaaaccaccgctaccagcggtggtttgtttgccggatcaagagctaccaactctttttccgaaggtaactggc
ttcagcagagcgcagataccaaatactgtCcttctagtgtagccgtagttaggccaccacttcaagaactctgtagca
ccgcctacatacctcgctctgctaatcctgttaccagtggctgctgccagtggcgataagtcgtgtcttaccgggttg
gactcaagacgatagttaccggataaggcgcagcggtcgggctgaacggggggttcgtgcacacagcccagcttggag
cgaacgacctacaccgaactgagatacctacagcgtgagctatgagaaagcgccacgcttcccgaagggagaaaggcg
gacaggtatccggtaagcggcagggtcggaacaggagagcgcacgagggagcttccagggggaaacgcctggtatctt
tatagtcctgtcgggtttcgccacctctgacttgagcgtcgatttttgtgatgctcgtcaggggggcggagcctatgg
aaaaacgccagcaacg
```

**Supplementary Table S10 | Representative EGFP plasmid sequence.** The complete plasmid sequence is shown as a single continuous entry, with one consistent colour assigned to each annotated feature across all of its segments. Feature key: rrnB T1 terminator; spy strong terminator; 18a-AAN-ONE gBlock; L3S1P22 strong terminator; gBlock 8xgRNA; pBetI promoter (Cho); lac operator (symmetric); RiboJ ribozyme insulator; RBS-64; EGFP; dCmR (chloramphenicol resistance knockout); 3GS linker; KanR; TAMPC cassette from SEVA191; lambda t0 terminator; AmpR promoter; AmpR; fd terminator; ColE1 promoter; RNAII ColE1; ColE1 P1 RNA I promoter; pMB1 mutated to medium copy number.

| Architecture | Template | Added input | Added operator | Orientation | Approximate added-site position | Role | Selected reporter | Selected call |
|---|---|---|---|---|---|---|---|---|
| pTet-PhlF | pTet* | PhlF | PhlO | Reverse | Downstream (+1 to +30) | Curated | RFP | Pass |
| pBet-Tac | pBetI | Tac | lacOsym | Forward | Downstream (+1 to +20) | Curated | RFP | Pass |
| pBet-Van | pBetI | Van | VanO2 | Forward | Downstream (+1 to +30) | Curated | GFP | Pass |
| pPhlF-Tac | pPhlF | Tac | lacOsym | Forward | Downstream (+1 to +20) | Curated | RFP | Pass |
| pPhlF-Tet | pPhlF | Tet | tetO2 | Forward | Downstream (+1 to +19) | Curated | RFP | Pass |
| pTac-Tet | pTac | Tet | tetO2 | Forward | Distal downstream (+20 to +38) | Curated | GFP | Fail |
| pTet-Bet | pTet* | Bet | BetO | Forward | Downstream (+1 to +22) | Curated | RFP | Pass |
| pTet-Ttg | pTet* | Ttg | TtgO | Reverse | Downstream (+1 to +30) | Curated | RFP | Pass |
| pTet-Van | pTet* | Van | VanO2 | Forward | Downstream (+1 to +30) | Curated | RFP | Pass |
| pTtg-Tac | pTtg | Tac | lacOsym | Forward | Downstream (+4 to +23) | Curated | RFP | Pass |
| pVan-Tac | pVanCC | Tac | lacOsym | Forward | Downstream (+5 to +24) | Curated | RFP | N.C. |
| pVan-Ttg | pVanCC | Ttg | TtgO | Reverse | Downstream (+5 to +34) | Curated | RFP | Fail |
| pTac-Van | pTac | Van | VanO2 | Forward | Exploratory downstream insertion | Exploratory | Incomplete non-toxic truth table | Not included in curated comparison |

**Supplementary Table S11 | Overview of the engineered promoter architectures.** This table consolidates the scaffold, added operator, orientation, approximate downstream position, curated versus exploratory status, and selected reporter call for the promoter architectures discussed across Supplementary Table S7.

a

| Template | AND promoter | Role | Template inducer | Added inducer | AND reporter | Template reporter | Requested conc. | Used conc. | Match | Distance |
|---|---|---|---|---|---|---|---|---|---|---|
| pBetI | pBet-Tac | curated | Cho | IPTG | GFP | GFP | 10000 | 10000 | exact | 0.000 |
| pBetI | pBet-Tac | curated | Cho | IPTG | RFP | GFP | 10000 | 10000 | exact | 0.000 |
| pBetI | pBet-Van | curated | Cho | Van | GFP | GFP | 10000 | 10000 | exact | 0.000 |
| pBetI | pBet-Van | curated | Cho | Van | RFP | GFP | 10000 | 10000 | exact | 0.000 |
| pPhlF | pPhlF-Tac | curated | DAPG | IPTG | GFP | GFP | 10.00 | 10.00 | exact | 0.000 |
| pPhlF | pPhlF-Tac | curated | DAPG | IPTG | RFP | RFP | 10.00 | 2.100 | nearest_available | 7.900 |
| pPhlF | pPhlF-Tet | curated | DAPG | aTc | RFP | RFP | 10.00 | 2.100 | nearest_available | 7.900 |
| pTac | pTac-Tet | curated | IPTG | aTc | GFP | GFP | 1000 | 500.00 | nearest_available | 500.00 |
| pTet* | pTet-Bet | curated | aTc | Cho | GFP | GFP | 100.00 | 100.00 | exact | 0.000 |
| pTet* | pTet-Bet | curated | aTc | Cho | RFP | RFP | 0.200 | 0.200 | exact | 0.000 |
| pTet* | pTet-PhlF | curated | aTc | DAPG | RFP | RFP | 0.200 | 0.200 | exact | 0.000 |
| pTet* | pTet-Ttg | curated | aTc | Nar | GFP | GFP | 100.00 | 100.00 | exact | 0.000 |
| pTet* | pTet-Ttg | curated | aTc | Nar | RFP | RFP | 100.00 | 0.200 | nearest_available | 99.80 |
| pTet* | pTet-Van | curated | aTc | Van | GFP | GFP | 0.200 | 0.200 | exact | 0.000 |
| pTet* | pTet-Van | curated | aTc | Van | RFP | RFP | 0.200 | 0.200 | exact | 0.000 |
| pTtg | pTtg-Tac | curated | Nar | IPTG | GFP | GFP | 280.00 | 280.00 | exact | 0.000 |
| pTtg | pTtg-Tac | curated | Nar | IPTG | RFP | RFP | 280.00 | 280.00 | exact | 0.000 |
| pVanCC | pVan-Tac | curated | Van | IPTG | RFP | RFP | 21.00 | 100.00 | nearest_available | 79.00 |
| pVanCC | pVan-Ttg | curated | Van | Nar | RFP | RFP | 100.00 | 100.00 | exact | 0.000 |
| pTac | pTac-Van | exploratory | IPTG | Van | GFP | GFP | 1000 | 500.00 | nearest_available | 500.00 |
| pTac | pTac-Van | exploratory | IPTG | Van | RFP | RFP | 1000 | 500.00 | nearest_available | 500.00 |

b

| Template | AND promoter | Reporter | Template [0] mean | Template [1] mean | Template [0] sd | Template [1] sd | n [0] | n [1] | R_YES | log2 R_YES | CI low | CI high |
|---|---|---|---|---|---|---|---|---|---|---|---|---|
| pBetI | pBet-Tac | GFP | 7396 | 49143 | 0.000 | 4601 | 1 | 8 | 6.644 | 2.732 | 2.648 | 2.820 |
| pBetI | pBet-Tac | RFP | 7396 | 49143 | 0.000 | 4601 | 1 | 8 | 6.644 | 2.732 | 2.645 | 2.821 |
| pBetI | pBet-Van | GFP | 7396 | 49143 | 0.000 | 4601 | 1 | 8 | 6.644 | 2.732 | 2.649 | 2.820 |
| pBetI | pBet-Van | RFP | 7396 | 49143 | 0.000 | 4601 | 1 | 8 | 6.644 | 2.732 | 2.648 | 2.820 |
| pPhlF | pPhlF-Tac | GFP | 62615 | 64180 | 2145 | 282.10 | 4 | 2 | 1.025 | 0.036 | -0.006 | 0.080 |
| pPhlF | pPhlF-Tac | RFP | 13678 | 54997 | 276.57 | 4023 | 3 | 3 | 4.021 | 2.008 | 1.900 | 2.106 |
| pPhlF | pPhlF-Tet | RFP | 13678 | 54997 | 276.57 | 4023 | 3 | 3 | 4.021 | 2.008 | 1.900 | 2.106 |
| pTac | pTac-Tet | GFP | 6611 | 76500 | 345.73 | 2528 | 6 | 6 | 11.57 | 3.533 | 3.467 | 3.598 |
| pTet* | pTet-Bet | GFP | 6578 | 70382 | 153.50 | 0.000 | 3 | 1 | 10.70 | 3.420 | 3.394 | 3.458 |
| pTet* | pTet-Bet | RFP | 0.000 | 34173 | 0.000 | 5038 | 3 | 12 | 1667 | 10.70 | 10.58 | 10.81 |
| pTet* | pTet-PhlF | RFP | 0.000 | 34173 | 0.000 | 5038 | 3 | 12 | 1667 | 10.70 | 10.58 | 10.81 |

| | | | | | | | | | | | | |
|---|---|---|---|---|---|---|---|---|---|---|---|---|
| pTet* | pTet-Ttg | GFP | 6578 | 70382 | 153.50 | 0.000 | 3 | 1 | 10.70 | 3.420 | 3.394 | 3.458 |
| pTet* | pTet-Ttg | RFP | 0.000 | 34173 | 0.000 | 5038 | 3 | 12 | 1667 | 10.70 | 10.58 | 10.81 |
| pTet* | pTet-Van | GFP | 6578 | 56517 | 153.50 | 3215 | 3 | 18 | 8.592 | 3.103 | 3.055 | 3.152 |
| pTet* | pTet-Van | RFP | 0.000 | 34173 | 0.000 | 5038 | 3 | 12 | 1667 | 10.70 | 10.58 | 10.81 |
| pTtg | pTtg-Tac | GFP | 8463 | 43899 | 666.68 | 2782 | 3 | 13 | 5.187 | 2.375 | 2.267 | 2.494 |
| pTtg | pTtg-Tac | RFP | 31.24 | 7322 | 0.000 | 338.84 | 1 | 12 | 234.39 | 7.873 | 7.835 | 7.908 |
| pVanCC | pVan-Tac | RFP | 0.000 | 55931 | 0.000 | 4339 | 6 | 7 | 2729 | 11.41 | 11.34 | 11.49 |
| pVanCC | pVan-Ttg | RFP | 0.000 | 55931 | 0.000 | 4339 | 6 | 7 | 2729 | 11.41 | 11.34 | 11.49 |
| pTac | pTac-Van | GFP | 6611 | 76500 | 345.73 | 2528 | 6 | 6 | 11.57 | 3.533 | 3.468 | 3.597 |
| pTac | pTac-Van | RFP | 0.000 | 32917 | 0.000 | 25551 | 3 | 17 | 1606 | 10.65 | 9.975 | 11.08 |

**c**

| Template | AND promoter | Reporter | AND [01] mean | AND [11] mean | AND [01] sd | AND [11] sd | n [01] | n [11] | R_AND | log2 R_AND | CI low | CI high |
|---|---|---|---|---|---|---|---|---|---|---|---|---|
| pBetI | pBet-Tac | GFP | 6206 | 46516 | 141.36 | 1651 | 3 | 3 | 7.495 | 2.906 | 2.851 | 2.960 |
| pBetI | pBet-Tac | RFP | 0.000 | 6139 | 0.000 | 143.08 | 3 | 3 | 299.55 | 8.227 | 8.193 | 8.260 |
| pBetI | pBet-Van | GFP | 5741 | 59190 | 217.03 | 1099 | 3 | 3 | 10.31 | 3.366 | 3.313 | 3.420 |
| pBetI | pBet-Van | RFP | 0.000 | 0.000 | 0.000 | 0.000 | 3 | 3 | 0.000 | | | |
| pPhlF | pPhlF-Tac | GFP | 61486 | 71953 | 2243 | 678.67 | 3 | 3 | 1.170 | 0.227 | 0.171 | 0.268 |
| pPhlF | pPhlF-Tac | RFP | 2988 | 13564 | 228.75 | 258.88 | 3 | 3 | 4.539 | 2.182 | 2.083 | 2.294 |
| pPhlF | pPhlF-Tet | RFP | 16800 | 63107 | 1509 | 1653 | 3 | 3 | 3.756 | 1.909 | 1.797 | 2.047 |
| pTac | pTac-Tet | GFP | 38428 | 74064 | 26605 | 1406 | 3 | 3 | 1.927 | 0.947 | 0.440 | 3.255 |
| pTet* | pTet-Bet | GFP | 6455 | 26924 | 761.93 | 1297 | 2 | 2 | 4.171 | 2.060 | 1.895 | 2.234 |
| pTet* | pTet-Bet | RFP | 0.000 | 8479 | 0.000 | 204.60 | 3 | 3 | 413.74 | 8.693 | 8.656 | 8.725 |
| pTet* | pTet-PhlF | RFP | 0.000 | 38690 | 0.000 | 1190 | 3 | 3 | 1888 | 10.88 | 10.84 | 10.92 |
| pTet* | pTet-Ttg | GFP | 5806 | 7813 | 74.51 | 679.44 | 3 | 6 | 1.346 | 0.428 | 0.335 | 0.518 |
| pTet* | pTet-Ttg | RFP | 0.000 | 1740 | 0.000 | 94.78 | 3 | 6 | 84.88 | 6.407 | 6.347 | 6.460 |
| pTet* | pTet-Van | GFP | 6395 | 8216 | 99.68 | 785.88 | 3 | 3 | 1.285 | 0.361 | 0.264 | 0.509 |
| pTet* | pTet-Van | RFP | 0.000 | 28030 | 0.000 | 2023 | 3 | 3 | 1368 | 10.42 | 10.35 | 10.53 |
| pTtg | pTtg-Tac | GFP | 6389 | 10002 | 129.11 | 754.13 | 3 | 3 | 1.566 | 0.647 | 0.563 | 0.763 |
| pTtg | pTtg-Tac | RFP | 0.000 | 739.87 | 0.000 | 15.17 | 3 | 3 | 36.10 | 5.174 | 5.145 | 5.204 |
| pVanCC | pVan-Tac | RFP | 0.000 | 10256 | 0.000 | 0.000 | 4 | 1 | 500.44 | 8.967 | 8.967 | 8.967 |
| pVanCC | pVan-Ttg | RFP | 0.000 | 36976 | 0.000 | 236.54 | 3 | 3 | 1804 | 10.82 | 10.81 | 10.83 |
| pTac | pTac-Van | GFP | 4907 | 7140 | 243.82 | 902.17 | 3 | 3 | 1.455 | 0.541 | 0.377 | 0.719 |
| pTac | pTac-Van | RFP | 0.000 | 37664 | 0.000 | 1410 | 3 | 3 | 1838 | 10.84 | 10.79 | 10.89 |

**d**

| Template | AND promoter | Reporter | Delta log2 | Delta CI low | Delta CI high | Ratio of ratios | Denominator nonpositive | Lower bound | Response floor |
|---|---|---|---|---|---|---|---|---|---|
| pBetI | pBet-Tac | GFP | 0.174 | 0.069 | 0.277 | 1.128 | No | No | |
| pBetI | pBet-Tac | RFP | 5.495 | 5.401 | 5.587 | 45.08 | Yes | Yes | 20.49 |
| pBetI | pBet-Van | GFP | 0.634 | 0.529 | 0.734 | 1.552 | No | No | |

| | | | | | | | | | |
|---|---|---|---|---|---|---|---|---|---|
| pBetI | pBet-Van | RFP | | | | | Yes | Yes | 20.49 |
| pPhlF | pPhlF-Tac | GFP | 0.191 | 0.124 | 0.255 | 1.142 | No | No | |
| pPhlF | pPhlF-Tac | RFP | 0.175 | 0.036 | 0.325 | 1.129 | No | No | |
| pPhlF | pPhlF-Tet | RFP | -0.098 | -0.253 | 0.068 | 0.934 | No | No | |
| pTac | pTac-Tet | GFP | -2.586 | -3.121 | -0.295 | 0.167 | No | No | |
| pTet* | pTet-Bet | GFP | -1.359 | -1.525 | -1.185 | 0.390 | No | No | |
| pTet* | pTet-Bet | RFP | -2.011 | -2.121 | -1.883 | 0.248 | Yes | Yes | 20.49 |
| pTet* | pTet-PhlF | RFP | 0.179 | 0.063 | 0.307 | 1.132 | Yes | Yes | 20.49 |
| pTet* | pTet-Ttg | GFP | -2.991 | -3.091 | -2.896 | 0.126 | No | No | |
| pTet* | pTet-Ttg | RFP | -4.296 | -4.420 | -4.161 | 0.051 | Yes | Yes | 20.49 |
| pTet* | pTet-Van | GFP | -2.742 | -2.861 | -2.598 | 0.150 | No | No | |
| pTet* | pTet-Van | RFP | -0.286 | -0.427 | -0.127 | 0.820 | Yes | Yes | 20.49 |
| pTtg | pTtg-Tac | GFP | -1.728 | -1.886 | -1.575 | 0.302 | No | No | |
| pTtg | pTtg-Tac | RFP | -2.699 | -2.743 | -2.653 | 0.154 | Yes | Yes | 20.49 |
| pVanCC | pVan-Tac | RFP | -2.447 | -2.527 | -2.373 | 0.183 | Yes | Yes | 20.49 |
| pVanCC | pVan-Ttg | RFP | -0.597 | -0.676 | -0.523 | 0.661 | Yes | Yes | 20.49 |
| pTac | pTac-Van | GFP | -2.992 | -3.169 | -2.801 | 0.126 | No | No | |
| pTac | pTac-Van | RFP | 0.194 | -0.234 | 0.868 | 1.144 | Yes | Yes | 20.49 |

**Supplementary Table S12 | Template-promoter YES-gate activation ratios and AND-promoter conditional activation ratios. a,** Template and AND-promoter pair identity and concentration matching. **b,** Template-promoter YES-gate activation ratios. **c,** AND-promoter conditional activation ratios. **d,** Ratio-of-ratios, lower-bound flags and response-floor values. R_YES = template [1]/[0] and R_AND = AND [11]/[01]. The delta log2 metric is log2(R_AND) - log2(R_YES). Lower-bound rows use the empirical response floor when the denominator was non-positive.

a

| Promoter | Reporter | Channel | State | Mean | n | Alt reporter | Alt mean | Alt n | Alt/formal | Match |
|---|---|---|---|---|---|---|---|---|---|---|
| pBet-Tac | GFP | GFP | [00] | 5973 | 3 | RFP | 6472 | 3 | 1.083 | ind. |
| pBet-Tac | GFP | GFP | [10] | 6075 | 3 | RFP | 6858 | 3 | 1.129 | ind. |
| pBet-Tac | GFP | GFP | [01] | 6206 | 3 | RFP | 6398 | 3 | 1.031 | ind. |
| pBet-Tac | GFP | GFP | [11] | 46516 | 3 | RFP | 5710 | 3 | 0.123 | ind. |
| pBet-Tac | RFP | RFP | [00] | 0.000 | 3 | GFP | 0.000 | 3 | | ind. |
| pBet-Tac | RFP | RFP | [10] | 0.000 | 3 | GFP | 0.000 | 3 | | ind. |
| pBet-Tac | RFP | RFP | [01] | 0.000 | 3 | GFP | 0.000 | 3 | | ind. |
| pBet-Tac | RFP | RFP | [11] | 6139 | 3 | GFP | 0.000 | 3 | 0.000 | ind. |
| pBet-Van | GFP | GFP | [00] | 5876 | 3 | RFP | 5876 | 3 | 1.000 | same |
| pBet-Van | GFP | GFP | [10] | 6180 | 3 | RFP | 6180 | 3 | 1.000 | same |
| pBet-Van | GFP | GFP | [01] | 5741 | 3 | RFP | 5741 | 3 | 1.000 | same |
| pBet-Van | GFP | GFP | [11] | 59190 | 3 | RFP | 59190 | 3 | 1.000 | same |
| pBet-Van | RFP | RFP | [00] | 0.000 | 3 | GFP | 0.000 | 3 | | same |
| pBet-Van | RFP | RFP | [10] | 0.000 | 3 | GFP | 0.000 | 3 | | same |
| pBet-Van | RFP | RFP | [01] | 0.000 | 3 | GFP | 0.000 | 3 | | same |
| pBet-Van | RFP | RFP | [11] | 0.000 | 3 | GFP | 0.000 | 3 | | same |
| pPhlF-Tac | GFP | GFP | [00] | 6200 | 3 | RFP | 5807 | 3 | 0.937 | ind. |
| pPhlF-Tac | GFP | GFP | [10] | 6986 | 3 | RFP | 5361 | 3 | 0.767 | ind. |
| pPhlF-Tac | GFP | GFP | [01] | 61486 | 3 | RFP | 5746 | 3 | 0.093 | ind. |
| pPhlF-Tac | GFP | GFP | [11] | 71953 | 3 | RFP | 7336 | 3 | 0.102 | ind. |
| pPhlF-Tac | RFP | RFP | [00] | 0.000 | 3 | GFP | 0.000 | 3 | | ind. |
| pPhlF-Tac | RFP | RFP | [10] | 0.000 | 3 | GFP | 0.000 | 3 | | ind. |
| pPhlF-Tac | RFP | RFP | [01] | 2988 | 3 | GFP | 0.000 | 3 | 0.000 | ind. |
| pPhlF-Tac | RFP | RFP | [11] | 13564 | 3 | GFP | 0.000 | 3 | 0.000 | ind. |
| pPhlF-Tet | RFP | RFP | [00] | 0.000 | 3 | not available | | 0 | | n.a. |
| pPhlF-Tet | RFP | RFP | [10] | 0.000 | 3 | not available | | 0 | | n.a. |
| pPhlF-Tet | RFP | RFP | [01] | 16800 | 3 | not available | | 0 | | n.a. |
| pPhlF-Tet | RFP | RFP | [11] | 63107 | 3 | not available | | 0 | | n.a. |
| pTac-Tet | GFP | GFP | [00] | 7339 | 3 | not available | | 0 | | n.a. |
| pTac-Tet | GFP | GFP | [10] | 8818 | 3 | not available | | 0 | | n.a. |

| | | | | | | | | | | |
|---|---|---|---|---|---|---|---|---|---|---|
| pTac-Tet | GFP | GFP | [01] | 38428 | 3 | not available | | 0 | | n.a. |
| pTac-Tet | GFP | GFP | [11] | 74064 | 3 | not available | | 0 | | n.a. |
| pTet-Bet | GFP | GFP | [00] | 7753 | 2 | RFP | 5253 | 3 | 0.678 | ind. |
| pTet-Bet | GFP | GFP | [10] | 21701 | 1 | RFP | 5957 | 3 | 0.274 | ind. |
| pTet-Bet | GFP | GFP | [01] | 6455 | 2 | RFP | 6100 | 3 | 0.945 | ind. |
| pTet-Bet | GFP | GFP | [11] | 26924 | 2 | RFP | 6583 | 3 | 0.245 | ind. |
| pTet-Bet | RFP | RFP | [00] | 0.000 | 3 | GFP | 0.000 | 2 | | ind. |
| pTet-Bet | RFP | RFP | [10] | 0.000 | 3 | GFP | 1063 | 1 | | ind. |
| pTet-Bet | RFP | RFP | [01] | 0.000 | 3 | GFP | 0.000 | 2 | | ind. |
| pTet-Bet | RFP | RFP | [11] | 8479 | 3 | GFP | 2188 | 2 | 0.258 | ind. |
| pTet-PhlF | RFP | RFP | [00] | 0.000 | 3 | not available | | 0 | | n.a. |
| pTet-PhlF | RFP | RFP | [10] | 3517 | 3 | not available | | 0 | | n.a. |
| pTet-PhlF | RFP | RFP | [01] | 0.000 | 3 | not available | | 0 | | n.a. |
| pTet-PhlF | RFP | RFP | [11] | 38690 | 3 | not available | | 0 | | n.a. |
| pTet-Ttg | GFP | GFP | [00] | 6649 | 4 | RFP | 6649 | 4 | 1.000 | same |
| pTet-Ttg | GFP | GFP | [10] | 12377 | 2 | RFP | 12377 | 2 | 1.000 | same |
| pTet-Ttg | GFP | GFP | [01] | 5806 | 3 | RFP | 5806 | 3 | 1.000 | same |
| pTet-Ttg | GFP | GFP | [11] | 7813 | 6 | RFP | 7813 | 6 | 1.000 | same |
| pTet-Ttg | RFP | RFP | [00] | 0.000 | 4 | GFP | 0.000 | 4 | | same |
| pTet-Ttg | RFP | RFP | [10] | 0.000 | 2 | GFP | 0.000 | 2 | | same |
| pTet-Ttg | RFP | RFP | [01] | 0.000 | 3 | GFP | 0.000 | 3 | | same |
| pTet-Ttg | RFP | RFP | [11] | 1740 | 6 | GFP | 1740 | 6 | 1.000 | same |
| pTet-Van | GFP | GFP | [00] | 7451 | 3 | RFP | 7451 | 3 | 1.000 | same |
| pTet-Van | GFP | GFP | [10] | 8425 | 3 | RFP | 8425 | 3 | 1.000 | same |
| pTet-Van | GFP | GFP | [01] | 6395 | 3 | RFP | 6395 | 3 | 1.000 | same |
| pTet-Van | GFP | GFP | [11] | 8216 | 3 | RFP | 8216 | 3 | 1.000 | same |
| pTet-Van | RFP | RFP | [00] | 0.000 | 3 | GFP | 0.000 | 3 | | same |
| pTet-Van | RFP | RFP | [10] | 0.000 | 3 | GFP | 0.000 | 3 | | same |
| pTet-Van | RFP | RFP | [01] | 0.000 | 3 | GFP | 0.000 | 3 | | same |
| pTet-Van | RFP | RFP | [11] | 28030 | 3 | GFP | 28030 | 3 | 1.000 | same |
| pTtg-Tac | GFP | GFP | [00] | 5386 | 3 | RFP | 6858 | 3 | 1.273 | ind. |
| pTtg-Tac | GFP | GFP | [10] | 5547 | 3 | RFP | 6899 | 3 | 1.244 | ind. |
| pTtg-Tac | GFP | GFP | [01] | 6389 | 3 | RFP | 6304 | 3 | 0.987 | ind. |
| pTtg-Tac | GFP | GFP | [11] | 10002 | 3 | RFP | 6092 | 3 | 0.609 | ind. |
| pTtg-Tac | RFP | RFP | [00] | 0.000 | 3 | GFP | 0.000 | 3 | | ind. |

| | | | | | | | | | | |
|---|---|---|---|---|---|---|---|---|---|---|
| pTtg-Tac | RFP | RFP | [10] | 6.831 | 3 | GFP | 0.000 | 3 | 0.000 | ind. |
| pTtg-Tac | RFP | RFP | [01] | 0.000 | 3 | GFP | 0.000 | 3 | | ind. |
| pTtg-Tac | RFP | RFP | [11] | 740 | 3 | GFP | 0.000 | 3 | 0.000 | ind. |
| pVan-Tac | RFP | RFP | [00] | 0.000 | 3 | not available | | 0 | | n.a. |
| pVan-Tac | RFP | RFP | [10] | 0.000 | 3 | not available | | 0 | | n.a. |
| pVan-Tac | RFP | RFP | [01] | 0.000 | 4 | not available | | 0 | | n.a. |
| pVan-Tac | RFP | RFP | [11] | 10256 | 1 | not available | | 0 | | n.a. |
| pVan-Ttg | RFP | RFP | [00] | 0.000 | 3 | not available | | 0 | | n.a. |
| pVan-Ttg | RFP | RFP | [10] | 36787 | 3 | not available | | 0 | | n.a. |
| pVan-Ttg | RFP | RFP | [01] | 0.000 | 3 | not available | | 0 | | n.a. |
| pVan-Ttg | RFP | RFP | [11] | 36976 | 3 | not available | | 0 | | n.a. |

**b**

| Promoter | Reporter | State | Formal β1 | Alt β1 | Alt/formal | Margin check | Formal margin | Δ background | Diagnostic margin | Outcome |
|---|---|---|---|---|---|---|---|---|---|---|
| pBet-Tac | GFP | [00] | 5973 | 6472 | 1.083 | [11] vs [00] | 40542 | -762 | 41304 | No |
| pBet-Tac | GFP | [01] | 6206 | 6398 | 1.031 | [11] vs [01] | 40310 | -688 | 40998 | No |
| pBet-Tac | GFP | [10] | 6075 | 6858 | 1.129 | [11] vs [10] | 40440 | -1149 | 41589 | No |
| pBet-Tac | GFP | [11] | 46516 | 5710 | 0.123 | minimum corrected margin: [11] vs [01] | 40310 | -688 | 40998 | No |
| pPhlF-Tac | GFP | [00] | 6200 | 5807 | 0.937 | [11] vs [00] | 65753 | 1530 | 64223 | No |
| pPhlF-Tac | GFP | [01] | 61486 | 5746 | 0.093 | [11] vs [01] | 10467 | 1591 | 8876 | No |
| pPhlF-Tac | GFP | [10] | 6986 | 5361 | 0.767 | [11] vs [10] | 64966 | 1976 | 62991 | No |
| pPhlF-Tac | GFP | [11] | 71953 | 7336 | 0.102 | minimum corrected margin: [11] vs [01] | 10467 | 1591 | 8876 | No |
| pTet-Bet | GFP | [10] | 21701 | 5957 | n.a. | not assessable | n.a. | n.a. | n.a. | Not assessable |
| pVan-Tac | RFP | [11] | 10256 | n.a. | n.a. | not assessable | n.a. | n.a. | n.a. | Not assessable |

**Supplementary Table S13 | Same-channel alternate-reporter background diagnostic and margin checks.** a, State-level diagnostic for each formal reporter-specific state mean from Supplementary Table S2. Parenthetical alternate-reporter values are reported only as diagnostics. Match codes indicate an independent selected source file (ind.), the same selected source file (same; not used as an independent background estimate), or no available alternate context (n.a.). b, Exact independent margin checks for positive same-channel alternate-reporter estimates and non-assessable n = 1 cases. The diagnostic margin is the formal [11]-versus-off-state margin minus the same-channel background differential. These diagnostics were not used as normalisation controls, subtraction terms, or formal reclassification criteria.
